# Temperature-Dependent Regulation of Upstream Open Reading Frame Translation in *S. Cerevisiae*

**DOI:** 10.1101/678409

**Authors:** Shardul D. Kulkarni, Fujun Zhou, Neelam Dabas Sen, Hongen Zhang, Alan G. Hinnebusch, Jon R. Lorsch

## Abstract

**Background:** Translation of an mRNA in eukaryotes starts at AUG in most cases. Near-cognate codons (NCCs) such as UUG, ACG and AUU are also used as start sites at low levels in *S. cerevisiae*. Initiation from NCCs or AUGs in the 5’-untranslated regions (UTRs) of mRNAs can lead to translation of upstream open reading frames (uORFs) that might regulate expression of the main ORF (mORF). Although there is some circumstantial evidence that the translation of uORFs can be affected by environmental conditions, little is known about how it is affected by changes in growth temperature.

**Results:** Using reporter assays, we found that changes in growth temperature can affect translation from NCC start sites in yeast cells, suggesting the possibility that gene expression could be regulated by temperature by altering use of different uORF start codons. Using ribosome profiling, we provide evidence that growth temperature regulates the efficiency of translation of nearly 200 uORFs in *S. cerevisiae*. Of these uORFs, most that start with an AUG codon have increased translational efficiency at 37 °C relative to 30 °C and decreased efficiency at 20 °C. For translationally regulated uORFs starting with NCCs, we did not observe a general trend for the direction of regulation as a function of temperature, suggesting mRNA-specific features can determine the mode of temperature-dependent regulation. Consistent with this conclusion, the position of the uORFs in the 5’-leader relative to the 5’-cap and the start codon of the main ORF correlates with the direction of temperature-dependent regulation of uORF translation. We have identified several novel cases in which changes in uORF translation are inversely correlated with changes in the translational efficiency of the downstream main ORF. Our data suggest that translation of these mRNAs is subject to temperature-dependent, uORF-mediated regulation.

**Conclusions:** Overall, our data suggest that alterations in the translation of specific uORFs by temperature can regulate gene expression in *S. cerevisiae*.

## Background

In eukaryotes, the recognition of the start codon in an mRNA during translation initiation is thought to occur by the scanning mechanism [1]. It begins with the formation of a ternary complex (TC) that consists of translation initiation factor 2 (eIF2) in its GTP-bound form along with the methionyl initiator tRNA (Met-tRNA_i_). The TC binds to the small (40S) ribosomal subunit with the aid of eIFs 1, 1A and 3 to form the 43S pre-initiation complex (PIC). The PIC binds to the mRNA near the 5’-cap with the aid of a group of initiation factors including the eIF4F complex, eIF3 and the poly(A) binding protein (PABP). The PIC then scans along the mRNA in a 5’ to 3’ direction in search of the start codon, which in most cases is an AUG. Upon recognition of the start codon, sequential events occur in the PIC that eventually lead to joining of the large (60S) ribosomal subunit and commencement of the elongation phase of protein synthesis. This series of events starts with stable base-pairing between the start codon and tRNAi anticodon, which triggers ejection of eIF1 from its binding site on the 40S subunit. Ejection of eIF1 from the PIC leads to the conversion of eIF2 to its inactive, GDP-bound state as a result of gated phosphate (P_i_) release. This converts the open, scanning-competent PIC into a closed, scanning-arrested PIC. Release of the other initiation factors and joining of the 60S ribosomal subunit results in formation of the 80S initiation complex which is now competent for translation elongation [2, 3].

Selection of the start codon in an mRNA by the translational machinery is one of the key steps of reading the genetic code. It defines the N-terminus of the translated protein as well as the reading frame for decoding. Although AUG is the start codon of most main ORFs, codons that differ from AUG by only one base (“near-cognate codons;” NCCs) can also be utilized as start sites to varying degrees in *S. cerevisiae* [4–6]. Use of alternative start codons, such as AUGs in weak sequence context or NCCs, is a potential mechanism to regulate gene expression [7–10]. For example, in addition to their main open reading frames (mORFs), some mRNAs contain one or more upstream open reading frame (uORF) in their 5’-leaders which can begin with AUGs in strong or weak sequence context or with NCCs [11]. The recognition and translation of uORFs can regulate the expression of the downstream mORF by various mechanisms, such as altering the level of the mRNA by triggering nonsense-mediated decay or by preventing PICs from reaching the mORF start codon [12, 13]. Several studies using ribosome-profiling have provided evidence that uORF translation is altered in response to a variety of stress conditions [8, 14]. It has also been reported that some mRNAs have multiple in-frame AUGs or NCCs that can be used as alternative start sites that can lead to production of protein isoforms with N-terminal extensions. These alternative initiation codons can be conserved throughout eukaryotes, suggesting their functional importance [11]. There are some known examples of yeast mRNAs that have in-frame upstream initiation codons that lead to formation of N-terminal extensions. The protein isoforms with and without the extension have been reported to localize differentially [5, 6, 15, 16] and a recent proteomic analysis of the yeast “N-terminome” indicates that ∼10% of yeast mRNAs have alternative, in-frame start codons that are utilized some fraction of the time [17].

A number of components of the eukaryotic translation initiation machinery have been shown to be involved in start codon recognition, including initiation factors such as eIF1, eIF1A, eIF2, and eIF5, and tRNA_i_, rRNA, and mRNA elements [18]. Mutations in these components can increase the efficiency with which NCCs are used as start sites, producing a phenotype referred to as Suppressor of initiation codon mutation (Sui^-^). A mutation in eIF1A with the opposite effect on fidelity was shown to confer heightened discrimination genome-wide against AUGs in poor sequence context [19]. The use of NCC start sites in a reporter mRNA in yeast was also shown to be enhanced by to two small molecules identified in a high-throughput screen [20], indicating that external agents can modulate the fidelity of start codon recognition.

We undertook this study starting with the hypothesis that the fidelity of start codon recognition might be a point of post-transcriptional regulation of gene expression. Changes in the fidelity of start codon recognition in response to external or internal stimuli could rapidly modify the proteome by changing the balance of translation of uORFs, N-terminal extensions and main ORFs. In an attempt to test this hypothesis, we used the same dual-luciferase reporter assay used for the high-throughput chemical screen for compounds that alter start codon recognition in *S. cerevisiae* [20] to search for other external stimuli that produce similar effects. We found that growth temperature modulates the use of near-cognate start codons in both the luciferase reporter system and an orthogonal *lacZ*-based system. However, when we used ribosome profiling to observe the effects of growth temperature on translation of uORFs transcriptome-wide, we found a more complicated distribution of effects than was suggested by the reporter assays. Although translation of most uORFs is not significantly affected by changes in growth temperature, a subset of uORFs are regulated by temperature shifts, with various combinations of increased or decreased translational efficiency at high or low temperature. Of the regulated uORFs, those starting with AUG are generally repressed at 20 °C and activated at 37 °C relative to their translation at 30 °C, whereas those starting with NCCs display a more distributed set of effects. The position of the uORF in the 5’-UTR and the length and degree of structure of the UTR appear to influence the effect of temperature on translation. We present a number of novel cases of temperature-dependent changes in uORF translation in which there are reciprocal changes in main ORF translation, suggesting uORF-mediated regulation of main ORF expression.

## Materials and Methods

### Yeast strains and plasmids

The yeast cells were transformed as described previously [21] and the transformants were selected on the appropriate media lacking the nutrients corresponding to auxotrophic marker/s. The detailed list of the strains and plasmids used in the study is provided in Additional file 2: Supplementary Table 1 and 2 respectively.

### Biochemical assays

The dual luciferase assay was carried out as previously described [4, 20] with some minor modifications. The wild type/mutant yeast cells having either control reporter (R^AUG^FF^AUG^) or test reporter (R^AUG^FF^XXX^) (Figure 1A) were grown overnight to saturation. The cells were then diluted to reach a desired OD_600_ of 0.6-0.8 in 16 h at the tested temperature (e.g., 20 °C, 30 °C, etc.). To calculate the luciferase activity, 2 μl of culture was added to 50 μl of 1X Passive Lysis Buffer (Promega #E1941) which was aliquoted in a reader plate (Corning #CLS3912), followed by lysis at room temperature for 50 min. The luciferase activity was measured using a Turner Modulus Microplate Reader at 24 °C. Briefly 50 μl of F-Luc reagent (15 mM Tris pH-8.0, 25 mM glycylglycine, 4 mM EGTA, 15 mM MgSO_4_, 1 mM DTT, 2 mM ATP, 0.1 mM CoA and 75 μM luciferin) was added to each well. The activity was measured with a delay time (the duration between the injection of the reagent and taking a measurement) of 2 s and an integration time (the duration of measurement per well) of 1 second. The R-Luc activity in the same well was immediately measured by adding 50 μl of R-Luc reagent (0.22 M citric acid-sodium citrate pH 5, 1.1 M NaCl, 2.2 mM Na_2_EDTA, 1.3 mM NaN_3_, 0.44mg/ml BSA, 1.43 μM coelenterazine) with the same settings of measurement as for F-Luc. The relative activity of firefly luciferase (F-Luc) was calculated by normalizing with the activity of the Renilla luciferase (R-Luc) to yield either FF^AUG^/R^AUG^ (AUG) or FF^UUG^/R^AUG^ (UUG) values. To calculate the normalized activity of firefly luciferase starting with UUG as an initiation codon, the F^UUG^/R^AUG^ value was normalized to F^AUG^/R^AUG^ for the UUG/AUG ratio. The normalized expression of other near cognate codons (e.g., ACG, AUU) was calculated in a similar manner.

**Figure 1.**
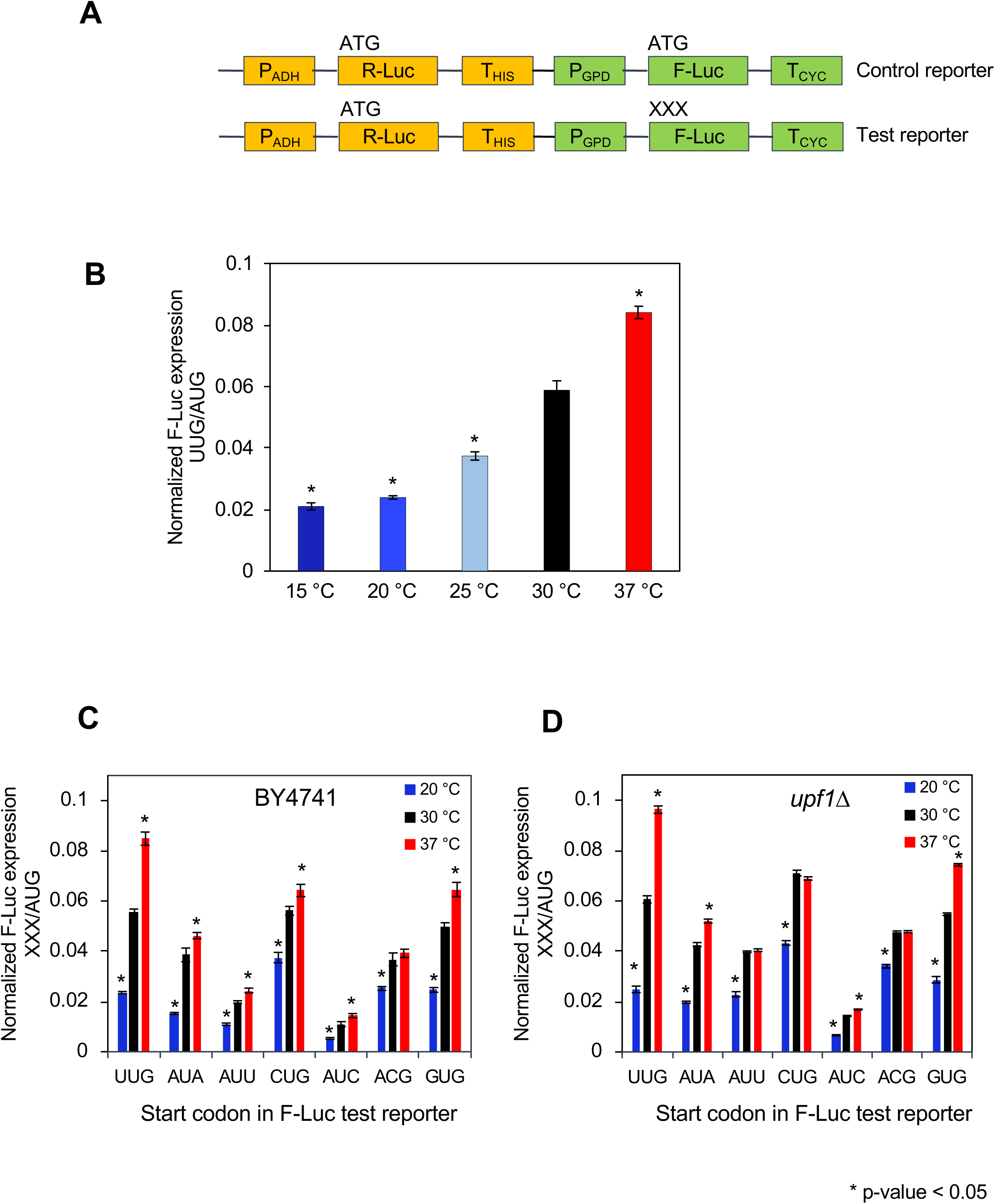
Changes in temperature result in changes in relative expression of firefly luciferase (F-Luc) reporters with AUG and near cognate codons (NCCs) as start sites. **(A)** Schematic of the reporters used in the study. (P) Promoter; (T) terminator. Renilla luciferase (R-Luc) mRNA was produced from the ADH promoter and HIS terminator and Firefly luciferase (F-Luc) mRNA was produced from the GPD promoter and CYC terminator. ‘XXX’ in the F-Luc test reporter represents a start codon that varies from AUG by 1 bp (NCC). The relative expression from firefly luciferase with respect to renilla luciferase was calculated for both test (F-Luc^XXX^/R-Luc^AUG^) and control (F-Luc^AUG^/R-Luc^AUG^) reporter. The normalized expression was then calculated as the ratio of relative F-Luc^XXX^ expression from the test reporter to relative F-Luc^AUG^ expression from the control reporter (XXX/AUG ratio). **(B)** Normalized F-Luc^UUG^ expression (UUG/AUG ratio) was measured in yeast strain BY4741 at different temperatures. **(C)** Normalized expression from F-Luc reporters (XXX/AUG ratio) with different NCCs as start sites in BY4741 cells cultured at multiple temperatures. **(D)** Same as C, except the experiments were done in Δ*upf1* cells. For **B**, **C**, **D** results are represented as averages of at least 4 biological replicates with error bars representing standard error of mean. For B-D: * denotes p-values < 0.05 calculated by Student’s *t*-test when compared with 30 °C.

β-galactosidase activity assays in whole cell extracts (WCEs) were performed as described previously [22]. To assay expression of eIF1 (encoded by *SUI1*) by western blot analysis, the WCEs were made under denaturing conditions as described previously [23] from 4 biological replicates (independent transformants). The immunoblot analysis was performed [24] using antibodies against eIF1 [25] and Ded1 (a kind gift from Tien-Hsien Chang). Two technical replicates were performed using the same extracts and two-fold different amounts of each extract were loaded in two successive lanes. Enhanced chemiluminescence (Amersham #RPN2106) was used to detect the immune complexes using a ProteinSimple imager (FluorChem systems #FM0261) and the signal intensities were quantified by densitometry using Adobe Photoshop after inversion of the image as described [26].

### Ribosome profiling

Cell culture and lysis: BY4741 cells harboring pR^AUG^F^UUG^ plasmid (dual luciferase reporter with R-Luc(AUG)-F-Luc(UUG) in a *URA3* vector (see Additional file 2: Supplementary Table 2) were used for the ribosome profiling, which was performed as described [8, 19, 27, 28] with some modifications. 750 ml of yeast cells in log phase (OD_600_ 0.6-0.8) grown in synthetic complete medium lacking uracil (SC-Ura) at either 20 °C, 30 °C or 37 °C for 16 hours, were harvested rapidly (≤ 1 min) by vacuum filtration at room temperature and snap-frozen in liquid nitrogen. No cycloheximide was added to the media before harvesting to avoid cycloheximide-induced artifacts [29–32]. This was followed by addition of ice-cold lysis buffer (20 mM Tris [pH 8.0], 140 mM KCl, 1.5 mM MgCl_2_, 1% Triton, 500 µg/mL cycloheximide). The concentration of cycloheximide in this lysis buffer was 5X the concentration originally used in ribosome profiling experiments [8] to minimize continued translation elongation during lysis and processing. The frozen pellets in lysis buffer were lysed in a freezer mill (Freezer/Mill® Dual Chamber Cryogenic Grinder, #6870, with the settings of 15 cycles, at 15 Hz, 5 minutes precool, 1 minute run followed by 2 minutes cooling). The frozen lysate was transferred to a 50 ml conical tube and thawed on ice with frequent agitation. The cell lysate was centrifuged at 3,000 rpm (Eppendorf #5810R), at 4 °C for 5 minutes, and the supernatant was collected and further centrifuged at 13,000 rpm (∼18,000 rcf) in a table top centrifuge (Eppendorf #5417R) at 4 °C for 10 minutes. The final supernatant was collected and the OD_260_ was measured. Aliquots of lysate containing 30 OD_260_ units were stored at −80 °C until used.

Isolation of 80S ribosomal footprints for sequencing library preparation: An aliquot of lysate containing 30 OD_260_ units, was thawed on ice and treated with 500 U of RNAse I (Ambion™ #AM2294)_for 60 minutes at 26 °C in a thermomixer at 700 rpm. 5 μl of SUPERase-In RNAse inhibitor (Ambion™ #AM2694) was added to the reaction, which was then used for isolation of 80S monosomes as previously described [27]. Briefly, to isolate 80S monosome, the reaction was loaded on a 10-50% (w/v) sucrose gradient made in lysis buffer, followed by centrifugation for 3 hours at 40,000 rpm in a SW 41 Ti rotor. The fractions were separated on density gradient fractionation system (Brandel) using 60% sucrose solution (made in lysis buffer) to pump the gradients. The fraction/s corresponding to 80S were collected and the ribosome-protected mRNA fragments (RPFs) were purified with SDS/hot acid phenol and chloroform.

Isolation of total mRNA for RNA-seq library preparation: Total RNA was isolated from the lysate using miRNeasy Mini Kit (Qiagen #217004) following the manufacturer’s protocol. The random fragmentation was carried out by addition of fragmentation reagent (Ambion #AM8740) and incubation at 70 °C for 8 minutes, followed by addition of stop solution (from the same kit).

Sequencing library construction: The ribosome protected fragments (RPFs) and fragmented total mRNA were each resolved by electrophoresis on a 15% TBE-Urea gel (Novex #EC68852BOX). Following the size selection, the RNA was gel extracted and was dephosphorylated using polynucleotide kinase (NEB #M0201S). A Universal miRNA cloning linker (NEB # S1315S) was ligated to the 3’-ends of the RNA using T4 RNA Ligase 2 (NEB #M0242L) in presence of PEG 8000 (2.5% w/v), DMSO and SUPERase-In at 37 °C for 2.5 hours. The ligated products were resolved by electrophoresis on 15% TBE-Urea gel and the appropriate size fragments eluted from the gel. The linker-ligated RNA recovered from RPFs was directly subjected to reverse transcription, while that extracted from fragmented total RNA samples was used as input for the Ribozero reaction (Illumina Ribo-Zero Gold rRNA Removal Kit-Yeast) to remove rRNAs, and subsequently subjected to reverse transcription. For reverse transcription, 10 μl RNA (dissolved in 10 mM Tris pH 8.0) from the previous reaction was mixed with 2 μl of 1.25 µM reverse transcription primer [5′- (Phos)AGATCGGAAGAGCGTCGTGTAGGGAAAGAGTGTAGATCTCGGTGGTCGC(SpC18)CACTA(SpC18)TTCAGACGTGTGCTCTTCCGATCTATTGATGGTGCCTACAG)], denatured at 80 °C for 2 minutes and incubated on ice. This was followed by addition of SuperScript™ III Reverse Transcriptase (#AM2694), dNTPs, DTT and Superase-In and incubation at 48 °C for 30 minutes. The RNA template was removed by addition of 2.2 μl of 1N NaOH followed by incubation at 98 °C for 20 minutes. The reaction was resolved on a 15% TBE-Urea gel and the cDNA was extracted from the gel. Circularization using CircLigase (Epicentre #CL4111K) was conducted by mixing 15 μl of cDNA (dissolved in 10 mM Tris pH 8.0) with 2 μl of 10X CircLigase buffer, 1 μl of 1 mM ATP, 1 μl of 50 mM MnCl_2_, and 1 μl of CircLigase. The reaction was incubated at 60 °C for 1 hour, followed by heat inactivation at 80 °C for 10 minutes.

The circularized products were used as templates for PCR amplification of the total RNA sample, or first subjected to subtractive hybridization to remove rRNA-derived sequences in case of RPF libraries. For the latter, the circularization reaction was mixed with a subtraction pool of biotinylated oligonucleotides [28] in the presence of 2X SSC (#AM9763), denatured at 100 °C for 90 seconds, followed by annealing at 37 °C. The reaction was then mixed with Dynabeads and incubated at 37 °C in a Thermomixer at 1000 rpm. The eluate was recovered and treated as the rRNA-depleted sample. The latter, as well as the circularized product derived from total RNA, were used as templates for PCR amplification to produce sequencing libraries using previously published primers [28] with slight changes in the barcodes. The resulting libraries were sequenced using the Illumina HiSeq system at the DNA Sequencing and Genomics Core, NHLBI, NIH. The reads were trimmed to remove the linker sequences (using fastx_toolkit/fastx_trimmer [http://hannonlab.cshl.edu/fastx_toolkit/index.html] in the relevant command line code), and then aligned to the *S. cerevisiae* rRNA database using Bowtie [33]. The reads that did not align with the reference database (non-rRNA reads) were aligned to the *S. cerevisiae* genome or to the FASTA file made from the reporter sequence (pR^AUG^F^UUG^) to generate alignments to the F-Luc reporter mRNA using TopHat [34].

### Data analysis, statistics and webtools

For ribosomal profiling, each experiment was performed with 2 independent cultures as biological replicates. The statistical analysis between the replicates was performed using DESeq2 [35] The DESeq2 statistical package addresses the typical challenge in next-generation sequencing experiments of having only two biological replicates for each condition by pooling information about the variances of read counts across the thousands of genes being analyzed in order to model count variances for genes of similar expression levels. The modeled variances are used in the framework of a generalized linear model (GLM), an extension of linear regression allowing for non-normal error distributions, to identify expression changes and place confidence intervals on the magnitude of changes, and also to exclude genes showing aberrantly high variability. Transcriptional and translational changes can be analyzed together in a GLM by including library type (mRNA-Seq or Ribo-Seq) as one of the factors, in addition to experimental variables like genotype or drug treatment, in a multi-factor design. The translational efficiency (TE) emerges as the effect of the Ribo-seq library type against the mRNA-Seq baseline, and significant interactions of TE with the experimental variables indicate translational control [36]. While the analysis is under-powered with two versus three biological replicates, only the chance of false-negatives, not false-positives, is increased. Using DESeq2 we calculated the changes in the ribosome density (ribosome protected fragments per million mapped reads), mRNA density (RNA-seq reads per million mapped reads) and the translation efficiency (TE, ribosome density/mRNA density) at different temperatures using a cutoff of ≥ 10 average mRNA reads in four samples. To calculate TE of a uORF (TE_uORF_) mRNA read counts for the mORF were employed rather than the mRNA read counts in the uORF alone. Spearman’s correlation was calculated using an online tool at https://www.wessa.net/rwasp_spearman.wasp. Notched box and whisker plot analysis was conducted using a webtool http://shiny.chemgrid.org/boxplotr/. The notches are defined as ±1.58*interquartile range /√n and represent the 95% confidence interval for each median. Non-overlapping notches give roughly 95% confidence that two medians differ. Heatmaps were generated using a webtool ‘Heatmapper’ (http://www.heatmapper.ca/) [37].

### Wiggle tracks

A combined alignment file (Bam file) was generated using two alignment files (one for each of the two biological replicates). The combined files were generated for both RPF samples and total RNA samples (for 20 °C, 30 °C and 37 °C). Wiggle files were generated from this combined alignment file. Files were generated for each gene on the Watson or Crick strand. The tracks were visualized using Integrative Genomics Viewer (IGV 2.4.14). The tracks were normalized according to total number of mapped reads in the combined file. To normalize the effects of changes in mRNA levels, the total read-normalized peaks were scaled with respect to the changes in mRNA levels to reflect the changes in translation efficiency (see Results section). Wiggle tracks, both with and without the scaling by changes in mRNA levels, are shown.

### Finding upstream ORFs

We took a similar approach to identify possible uORFs and confirm their translation in our experiments as that described previously [19]. First, putative translated uORFs were identified essentially as described [38]. Briefly, for all open reading frames in annotated 5’-UTRs that initiate from either AUGs or near cognate codons, the ratio of RPF counts at the +1 position (start codon of uORF) to −1 position (upstream of the start codon) was calculated. Those uORFs with ratios > 4, with >14 RPF counts at the +1 and −1 positions combined, and with at least 50% of the counts reads in the 0-frame with respect to the start site (i.e. the relevant line of code is -c15-r4-z0.5), were selected for further analysis. The multiple ribosome profiling datasets we used to identify potential uORFs were described previously [19] and have been submitted to the NCBI Gene Expression Omnibus and the accession numbers are listed https://elifesciences.org/articles/31250/figures#supp1 in the additional file table S2.

To determine which of these putative uORFs were translated in our experiments, we employed an ORF identification tool (RibORF) described previously [39], which uses 3-nucleotide periodicity and a uniform distribution of RPFs counts across the uORF as scoring criteria. We applied a moderately stringent cutoff of the probability of prediction of 0.5 and used a combined alignment file generated from footprint libraries of all 6 biological replicates grown at three different temperatures to identify uORFs with evidence of translation at one or more growth temperatures. After excluding uORFs shorter than three codons, we identified 1367 uORFs starting with AUG (N = 142) or a NCC (N = 1225). Quantifying the total mRNA and RPF counts in the 5’-UTR, uORF or mORF was done as previously described [19].

To identify potential uORFs in all yeast mRNA 5’-UTR transcriptome, 5’-UTR sequences for all mRNAs were extracted, and each AUG and NCC nucleotide triplets were searched throughout the 5’-UTR. We identified the first in frame stop codon for each start codon as the end of the uORF. For uORFs without an in-frame stop codon in 5’-UTR, we defined the end of the 5’-UTR as the end of uORF. uORFs with less than 3 codons in length were excluded in downstream analysis.

## Results

### 3.1 Growth temperature affects the efficiency of using non-AUG start codons in reporter mRNAs in yeast

We previously developed and validated a dual luciferase assay to calculate the efficiency of utilization of near-cognate codons (NCCs) as translational start sites in yeast [20]. In this assay, Renilla luciferase (R-Luc) and Firefly luciferase (F-Luc) are expressed using separate promoters and transcription terminators from a single low-copy plasmid (Figure 1A). R-Luc mRNA has an AUG as the start site and acts as an internal control for cell growth, lysis efficiency and pipetting inconsistency. The start codon of the F-Luc reporter is varied and can be AUG or any NCC (e.g. UUG, ACG, etc.). F-Luc expression is normalized to that of R-Luc to control for any global changes in gene expression or cell growth. F-Luc^AUG^/R-Luc^AUG^ represents the normalized expression value for a F-Luc reporter starting with AUG. The relative expression from UUG (or any other near-cognate) is calculated by normalizing with respect to the normalized F-Luc^AUG^ expression (F-Luc^UUG^/R-Luc^AUG)^)/(F-Luc^AUG^/R-Luc^AUG^).

We used this assay to investigate the effects of changes in growth temperature on the use of NCCs as translational start sites. Yeast cells (BY4741) were transformed with the dual luciferase reporter plasmid with either an F-Luc^AUG^ or F-Luc^UUG^ gene, cultured at various temperatures, and the luciferase activity of the F-Luc^UUG^ reporter relative to the F-Luc^AUG^ reporter was measured. (F-Luc and R-Luc assays in cell extracts were performed at 24 °C regardless of the temperature at which the yeast cells were cultured.) Elevating the growth temperature from 30 °C to 37 °C led to ∼1.5-fold increase in the normalized expression of F-Luc^UUG^, while lowering the growth temperature from 30 °C to 25 °C or 20 °C led to ∼1.6-fold and ∼2.5-fold reduction, respectively, in the normalized F-Luc^UUG^ expression (Figure 1B). These findings suggest that at lower growth temperatures the efficiency of use of the F-Luc UUG start codon is decreased, and that at higher temperatures it is increased.

To try to confirm these findings and test the generalizability of the observed effects of temperature on start codon usage, we performed a similar experiment with two otherwise identical *HIS4-lacZ* fusion reporters with an AUG or UUG start codon [40] (Additional file 1: Figure S1A). The normalized expression of a *HIS4-lacZ* reporter with UUG as a translational start site was reduced ∼5-fold at 20 °C with respect to 30 °C and was elevated ∼2.8-fold at 37 °C (Additional file 1: Figure S1B). These results are consistent with the findings from the dual luciferase reporter that the use of a UUG relative to an AUG start codon is reduced at 20 °C and increased at 37 °C. Frequently, the finding that two orthogonal reporter assays give similar results might be taken to indicate that the observed effect is generalizable to most mRNAs. However, as described below, this turns out not to be the case in this system.

We next tested the effects of changes in growth temperature on the normalized expression from F-Luc reporter mRNAs starting with other NCCs (GUG, CUG, ACG, AUA, AUC and AUU), all of which have been shown to be utilized as start sites in yeast cells to varying extents [4, 20]. Like UUG, the normalized expression from all NCCs was significantly lowered (∼2 fold) at 20 °C indicating that the effects of lowering the growth temperature were not specific to the UUG start site (Figure 1C). On the other hand, elevation in growth temperature resulted in differential changes in normalized expression, ranging from no change for ACG to ∼30% increase (for AUC and GUG) as compared to their expression at 30 °C. This suggested that the efficiency of use of NCCs might be differentially affected at some temperatures.

The changes in expression of the F-Luc reporters starting with different initiation codons could be due to changes in mRNA stability induced by the nonsense-mediated decay (NMD) pathway triggered by altered translation of an upstream open reading frame (uORF) starting from an NCC. In the absence of an AUG start codon, it is possible that an upstream or out-of-frame NCC codon is used for initiation leading to premature termination, and potentially NMD [41]. Alternatively, it is also possible that the scanning 43S pre-initiation complexes bypass the NCC start site (leaky scanning), initiate at an out-of-frame AUG or NCC in the mORF, terminate in the mORF, and thereby trigger NMD [42]. To test these possibilities, we performed the luciferase assay in a strain in which the *UPF1* gene, which encodes a protein essential for NMD, had been deleted (*upf1Δ*) [43]. Deletion of *UPF1* did not affect the temperature-dependent changes in expression observed in WT cells for the majority of the start codons tested at both 20 °C and 37 °C (Figure 1D). This result suggests that the NMD pathway does not play a role in the observed changes in expression from these reporters at different temperatures.

The alterations in the use of UUG as a start site could be attributed to changes in the levels of eIF1, which has been shown to be a ‘gatekeeper’ in the start codon recognition process [44], helping to restrict start codon selection to AUGs and block the selection of NCCs. To test the possibility that temperature-dependent changes in eIF1 expression might be responsible for the observed effects of temperature on start codon usage, we assessed the levels of the factor using western blot analysis of whole cell lysates from cells cultured at different temperatures. Levels of eIF1 protein were not significantly altered at 20 or 37 °C relative to 30 °C (Additional file 1: Figure S2A), suggesting changes in start codon use are not due to changes in eIF1 concentration.

We also tested the effect of over-expression of eIF1 on the observed temperature-dependence of F-Luc start codon utilization. Over-expression of eIF1 from a high-copy (hc) plasmid has been shown to suppress the reduced stringency of start codon recognition (Sui^-^) phenotype caused by mutations in several initiation factors [45–48]. Consistent with its role as a central gatekeeper of start codon recognition, over-expression of eIF1 (hc-*SUI1*) suppressed the use of UUG as a start site at all three temperatures (Additional file 1: Figure S2B). The decrease in F-Luc^UUG^ expression at 20 °C and the increase at 37 °C were still observed in the hc-*SUI1* strain, although the magnitude of the increase at 37 °C was reduced relative to WT cells in this experiment. Consistent with these results, reducing the concentration of eIF1 relative to WT cells by using a *SUI1*/*sui1Δ* heterozygous diploid strain, resulted in increased expression of F-Luc^UUG^ relative to F-Luc^AUG^ at all three temperatures relative to expression in a WT diploid containing two wild-type chromosomal alleles of *SUI1* (Additional file 1: Figure S2C). No decrease in the magnitude of the temperature-dependence of normalized F-Luc^UUG^ expression was observed in the *SUI1*/*sui1*Δ haploinsufficient diploid. In addition, haplo-insufficiency of eIF1A (+/*tif11*Δ) or eIF5 (+/*tif5*Δ), also factors involved in start codon recognition, did not significantly alter the effect of temperature on expression of F-Luc^UUG^ relative to F-Luc^AUG^ (Additional file 1: Figure S2C). It is noteworthy that, for reasons unknown, the increase at 37 °C is dampened in the WT *SUI1/SUI1* diploid versus the *SUI1* haploid strain analyzed in Figure 1A, and that the larger differences resurface in the *SUI1*/*sui1Δ* heterozygote. Although altering the dosage of the *SUI1* gene appears to modulate somewhat the effects of 37 °C on UUG initiation, overall it appears that the effects of temperature on NCC utilization are not dictated by altered cellular levels of eIF1.

Taken together, these results suggest that the effect of temperature on F-Luc^UUG^ expression is not due to changes in the concentrations of eIF1, eIF1A or eIF5. Additionally, when we performed ribosome profiling in cells grown at different temperatures (see below) we did not observe any obvious changes in the ribosome protected fragments (RPF) counts and translational efficiencies for the mORF of eIF1, further confirming that the level of eIF1 does not change as a function of growth temperature in WT haploid cells (Additional file 1: Figure S2D).

### 3.2 Ribosome profiling elucidates temperature-dependent changes in start codon utilization transcriptome wide

To investigate the effects of changes in growth temperature on the relative use of different codons as translational start sites throughout the transcriptome, we performed ribosome profiling in yeast cells cultured at multiple temperatures. WT yeast cells (BY4741) transformed with the F-Luc^UUG^ reporter plasmid were cultured in SC-Ura for 16 hours at 20, 30 and 37 °C and ribosome profiling was performed (Figure 2A) as previously described [8, 27], with some notable changes (see Methods section). In particular, we did not add cycloheximide to the intact cells to stop translation because of the known artifacts it creates [29–32], but instead flash froze the cells in liquid nitrogen and added cycloheximide to the cell lysis buffer only.

**Figure 2.**
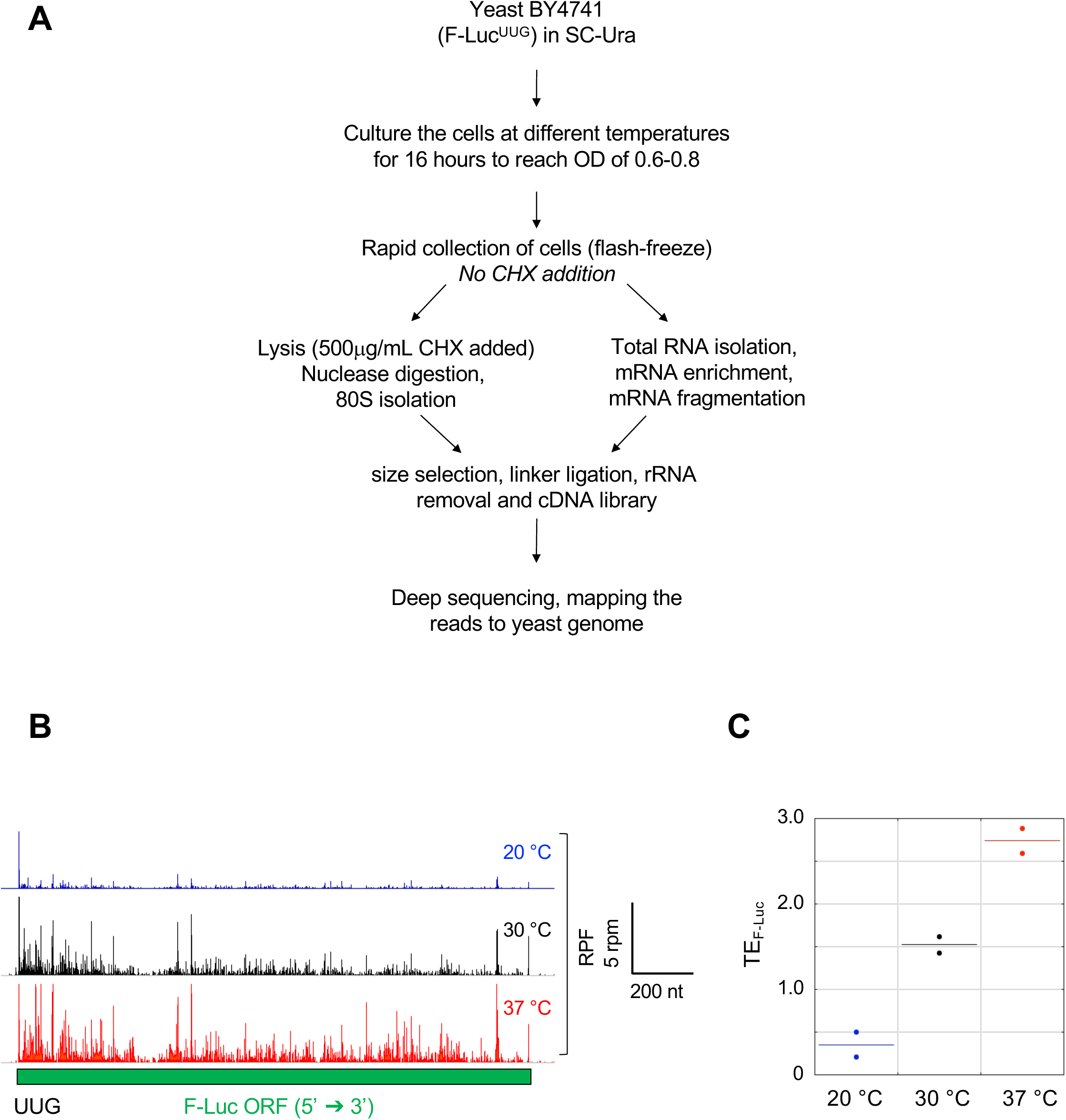
Ribosome profiling under multiple temperatures. **(A)** Schematic of the ribosome profiling experiment. BY4741 cells harboring the F-Luc^UUG^ reporter plasmid were grown at multiple temperatures and then transcriptome-wide translation was analyzed by ribosome profiling. **(B)** Wiggle track image of F-Luc^UUG^ reporter mRNA at multiple temperatures. Ribosome-protected fragments (RPFs) on the F-Luc^UUG^ reporter mRNA in cells cultured at 20, 30 or 37 °C, in units of rpm (reads per million mapped reads from two replicates at each temperature). The RPF-tracks were normalized to the mRNA levels (see methods) at each temperature to reflect the changes in translation efficiencies (ΔTE). **(C)** Translation efficiency (TE) values, calculated as ribosome density (RPF reads on F-Luc mRNA normalized to total number of RPF reads mapped) divided by the mRNA density (mRNA reads of reporter mRNA normalized to total number of mRNA reads mapped) (ribo-density/mRNA-density), for the F-Luc^UUG^ reporter mRNA from both biological replicates at 20 °C (blue), 30 °C (black) and 37 °C (red). Each point represents the TE-value for the F-Luc reporter from one replicate and the horizontal solid line represents the mean.

We calculated ribosomal-read density as the number of 80S ribosomal footprint reads mapped to an mRNA sequence relative to the total number of reads in the footprint library (ribo-seq), and we calculated the mRNA read density by normalizing RNA-seq reads mapped to an mRNA sequence relative to the total number of reads in the RNA-seq library. The translation efficiency (TE) for each mRNA is calculated as ribosomal read density normalized to mRNA read density [8]. The ribosome footprint and RNA-seq data indicate that the two biological replicates were highly reproducible for all the temperatures (Pearson’s r > 0.98; Additional file 1: Figure S3A).

We mapped the ribo-seq (ribosome protected fragments, RPFs) and RNA-seq reads to the F-Luc^UUG^ mRNA reporter to confirm our findings with the reporter assay. The expression of F-Luc^UUG^ mRNA reporter was significantly altered at multiple temperatures (Figure 2B and Additional file 1: Figure S3B). The RPF read count for the F-Luc mRNA was decreased at 20 °C by ∼25% and increased at 37 °C by ∼46% as compared to 30 °C (Additional file 1: Figure S3B). In contrast, the mRNA-read count for the F-Luc mRNA was increased at 20 °C by ∼300%, and decreased at 37 °C by ∼20%, as compared to 30 °C (Additional file 1: Figure S3B). To control for the changes in mRNA abundance, we calculated the TE of the F-Luc reporter (Figure 2C). TE for F-Luc^UUG^ mRNA was reduced at 20 °C by ∼77% and elevated at 37 °C by ∼80%, as compared to 30 °C, indicating that the translation of F-Luc mRNA is significantly altered at 20 °C and 37 °C, consistent with the findings from the luciferase assay where the normalized expression of this reporter (with respect to the F-Luc^AUG^ control reporter, Figure 1A) was altered in an analogous manner (Figure 1B, C).

### 3.3 Identification of upstream open reading frames (uORFs) using ribosome profiling data

To better understand the effects of changes in temperature on start site selection, we identified a set of 1367 uORFs that show evidence of translation in our yeast strain at one or more temperatures. To this end, we employed a two-step strategy for translated uORF discovery described previously [19] (Figure 3A). The first step employs the Yassour-uORF algorithm [38] in which putative translated uORFs are identified from among the set of all possible uORFs initiating with an AUG or NCC on the basis of a strong peak of ribosome density at the start codon and the occurrence of >50% of downstream read counts in the zero frame of the start codon. After excluding uORFs shorter than three codons, we identified 6061 potential uORFs by applying this algorithm to several previously reported ribosome profiling datasets (see Methods). In the second step, we examined which of these 6061 putative uORFs show evidence of translation in our ribosome profiling data using a different identification tool, RibORF [39], which is based on the criteria of 3-nt periodicity (a hallmark of mRNA fragments protected by actively translating ribosomes) and a uniform distribution of reads across uORF codons. This tool generates a *predicted translating probability* ranging from 0 to 1. Lower probability values indicate skewed distributions of reads and equally distributed fractions of reads at the zero, 1^st^ and 2^nd^ reading frames. Higher values indicate a uniform distribution of reads and a majority of reads aligned in the zero reading frame. Applying a moderately stringent probability of prediction of >0.5, we found evidence for translation in our datasets for 1367 uORFs among the 6061 potential uORFs detected in the first step, located on 755 different mRNAs. These uORFs were further analyzed for changes in their expression at different temperatures using DESeq2 (see methods).

**Figure 3.**
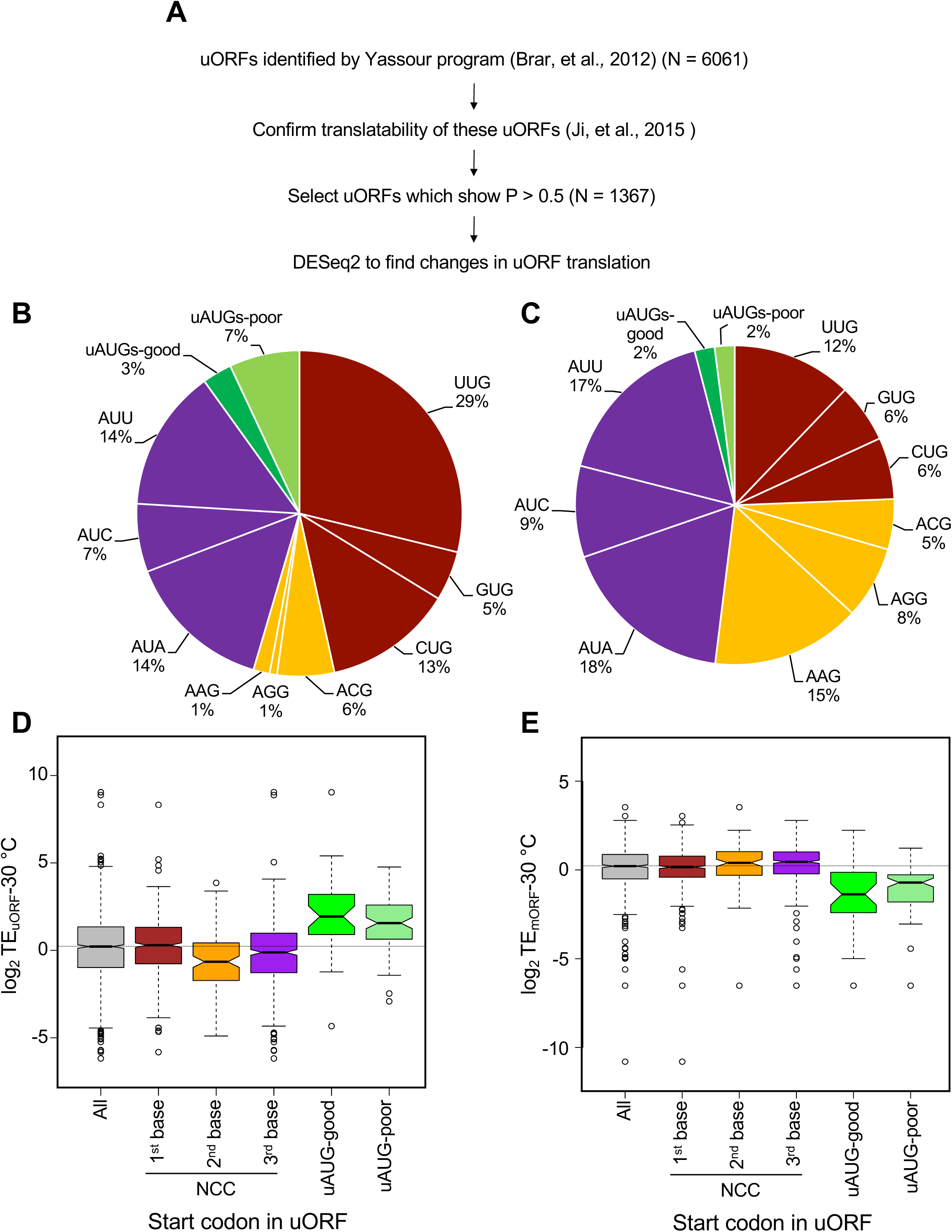
Identification of upstream Open Reading Frames (uORFs) in *S. cerevisiae*. **(A)** Workflow for identifying uORFs. Multiple datasets described previously in [19] were used to identify potential uORFs in yeast using a program described previously [38]. The potential uORFs ≥ 3 codons (N = 6061) that were identified were run through a program previously described [39] in order to determine which are translated in our data sets. 1367 uORFs showed evidence of translation in the combined dataset generated in this study (each replicate, all temperatures) at probability of prediction P > 0.5 (see Methods). The differential expression at 20 °C or 37 °C with respect to 30 °C was analyzed by DESeq2. **(B)** Distribution of start sites in translated uORFs (N = 1367) that were identified through the protocol described in **A**. The AUG uORFs were sorted into good context (A/G) or poor (U/C) based on the nucleotide at the −3 position with respect to AUG. Percentages were calculated with respect to the total number of translated uORFs identified using the pipeline described in **A**. **(C)** Transcriptome-wide distribution of potential upstream start sites including AUGs (uAUGs) and NCCs. The upstream start sites that can lead to formation of uORFs < 3 codons were excluded so that an appropriate comparison with the translated set of uORFs could be done. Percentages were calculated with respect to the total number of potential upstream start sites (N = 58,353). **(D)** The uORFs were binned into NCCs or uAUGs and then boxplot analysis of TE_uORF_ at 30 °C was performed. *All* indicates all the translated uORFs identified through our pipeline (N∼1350); *1^st^ base change* includes translated uORFs starting with UUG, CUG, GUG (N = 631); *2^nd^ base change* includes translated uORFs starting with AAG, ACG, AGG (N = 109); and *3^rd^ base change* includes translated uORFs starting with AUC, AUA, AUU (N = 483). AUG uORFs were sorted into good context (N = 36) or poor (N = 93). The dotted horizontal line indicates the median TE value of *All*. The notches indicate ±1.58*Inter Quartile Range (IQR)/√n, where IQR is the difference between the 75th and 25th percentiles and n represents the number of uORFs in that bin. Non-overlapping notches give roughly 95% confidence that the two medians differ. **(E)** Analysis of translation efficiency of main ORFs (mORFs) located downstream of the various translated uORFs. uORFs were binned depending on their start sites as described in **D**, and a boxplot analysis of TE_mORF_ at 30 °C was performed. *All* indicates the set of mORFs located downstream of all translated uORFs analyzed in this study (N = 748). The dotted horizontal line indicates the median TE_mORF_ of *All*.

These 1367 translated uORFs start with either an AUG (∼10%) or a NCC (∼90%). We observed a range of NCCs as start sites (Figure 3B), with UUG the most common (∼30% of all uORFs) and AGG the least (∼1% of all uORFs). The uORFs with near-cognate start codons with 2^nd^ base changes from AUG (AAG, ACG, AGG) contributed only 8% of all the uORF start codons, with ACG at 6% and both AAG and AGG at 1%,, which is consistent with previous findings indicating that AAG and AGG are the least efficiently used near-cognate start codons in yeast cells [4]. Near-cognate codons with 1^st^ base changes (UUG, GUG, CUG) comprised ∼50% of the total uORF start codons indicating that they are the most efficient near-cognate start sites, also consistent with previous studies [4] and our luciferase reporter analyses (Figure 1C). Among the ∼10% of all uORFs with an AUG start codon, ∼33% have a preferred Kozak context at the −3 position (A/G), while the remaining ∼66% have poor-context (U/C at −3 position) [49, 50] (Figure 3B), which is consistent with previous studies suggesting that good context uAUGs have been selected against evolutionarily [9].

We also looked at the overall abundance of AUGs and NCCs in the 5’-UTR transcriptome after removing any that initiate predicted uORFs less than three codons in length (in order to match the conditions used for identification of translated uORFs in our datasets, which also eliminated potential uORFs less than three codons long; Figure 3C). Comparison of the start codon distribution of the set of translated uORFs we identified (Figure 3B) with the 5’-UTR transcriptome abundance of potential uAUG and NCC start codons (Figure 3C) indicates that NCCs with first position changes are over-represented as start codons for the translated uORFs relative to their inherent abundance and those with second position changes are under-represented. These results are consistent with the general efficiency of use of these classes of NCCs as start codons. Also as expected, AUG codons are used more frequently as start codons for translated uORFs than their representation among potential start codons in 5’-UTRs.

### 3.4 uORFs with different NCCs as start sites show differential translation efficiencies

To investigate the translatability of the uORFs we calculated translational efficiency (TE) at 30 °C for all the uORFs starting with AUGs or NCCs. TE, as described above, is the ratio of ribosomal footprint read density to mRNA read density. As shown in Figure 3D, uORFs starting with different initiation codons were translated with differing median efficiencies at 30 °C. As might be expected, the median TE for uORFs starting with AUG codons (AUG uORFs) was significantly higher than the medians for uORFs starting with any NCC (NCC uORFs). In this and all other box and whisker plots below, lack of overlap in the notches of two adjacent plots indicates that their medians differ with >95% confidence (Chambers et al., 1983 Graphical Methods for Data Analysis. Wadsworth, Bellmont). The median TE for AUG uORFs in good context was comparable to that for AUG uORFs in poor context (Figure 3D), suggesting that other features of these mRNAs might modulate the effect of sequence context for this subset of uORFs.

The median TEs for NCC uORFs varied by a factor of about 2 to 3-fold, depending on which base varied from AUG. Consistent with our analysis of the start codon distribution of translated uORFs (Figure 3B), uORFs starting with NCCs with 2^nd^ base changes were the least efficiently translated, whereas uORFs starting with NCCs with 1^st^ base changes were translated the most efficiently (Figure 3D). These data, together with the results described in Figure 3B, suggest differential recognition and utilization of NCCs as start sites for uORFs in yeast in a manner consistent with previous analyses of the efficiency of different start codons for main ORF translation [5, 6, 20].

We next examined the relationship between the translation efficiency of the uORFs and that of the downstream main ORF (mORF). We calculated the TEs for the mORFs downstream of each subset of uORFs grouped according to the uORF start codon. We observed that the median TEs for the mORFs downstream of AUG uORFs were significantly lower than any other group (Figure 3E), suggesting that uORFs with AUG start sites were typically inhibitory of downstream translation at 30 °C, as expected. The median TE of mORFs downstream of uORFs starting with near-cognate codons with 1^st^ base changes was essentially the same as for all mORFs with translated uORFs, whereas mORFs downstream of the more poorly translated uORFs starting with near-cognates with 2^nd^ or 3^rd^ base changes had a slightly higher median TE compared to all mORFs (compare “All” in Figure 3E with 1^st^, 2^nd^, 3^rd^ base changes and uAUGs). Overall, these data provide a transcriptome-wide view of uORF translation and the inverse correlation between uORF and mORF TE values. These data are consistent with previous reports that the presence of uAUGs in 5’-UTRs of yeast mRNAs is inversely correlated with the polysome density of the mRNAs [9].

### 3.5 Changes in growth temperature have varying effects on uORF translation for different mRNAs

To examine the effects of changes in growth temperature on the translation of AUG and NCC uORFs, we calculated the changes in TEs of uORFs at either 20 °C or 37 °C with respect to 30°C. The TE of a uORF (TE_uORF_) at any given temperature was defined as the ribosome footprint (RPF) density of the uORF divided by the mRNA density of the downstream mORF. To calculate the changes in TE_uORF_ (ΔTE_uORF_) at either 20 °C or 37°C as compared to 30 °C, we performed DESeq2 analysis. As described in Methods, DESeq2 is a statistical package using the framework of a generalized linear model (GLM) that can identify changes in RPF and mRNA densities, as well as TEs, for each ORF between two conditions, place confidence intervals on the magnitude of changes, and exclude genes with less than a minimum number of read counts or with aberrantly high variability. Applying this analysis to our data revealed greater changes in uORF RPF densities versus the corresponding mORF mRNA densities for cells grown at 20 °C versus 30 °C (Figure 4A), and also for cells grown at 37 °C versus 30 °C (Figure 4B), as indicated by the greater spread in the uORF RPF density versus mORF RNA density scatterplots [panels (ii) vs. (i)]. These findings suggest that the changes in growth temperature led to more extensive changes in translation of uORFs than transcription and/or stability of the mRNAs containing the uORFs.

**Figure 4.**
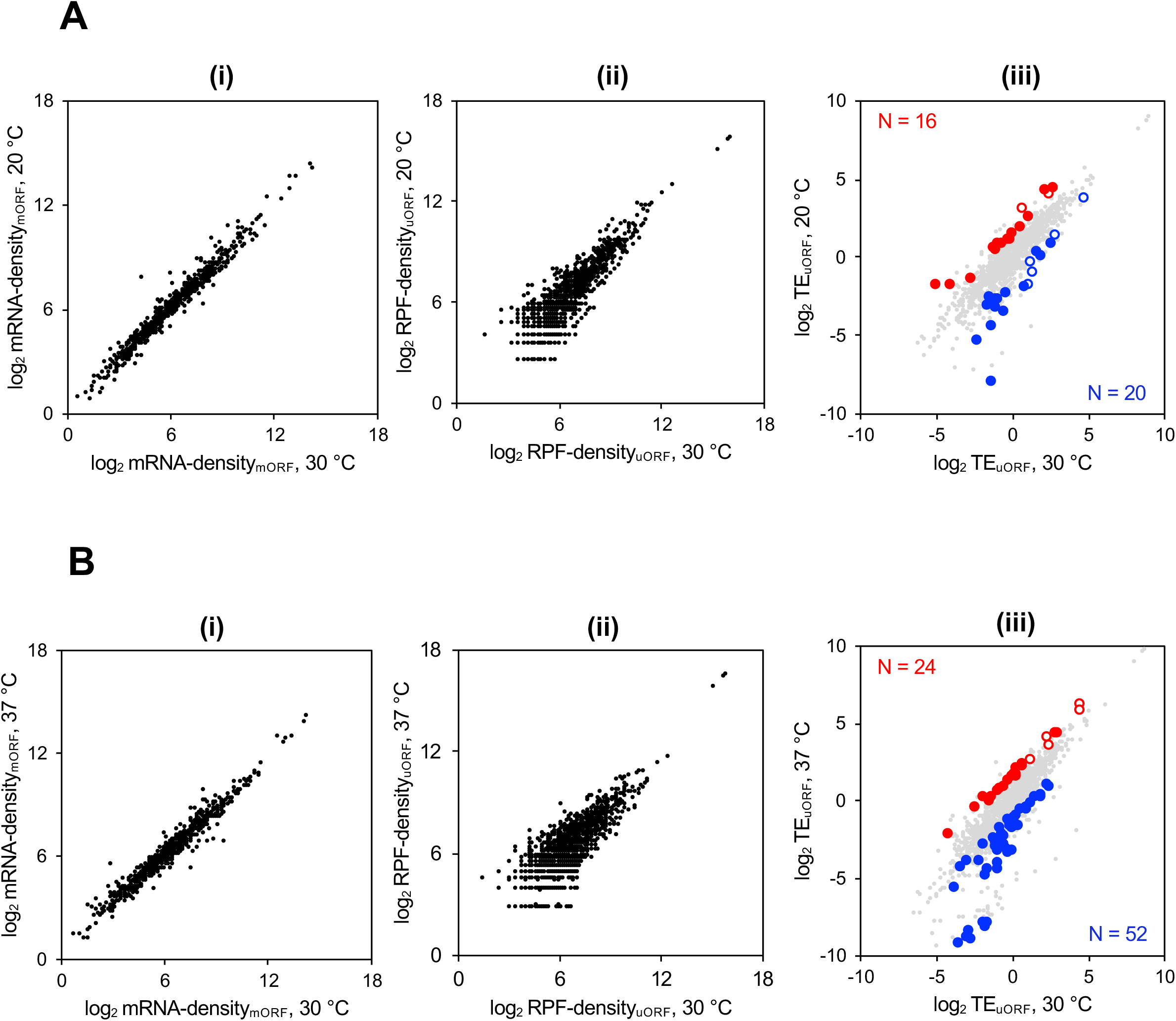
Changes in temperature result in changes in translation of some uORFs. (A, B) Scatterplots of normalized read densities. The mRNA density_mORF_ was calculated as the number of RNA-seq reads mapped to the corresponding mORF coding region normalized to the total number of RNA-seq reads. The RPF density_uORF_ was calculated as the number of ribosomal footprint reads (ribosome protected fragments) mapped to the uORF coding region normalized to the total number of RPF reads. TE_uORF_ was calculated as RPF density_uORF_ divided by mRNA density_mORF_ at either 20, 30 or 37 °C. **(A)** Scatterplots of mRNA densities (panel (i)), RPF densities (panel (ii)), and translational efficiencies (TEs; panel (iii)) for cells grown at 30 °C (X axis) and 20 °C (Y axis). The scatterplot of TEs also shows uORFs exhibiting ≥ 2-fold changes in both TE_uORF_ and relative TE (TE_uORF_/TE_mORF_) at a False Discovery Rate (FDR) < 0.1. Red circles represent uORFs meeting these criteria whose translation is activated at 20 °C as compared to 30 °C (N = 16), while blue circles represent uORFs whose translation is repressed at 20 °C as compared to 30 °C (N = 20). The open circles are AUG uORFs while the filled circles are NCC uORFs. **(B)** Same as in **A**, except the circles represent read densities for cells grown at 30 °C (X axis) and 37 °C (Y axis). As before, the scatterplot of TE_uORF_ [panel (iii)] shows uORFs with ≥ 2-fold changes in both TE and relative TE at FDR < 0.1. Red circles represent uORFs whose translation is activated at 37 °C as compared to 30 °C (N = 24), while blue circles represent uORFs whose translation is repressed at 37 °C as compared to 30 °C (N = 52). The open circles are AUG uORFs while the filled circles are NCC uORFs.

To identify uORFs showing changes in TE that appear to be activated or repressed by a change in growth temperature, we applied two criteria. First, we considered only those uORFs showing an increase or decrease in TE_uORF_ of ≥ 2-fold at a given temperature with respect to 30°C using a false discovery rate (FDR) of ≤ 0.1. 39 uORFs showed significant TE changes at 20°C versus 30 °C; whereas 84 uORFs displayed such TE changes at 37 °C versus 30 °C (Additional file 1: Figure S4A, S4B). We reasoned that changes in TE_uORF_ could result because of multiple mechanisms. For example, translation initiation on the mRNA as a whole could increase or decrease because of changes in the efficiency of PIC attachment or scanning processivity, leading to corresponding increases or decreases in the TEs of both the uORF(s) and mORF. To exclude such changes in TE_uORF_ occurring concurrently with similar changes in TE_mORF_, we devised a term called ‘relative-TE_uORF_’ which is TE_uORF_/TE_mORF_ at any given temperature. Calculating changes in Relative-TE_uORF_ (ΔRelative-TE_uORF_) helped to identify changes in uORF translation not occurring simultaneously with similar changes in the TE of the mORF. Thus, according to our second criterion, translation of a uORF was called ‘*regulated’* if there was ≥2-fold change (increase or decrease) in relative-TE_uORF_ at a given temperature with respect to 30°C; that is, TE_uORF_ changed ≥2-fold more than TE_mORF_, or their changes were in opposite directions. Applying these criteria, we identified uORFs whose translation is specifically regulated by changes in growth temperature. We classified uORF translation as *activated* if both ΔTE_uORF_ and ΔRelative-TE_uORF_ are ≥ 2 and as *repressed* if both ΔTE_uORF_ and ΔRelative-TE_uORF_ are ≤ 0.5.

After applying these criteria, we found 36 uORFs showing temperature dependent translational regulation at 20 °C. There were 16 uORFs whose translation was significantly activated (Figure 4A, red circles in panel (iii)) of which 2 were AUG uORFs (open circles) and 14 were NCC uORFs (solid circles). There were 20 uORFs whose translation was significantly repressed at 20 °C (Figure 4A, blue circles in panel (iii)) as compared to 30 °C, of which 5 were AUG uORFs (open circles) and 15 were NCC uORFs (solid circles). We also found 76 uORFs showing temperature dependent translational regulation at 37 °C. There were 24 uORFs whose translation was significantly activated (Figure 4B, red circles in panel (iii)) as compared to 30 °C, of which 7 were AUG uORFs (open circles) and 17 were NCC uORFs (solid circles) and 52 uORFs whose translation was significantly repressed at 37 °C (Figure 4B, blue circles in right panel) all of which are NCC uORFs. The changes in translation efficiency of uORFs (ΔTE_uORF_) were driven by changes in ribosome density (ΔRPF-density) and not by changes in mRNA levels (ΔmRNA-density) for the uORFs regulated at either 20 °C or 37 °C (Additional file 1: Figure S4C), as well as for all 1359 translated uORFs identified in this study (Additional file 1: Figure S4D), as revealed by the high Spearman correlation coefficient values between ΔTE_uORF_ and ΔRPF-density, and low coefficient values between ΔTE_uORF_ and ΔmRNA-density.

It has been reported that alternative transcription start sites can produce mRNA isoforms in yeast with different translational efficiencies [51]. Thus, we also calculated Spearman correlation coefficients between ΔTE_uORF_ and the changes in reads of just the 5’-UTRs of these sets of mRNAs (ΔmRNA_5’UTR_-density) and found that they are much smaller than those for ΔRPF-density and are not statistically significant (Additional file 1: Figure S4C and S4D, orange bars). This result suggests that the changes in TE_uORF_ observed are not due to temperature-dependent alterations in transcriptional start sites that produce different levels of mRNA isoforms including or excluding the uORFs in question. Furthermore, high Spearman correlation coefficient values between ΔTE_uORF_ and ΔRPF-density, and low coefficient values between ΔTE_uORF_ and both ΔmRNA-density and mRNA_5’UTR_-density were observed for all translated AUG and NCC uORFs (Additional file 1: Figure S4E and S4F), indicating that for both these sets of uORFs the changes in translational efficiency were driven by changes in RPF density and not in mRNA or 5’-UTR density.

The F-Luc^UUG^ and *HIS4^UUG^*-*LacZ* reporters showed decreased translation at 20 °C and increased translation at 37 °C (Figures 1B, 2B, Additional file 1: Figure S1B) suggestive of altered efficiency of use of the NCCs. In contrast to these reporters, we found changes in growth temperature lead to a more diverse transcriptome-wide response of translation of uORFs starting with not only NCCs but also AUGs. As described above, we identified 112 uORFs (14 AUG uORFs and 98 NCC uORFs) (Additional file 2: Supplementary Table 3) on 84 different mRNAs whose translation was activated or repressed in response to changes in growth temperature. It is noteworthy that less than 10% of the set of 1359 translated uORFs have significantly altered translation relative to changes in mORF translation at reduced or elevated growth temperatures (20 or 37 °C), indicating that temperature changes in this range do not produce global effects on the initiation efficiency of uORFs, but rather have specific, mRNA-dependent effects.

### 3.6 Influence of uORF start codon sequence on temperature-dependent changes in translation

The distribution of start codons of the 112 regulated uORFs described above is shown in Figure 5A and B. We separated the uORFs based on their start sites into four classes: uAUGs; NCCs with 1^st^ base changes with respect to AUG (UUG, CUG, GUG), 2^nd^ base changes (AAG, ACG, AGG) and 3^rd^ base changes (AUC, AUA, AUU). The number of uORFs in each bin is too small to make inferences about the statistical significance of the differences. Thus we next analyzed the changes in TE (ΔTE_uORF_) of all 1359 translated uORFs by binning them into groups based on their start codon triplets without applying the criteria used to identify significant changes (Figure 5C, D). The black horizontal dotted line indicates the median ΔTE_uORF_ for all uORFs analyzed in this study at each temperature, which is close to unity. The NCC uORFs did not show a significant difference in median ΔTE_uORF_ at 20 °C versus 30 °C when compared to all uORFs (Figure 5C). In contrast, the AUG uORFs showed a significantly lower median ΔTE_uORF_ at 20 °C when compared to all uORFs, suggesting that use of uAUGs tends to be decreased at 20 °C.

**Figure 5.**
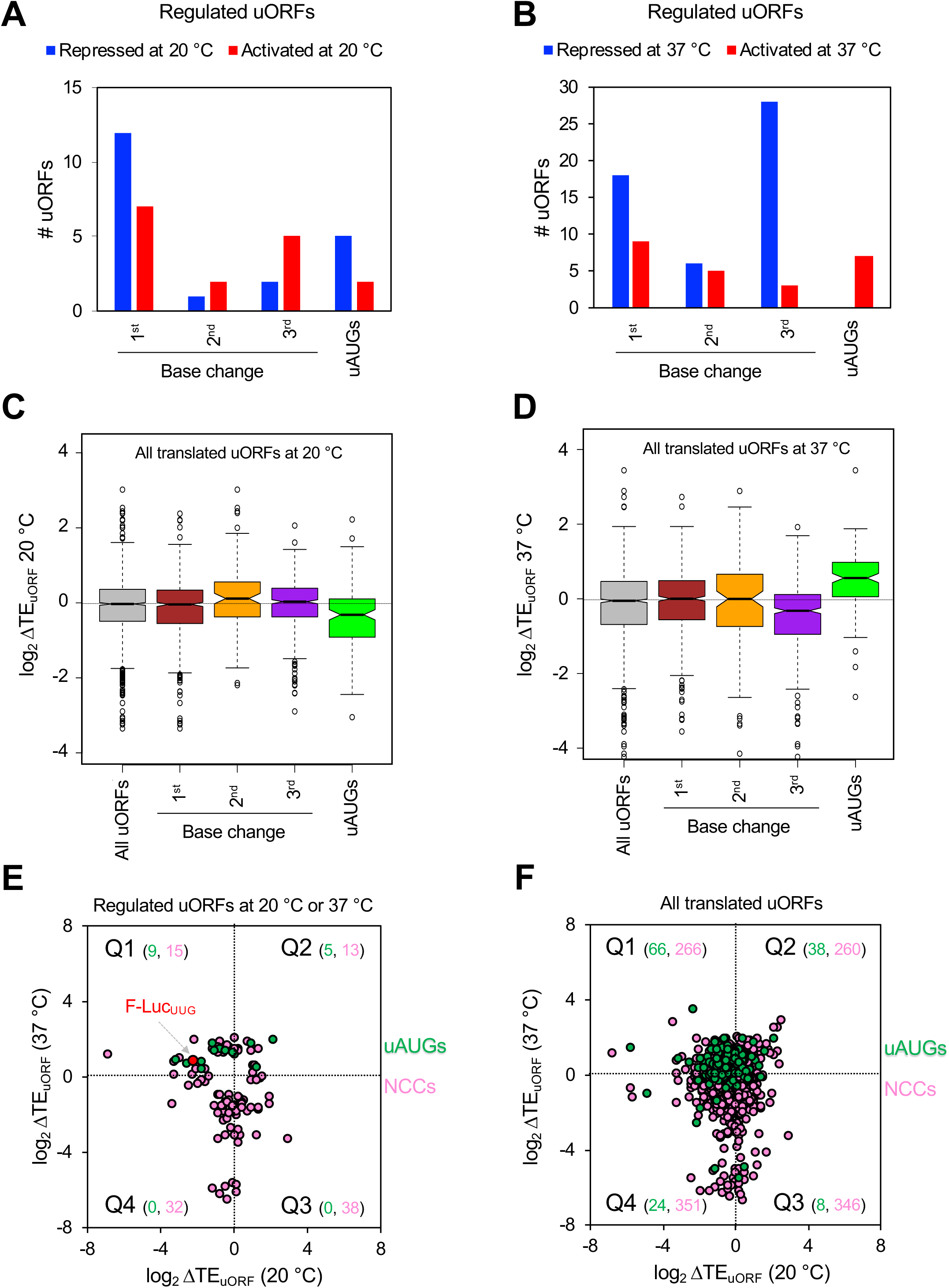
Varied response to temperature of uORF translation. (A, B) Responses of uORF translation to growth temperature grouped by start codons triplets. The temperature-regulated uORFs [whose translation is repressed or activated at either 20 °C or 37 °C, see figures 4A and 4B; highlighted in red or blue in panel (iii)], were binned according to their start site. *uAUGs* represents AUG uORFs. *Base change* (with respect to AUG) corresponds to uORFs starting with UUG, GUG, CUG (*1^st^*), ACG, AGG, AAG (*2^nd^*), or AUA, AUC, AUU (*3^rd^*). **(A)** Analysis done for changes observed at 20 °C relative to 30 °C. **(B)** Analysis done for changes observed at 37 °C relative to 30 °C. **(C, D)** Notched boxplot analysis for changes in TE_uORF_ (ΔTE_uORF_) on all the uORFs starting with each near-cognate start codon triplet at either 20 °C or 37 °C with respect to 30 °C. *All uORFs* indicates all translated uORFs identified and analyzed in this study (N = 1359); *uAUGs* represent all translated AUG uORFs (N = 136); *1^st^ base change* (N = 631) represents uORFs starting with UUG, CUG, GUG; *2^nd^ base change* (N = 109) represents uORFs starting with ACG, AAG, AGG; and *3^rd^ base change* (N = 483) represents uORFs starting with AUA, AUC, AUU. The dotted horizontal line shows the median for *All uORFs*. **(C)** Analysis done for ΔTE_uORF_ at 20 °C with respect to 30 °C. **(D)** Analysis done for ΔTE_uORF_ at 37 °C with respect to 30 °C. **(E, F)** Scatterplots of ΔTE_uORF_ at 20 °C (X axes) versus ΔTE_uORF_ at 37 °C (Y axes). **(E)** The ΔTE_uORF_ values of temperature-regulated uORFs [Figure 4A and 4B, panel (iii)]. Green circles represent AUG uORFs (uAUGs, N = 14) and pink circles represent NCC uORFs (NCCs, N = 98). The plot is broken into four quadrants, Q1-4, depending on the direction of ΔTE_uORF_ at each temperature. The numbers in parenthesis represent the number of uORFs present in that specific quadrant (AUG uORFs in green and NCC uORFs in pink). The red circle shows the main ORF of F-Luc^UUG^ reporter mRNA. **(F)** Same as in **(E)**, except all translated uORFs identified in this study whose changes in TE could be determined, are shown. Green circles represent AUG uORFs (N = 136) and pink circles represent NCC uORFs (N = 1223).

We performed a similar analysis with uORF translation at 37 °C (Figure 5D). The TE of uORFs starting with NCCs with 3^rd^ base changes had a significant tendency to be downregulated at 37 °C, whereas those with 1^st^ and 2^nd^ base changes showed no clear trend. AUG uORFs displayed a strong overall increase in TE_uORF_ (positive ΔTE_uORF_) at 37 °C. Thus, use of uAUGs is significantly altered at both temperatures: reduced at 20 °C and elevated at 37 °C), similar to the behavior of the F-Luc and *HIS4*-LacZ reporters with UUG start codons. In contrast to AUG uORFs, the translation of NCC uORFs displays no clear trends in response to changes in growth temperature, save for reduced utilization of NCCs with 3^rd^ base changes at 37 °C.

To better understand the global translational response of uORFs to the changes in growth temperature, we plotted ΔTE_uORF_ at 20 °C versus ΔTE_uORF_ at 37 °C (both with respect to 30 °C). First, we examined the 112 uORFs showing temperature dependent translational regulation at either 20 °C or 37 °C (Figure 5E). The distribution of these uORFs on the scatterplot reflects their translational behavior at each growth temperature relative to 30 °C: repression at 20 °C and activation at 37 °C (Quadrant 1); activation at both temperatures (Quadrant 2); repression at 37 °C and activation at 20 °C (Quadrant 3); repression at both temperatures (Quadrant 4). The position of the F-Luc^UUG^ reporter in Quadrant 1 (red circle) indicates translational regulation at altered temperatures, with decreased expression at 20 °C and increased expression at 37 °C. Similarly, the majority (9/14) of the AUG uORFs (Figure 5E, green circles) are positioned in Quadrant 1, indicating that translation of regulated uORFs starting with AUGs tends to be decreased at 20 °C (log_2_TE_uORF_ < 0) and increased at 37 °C (log2TE_uORF_ > 0). All of the AUG uORFs that met our criterion for regulated changes in translation are in Quadrants 1 or 2, indicating that translation of AUG uORFs is generally increased at 37 °C relative to 30 °C. The regulated NCC uORFs (Figure 5E, pink circles), on the other hand, showed a scattered distribution across all the quadrants indicating that changes in translation of these NCC uORFs is more variable.

We performed a similar analysis with all translated uORFs (N = 1359) (Figure 5F). The overall distribution on this scatter plot is similar to that for the regulated uORFs in Figure 5E, except that many uORFs whose translation is unaffected by temperature are present near the middle of the plot. The significant numbers of points representing NCC uORFs along the vertical axis between Quadrants 3 and 4 in both plots (Figure 5E and 5F, pink circles) indicates that many of these uORFs are repressed at 37 °C relative to 30 °C, but are relatively unaffected at 20 °C, consistent with the box plot analyses of these same uORFs (Figure 5C,D). As with the AUG uORFs that met the criteria for regulated changes (Figure 5E), 66/136 of all translated AUG-uORFs are repressed at 20 °C and activated at 37 °C (Figure 5F, green circles; Quadrant 1) and most (104/136) are activated at 37 °C (Figure 5F, green circles; Quadrants 1 and 2), indicating that this is a general phenomenon. In contrast, NCC uORFs have less coherent behavior upon change in growth temperature and can be unaffected, activated or repressed (Figure 5F, pink circles) in an apparently mRNA-specific manner.

Recently, 982 uORFs were identified from *S. cerevisiae* in 791 mRNAs using a comparative genomics approach to identify translated uORFs that are conserved in length or sequence among yeast species [10]. Approximately 44% of these are AUG uORFs and ∼31% are UUG uORFs. When we interrogated this conserved uORF set, we found that, similar to our observations with the translated uORFs described above, translation of the conserved AUG uORFs is significantly repressed at 20 °C and activated at 37 °C (Additional file 1: Figure S5A-C, green boxplots and circles). Intriguingly, the TE of the conserved NCC uORFs is on average slightly elevated at 20 °C and slightly repressed at 37 °C (Additional file 1: Figure S5A-C, pink boxplots and circles). Thus, a set of conserved AUG uORFs identified in a different manner than was our set of translated AUG uORFs displays a similar overall response to temperature.

### 3.7 Translated AUG uORFs tend to be in shorter, less structured 5’-UTRs than do NCC uORFs

To look for possible mechanisms influencing translation of NCC and AUG uORFs, we investigated whether intrinsic properties of these uORFs or their mRNAs display any significant correlations with uORF translational efficiencies. We first used a dataset of ∼2700 yeast mRNA 5’-UTR lengths and propensities of forming secondary structures [52] to look for trends in the translated uORFs. We found that the 5’-UTRs of mRNAs with translated uORFs are on average longer and have a higher propensity to form secondary structure than the genomic average (Additional file 1: Figures S6A-C; see Supplementary Methods for details). This result might be expected because shorter 5’-UTRs have less space in which to have a uORF and fewer possibilities for base pairing. Translated AUG uORFs tend to be significantly closer to the 5’-cap (Additional file 1: Figure S6D) and shorter (Additional file 1: Figure S6F) than are all translated uORFs or NCC uORFs. AUG uORFs also tend to be on shorter 5’-UTRs (Additional file 1: Figure S6G) and have less overall structure in their 5’-UTRs and in their mORFs near the start codon (Additional file 1: Figures S6H, I, K, L, M) compared to the translated NCC uORFs (green versus pink boxplots) or all translated uORFs (green versus gray boxplots). No statistical difference is seen between AUG and NCC uORFs or all uORFs when distance from the mAUG is compared (Additional file 1: Figure S6E).

Next, we calculated the ‘context adaptation scores’ for the uORFs, as described previously [19, 53], quantifying the similarity between the start codon context of each uORF to that of the mORF AUGs of the 2% of yeast mRNAs with the highest ribosomal loads [54]. The start codons of the AUG uORFs have a significantly lower context score when compared to NCC uORFs (Additional file 1: Figure S6N, green versus pink boxplots) or all translated uORFs (green versus gray boxplots), consistent with the notion that strong AUG codons in the 5’-UTR have likely been selected against evolutionarily [9]. Overall, these data indicate that AUG uORFs, on average, occur on shorter, less structured 5’-UTRs closer to the cap, and exhibit poorer context, compared to NCC uORFs.

### 3.8 Position of a uORF within a 5’-UTR influences its translational response to altered growth temperatures

We next investigated whether any of these intrinsic properties of the translated uORFs and their mRNAs influence the temperature-dependent regulation of uORF translation by calculating the Spearman’s correlation coefficients between ΔTE_uORF_ and the features analyzed in Figures S6D-N (Additional file 1). We found a modest but significant positive correlation between the distance from the 5’-cap and ΔTE_uORF_ at 20 °C versus 30 °C for all uORFs, AUG uORFs and NCC uORFs (Figure 6, top panel). We also found a significant positive correlation between the length of the 5’-UTR and ΔTE_uORF_ at 20 °C for all NCC uORFs and all uORFs. Similar correlation was observed for AUG uORFs, but they did not meet statistical significance because of the smaller number of these uORFs. At 37 °C, the distance between the uORF start codon and the mAUG, the length of the uORF, and the length of the 5’-UTR all had a significant positive correlation with ΔTE_uORF_ for all uORFs and NCC uORFs (Figure 6, bottom panel). Again, similar correlations were observed for these same parameters for AUG uORFs, but they did not meet statistical significance because of the smaller number of these uORFs. Taken together, these data suggest that the position of a uORF in the 5’-UTR and the length of the 5’UTR can influence how translation of the uORF responds to changes in growth temperature.

**Figure 6.**
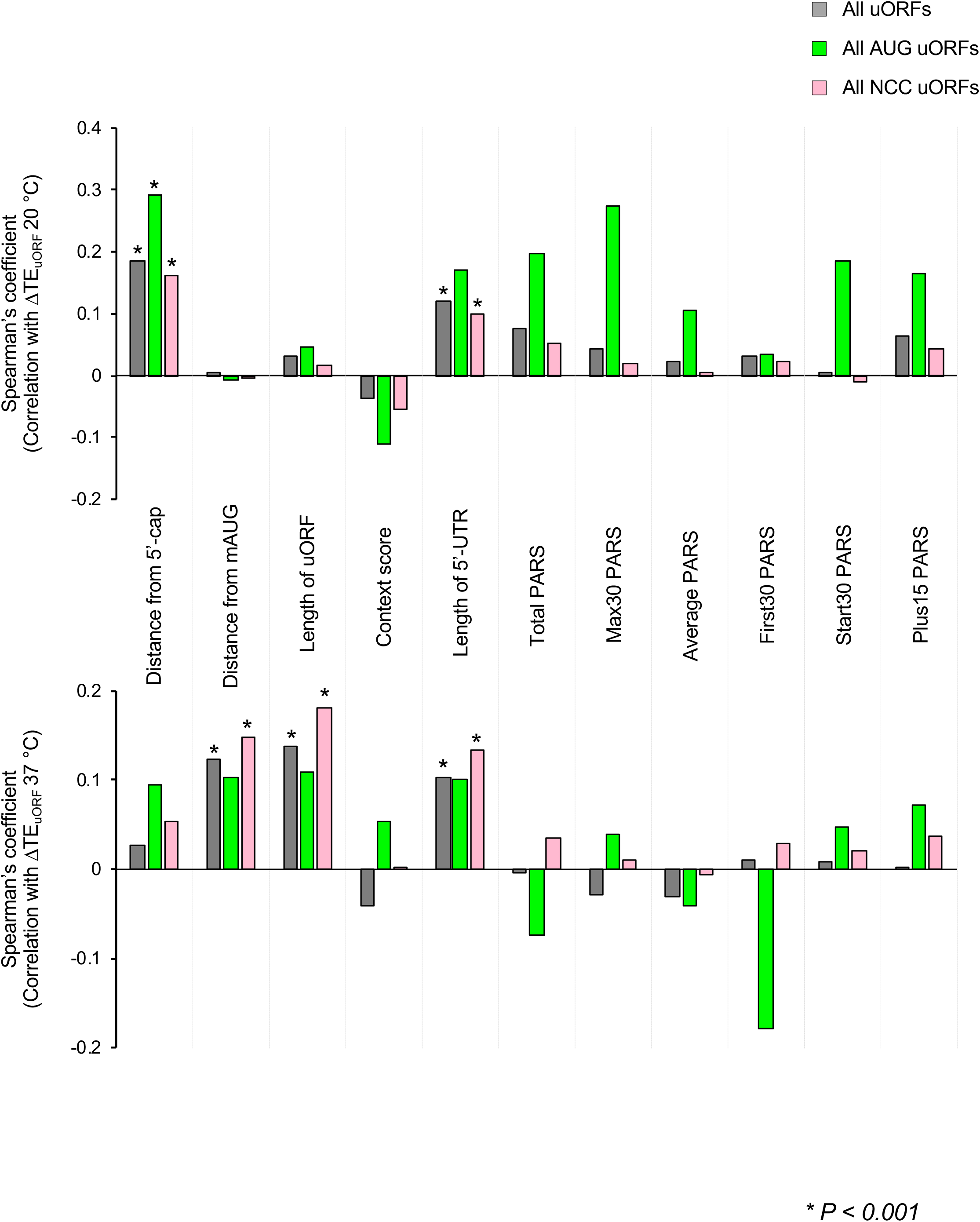
Position of a uORF within a 5’-UTR influences its translational response to altered growth temperatures. Spearman’s correlation coefficients between ΔTE_uORF_ at 20 °C (top panel) or 37 °C (bottom panel) and the parameters described in Additional file 1: Figure S6. The uORFs whose ΔTE could be calculated were analyzed here. Coefficients with significance levels of p < 0.001 are indicated with asterisks (*).

In an effort to confirm these relationships between TE changes and uORF position relative to the cap or mORF, we examined the groups of NCC uORFs exhibiting the greatest TE changes at either high or low temperature. Sorting all ∼1200 translated NCC uORFs according to ΔTE_uORF_ values at 20 °C or 37 °C revealed that TE changes at both temperatures vary over an ∼1000-fold range, from 8-fold to ∼0.01-fold. We then selected the 100 uORFs with the largest increases in TE (‘TE_up’) or the largest decreases in TE (‘TE_down’) for subsequent analysis (Figure 7A, braces and dotted boxes). Boxplots of the TE values of TE_up and TE_down uORFs confirm that the median TE values of these groups of uORFs differ significantly between 30 °C and 20 °C (Figure 7B), or between 30 °C and 37 °C (Figure 7C).

**Figure 7.**
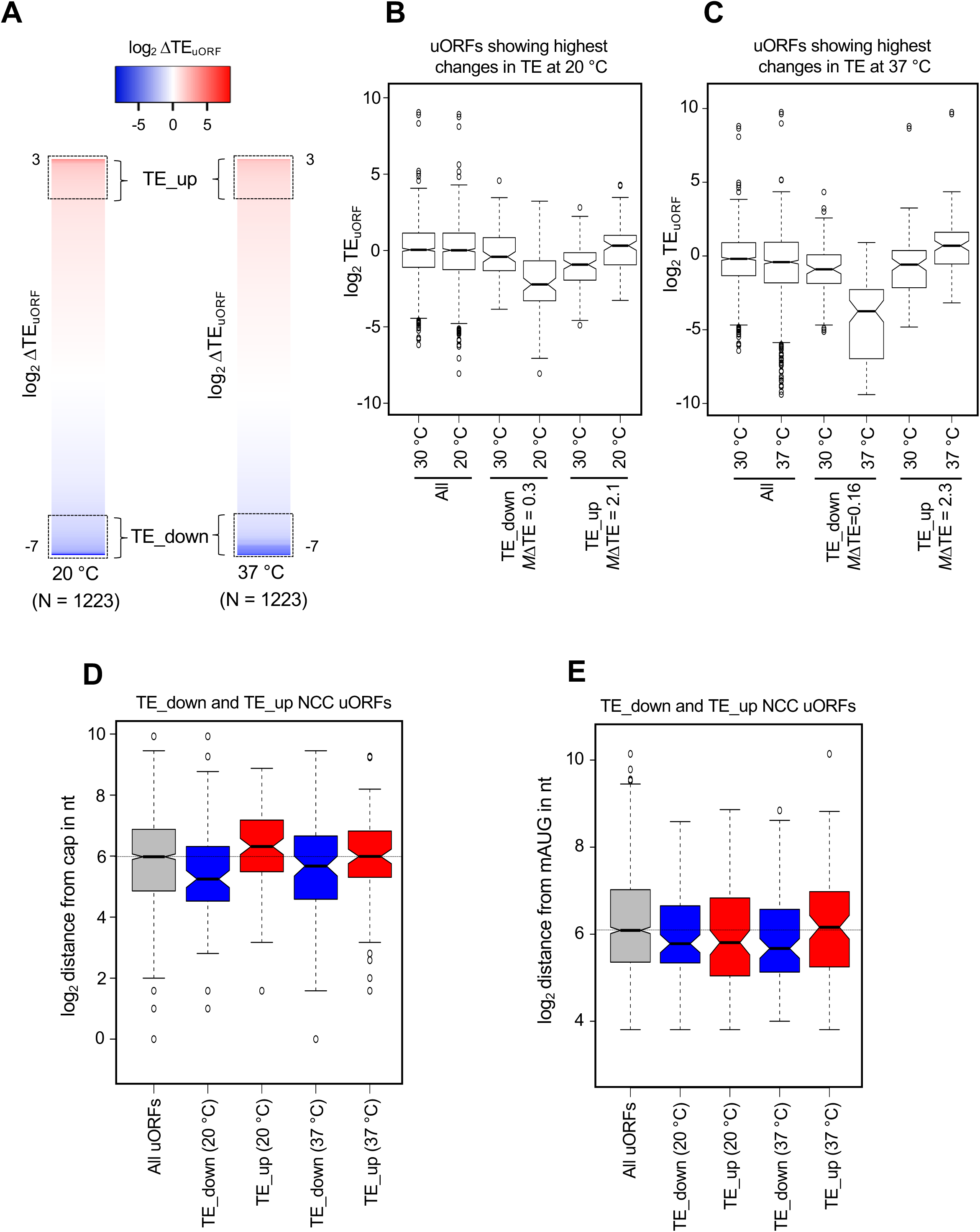
Identification and analysis of NCC uORFs with the greatest changes in TE at 20 °C and 37 °C. **(A)** Heat-maps of ΔTE_uORF_ values for all translated NCC uORFs (N = 1223), ranked according to ΔTE. The scale on the top indicates the range of ΔTE values. The dotted boxes indicate the 100 uORFs with the greatest decreases in TE_uORF_ (TE_down) or the greatest increases in TE_uORF_ (TE_up) at either 20 °C or 37 °C with respect to 30 °C. **(B)** Boxplot analysis of the 100 uORFs with the greatest decrease (TE_down) or increase (TE_up) in TE_uORF_ at 20 °C, identified as described in **A**, above. The median ΔTE_uORF_ values for each group are shown below the plot (mΔTE). *All* indicates all translated NCC uORFs (N = 1223). **(C)** Same as in **B** but for ΔTE_uORF_ at 37 °C. **(D)** Boxplot analysis of the distance between the start site of the uORF and the 5’-end of the mRNA. The dotted horizontal line indicates the median distance of All translated NCC uORF start codons from the 5’-ends of the mRNAs. **(E)** Boxplot analysis of the distance between uORF start sites and the downstream mORF start (mAUG) codons. The dotted horizontal line indicates the median distance of All translated NCC uORF start codons from the mORF start codons.

As shown in Figure 7D, the group of TE_down uORFs at 20 °C (col. 2) are located significantly closer to the 5’-cap relative to all NCC uORFs. In contrast, the TE_up uORFs at 20 °C (col. 3) are located significantly farther from the 5’-cap. The median distances between the 5’-cap and uORF start site for TE_down (20 °C), TE_up (20 °C), and all NCC uORFs are 38, 79, and 63 nt, respectively. These data are consistent with the correlation analysis performed on all translated uORFs (Figure 6, col. 1, upper plot, pink) in suggesting that NCC uORFs located farther from the cap tend to exhibit increased translation at 20 °C, whereas those closer to the cap tend to exhibit the opposite trend. In contrast, there is no significant difference in median distance from the cap for the two groups of NCC uORFs classified as TE_down and TE_up at 37 °C (Figure 7D, cols. 4-5). These data also are consistent with the correlation analysis performed for all translated NCC uORFs at 37 °C (Figure 6, col. 1, lower plot, pink).

The TE_down (37 °C) NCC uORFs are located closer to the mAUG (median 51 nt) compared to all NCC uORFs (median 68 nt) and the TE_up (37 °C) group of NCC uORFs (median 71 nt) (Figure 7E, columns 4 vs. 1 & 5). Although the TE_down (20 °C) NCC uORFs had a significantly shorter median distance from the mAUG compared to all NCC uORFs (median 55 and 68 nt, respectively), the two groups of NCC uORFs with TE_up or TE_down at 20 °C do not differ significantly from one another in this parameter (median 55 and 56 nt, respectively; Figure 7E, compare columns 1, 2 and 3). These results are consistent with the correlation analysis for all translated NCC uORFs (Figure 6, col. 3, upper & lower plots, pink) in suggesting that proximity to the mORF is associated with reduced translation of NCC uORFs at 37 °C but not at 20 °C.

We did not observe any significant influences of start codon context, the uORF length, or propensity to form secondary structure, on the temperature-dependent changes in translational efficiency for the TE_up or TE_down NCC uORFs at either temperature (Additional file 1: Figures S7A, S7C and S8A-F). The mRNAs with TE_down uORFs tend to have shorter 5’-UTRs than do the mRNAs with TE_up uORFs (Additional file 1: Figure S7B).

### 3.9 Disparate effects of changes in uORF translation on mORF translation

We next investigated the effects of the temperature-dependent changes of uORF translation on the translational efficiencies of their downstream mORFs. Scatterplots of mORF TEs at either 20 versus 30 °C or 37 versus 30 °C revealed the absence of widespread TE changes as a function of growth temperature (Additional file 1: Figures S9A and S9B); only 25 mRNAs exhibit ≥ 2-fold changes (activation or repression) in mORF TE (FDR < 0.1) at either 20 or 37 °C. To visualize the effects of changes in uORF translation on mORF expression, we plotted ΔTE_uORF_ versus ΔTE_mORF_ at 20 or 37 °C for our set of uORFs (described in Additional file 1: Figures S4A and S4B) showing significant changes in translation at either 20 or 37 °C (Figure 8A, B). Changes in uORF translation had varying effects on the translation of the mORF, which can be categorized according to the quadrant in which the uORF/mORF pair falls in the scatterplots in Figure 8. Cases in which a decrease in TE_uORF_ is accompanied by an increase in TE_mORF_ (Quadrant 1) or an increase in TE_uORF_ is associated with a decrease in TE_mORF_ (Quadrant 3) suggest that the uORF plays a canonical inhibitory role in terms of its effect on translation of the mORF. Quadrants 2 and 4 represent cases in which uORF and mORF translation both increase or decrease, respectively. The simplest explanation for this behavior is that overall initiation on these mRNAs (e.g., PIC loading onto the 5’-UTR) increases or decreases, leading by mass action to increases or decreases in translation of both the uORF and the mORF. Alternatively, it is possible that some cases in these two quadrants represent mRNAs on which re-initiation after translation of the uORF is very efficient and thus an increase or decrease in uORF translation has a corresponding effect on mORF translation.

**Figure 8.**
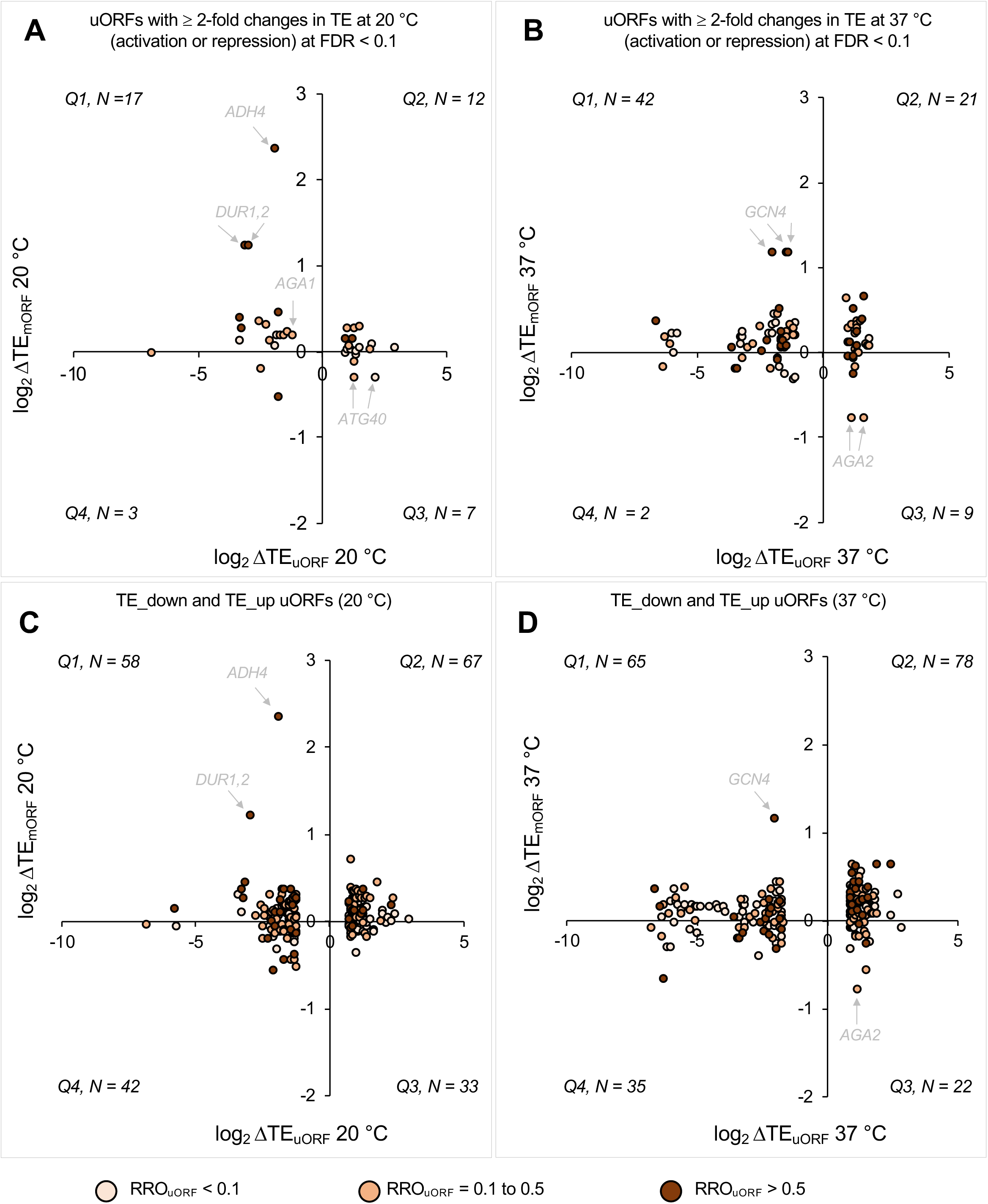
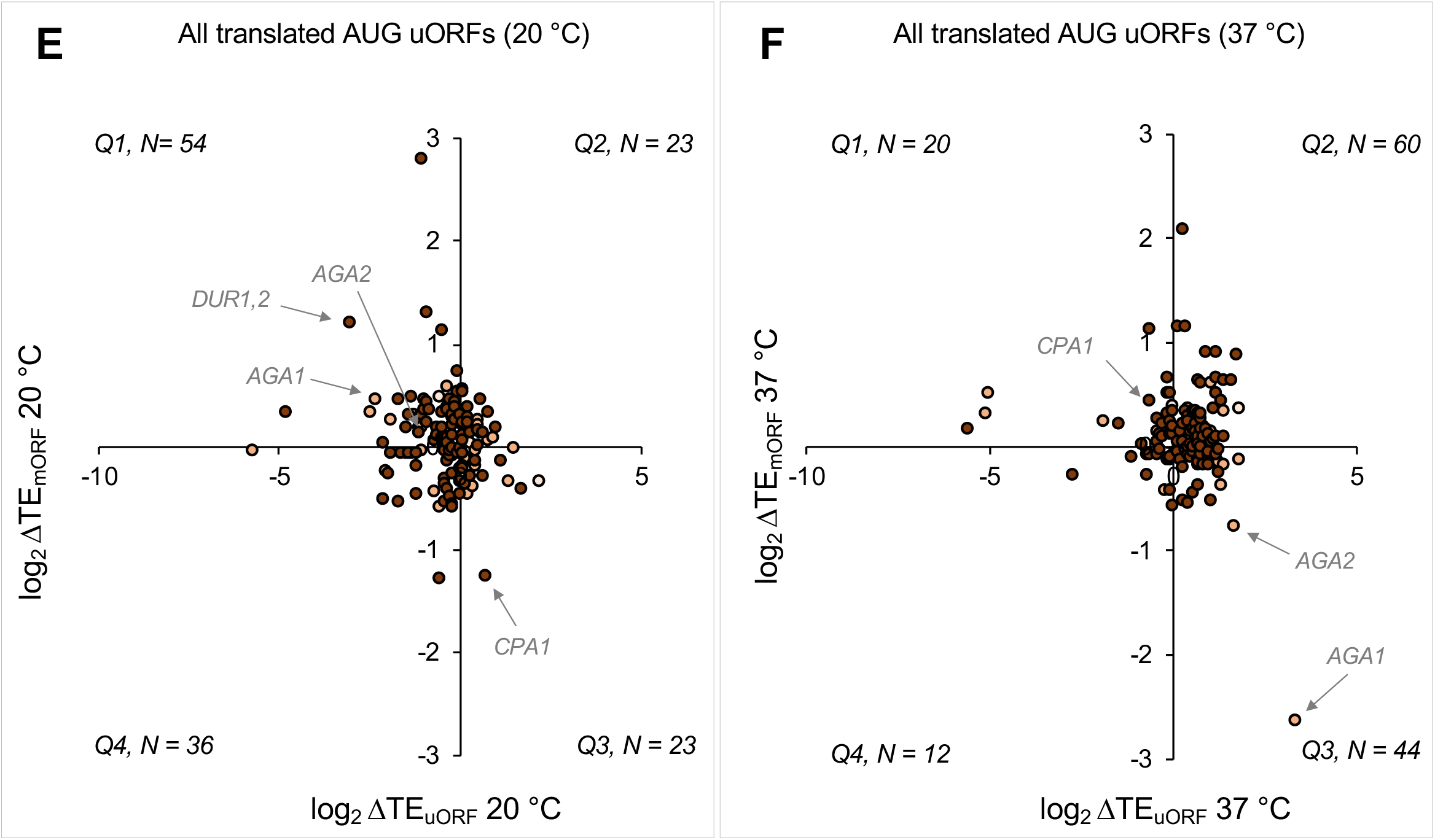
Correlations between changes in uORF and mORF translation suggest possible novel cases of uORF-mediated translational regulation. (A-F) Correlation analysis between ΔTE_uORF_ and ΔTE_mORF_ at multiple growth temperatures. Each circle in the plots represents a uORF. The circles are color-coded according to the relative ribosome occupancy of the uORF (RRO_uORF_; color key is shown at the bottom) which is the ribosome occupancy on the uORF (RO_uORF_) normalized to the ribosome occupancy on the mORF (RO_mORF_). RO_uORF_ is the RPF density on the uORF at 30 °C normalized to its length and RO_mORF_ is the RPF density on the mORF at 30 °C normalized to its length. The plot is divided into 4 quadrants (Q1-Q4). The number of uORFs in each quadrant is shown on every plot. Multiple sets of uORFs are used in this analysis. **(A, B)** The uORFs with ≥ 2-fold changes in TE (activation or repression) at FDR < 0.1 were analyzed (highlighted circles in red and blue in Additional file 1: Figures S4A and S4B). **(A)** Analysis of changes observed at 20 °C. **(B)** Analysis of changes observed at 37 °C. **(C, D)** NCC uORFs with the greatest changes in TE (TE_down and TE_up) at 20 °C **(C)** and 37 °C **(D)** as described in Figure 7A were analyzed. **(E, F)** All translated AUG uORFs (N = 136) were analyzed for changes in translation at 20 °C **(E)** and 37 °C **(F)**. For A-F, cases representing the uORFs with a possible canonical inhibitory function for the uORFs are indicated with an arrow.

We performed a similar analysis using the TE_up and TE_down NCC uORF sets described above and in Figure 7A-C (Figure 8C, D). As with the set of regulated NCC uORFs, this set was also distributed into all four quadrants, with a preponderance in Quadrants 1 and 2.

We also performed this analysis for all translated AUG uORFs (Figure 8E, F). Again, the plot shows a distribution of mRNAs into all four quadrants. However, consistent with the behavior of AUG uORFs described above, a majority (66%) are in Quadrants 1 and 4 (i.e., negative ΔTE_uORF_) at 20 °C, whereas at 37 °C a majority (76%) are in Quadrants 2 and 3 (i.e., positive ΔTE_uORF_).

Why TE_mORF_ increases for the mRNAs described in Figure 8A-F more often than it decreases regardless of the direction of ΔTE_uORF_ is unclear, although it is possible that mRNAs whose main ORF translation decreases (negative ΔTE_mORF_) are more likely to be degraded due to the coupling between active translation and mRNA stability [55, 56] and thus less likely to appear in the ribo- or RNA-seq data.

In order to assess possible trends in the relative translation of the uORFs and mORFs, we colored the circles corresponding to each uORF in the plots in Figure 8 according to their relative ribosome occupancies (RRO_uORF_) values at 30 °C: tan for an RRO < 0.1, light brown for an RRO between 0.1 and 0.5, and dark brown for an RRO > 0.5. In general, no obvious trends in RROs emerge for mRNAs with temperature-regulated uORFs. However, the plots for AUG uORFs (Figure 8E, F) make clear that most AUG uORFs are well-translated relative to their downstream uORFs (dark brown circles, RRO > 0.5), in contrast to the situation with NCC uORFs (Figure 8A-D), where a wide range of RROs is observed and a majority are ≤ 0.5.

### 3.10 uORF-dependent regulation of translation by changes in growth temperature

Figure 8 highlights a number of clear cases in which uORF TE decreases and the TE of the downstream mORF correspondingly increases (labeled circles in Quadrant 1) or the uORF TE increases and the TE of the downstream mORF decreases (labeled circles in Quadrant 3). One very well-established example of uORF-dependent translational regulation is the *GCN4* mRNA in which four AUG uORFs are involved in controlling nutrient-dependent modulation of the translation of the mORF [57]. Recently, ribosome profiling studies indicated that there was also a translated uORF starting with an NCC (AUA) upstream of the canonical AUG uORFs [58], and that its TE was increased upon amino acid starvation in yeast [8], and under sustained histidine limitation [58]. However, evidence was presented that increased or decreased translation of this NCC uORF is not associated with a significant change in *GCN4* expression under non-starvation, starvation or stress conditions at 30 °C [58].

Intriguingly, the *GCN4* NCC uORFs have the strongest reciprocal increases in mORF translation when their translation is repressed at 37 °C of any NCC uORFs (Figure 8B, D, Quadrant 1). We therefore examined the ribo and mRNA read density plots (“wiggle tracks”) from the 20, 30 and 37 °C data (Figure 9A, Additional file 1: Figure S10A). At all three temperatures, very strong ribosome footprint density was observed for AUG uORFs 1 and 3 with clear but lower density at AUG uORFs 2 and 4. We also observed significant ribosome occupancy at several NCC uORFs upstream of the canonical AUG uORFs, including the AUA uORF reported previously [58] (Figure 9A, left, and bottom panel, Additional file 1: Figure S10A). At 20 °C the TEs of these NCC uORFs were increased on average 1.6-fold relative to 30 °C, which had no detectable effect on the TE of the mORF (Figure 9A, compare black and blue traces, effective increase in RRO_uORF_ ∼2-fold). In contrast, at 37 °C the TEs of the NCC uORFs decreased ∼3-fold, which was accompanied by an ∼2-fold increase in mORF translation (effective decrease in RRO_uORF_ ∼6-fold), suggesting that the NCC uORFs may be involved in regulating translation of the *GCN4* mORF at elevated temperatures. Further, we calculated the TEs of the AUG uORFs manually as our pipeline does not allow the validation of uORFs shorter than three codons. We found that these uORFs also show temperature dependent changes in TEs. At 20 °C, the TE of uORF1 is decreased ∼1.7-fold relative to 30 °C, while the TE of uORF3 is increased by ∼1.5-fold. At 37 °C, the TE of uORF2 is decreased ∼3-fold relative to 30 °C, while the TEs of uORFs 3 and 4 are increased ∼1.7- and 2.7-fold, respectively. The differing behavior of each *GCN4* uORF with respect to temperature seems to underscore the conclusion that no single variable is solely responsible for the observed effects of growth temperature on uORF translation.

**Figure 9.**
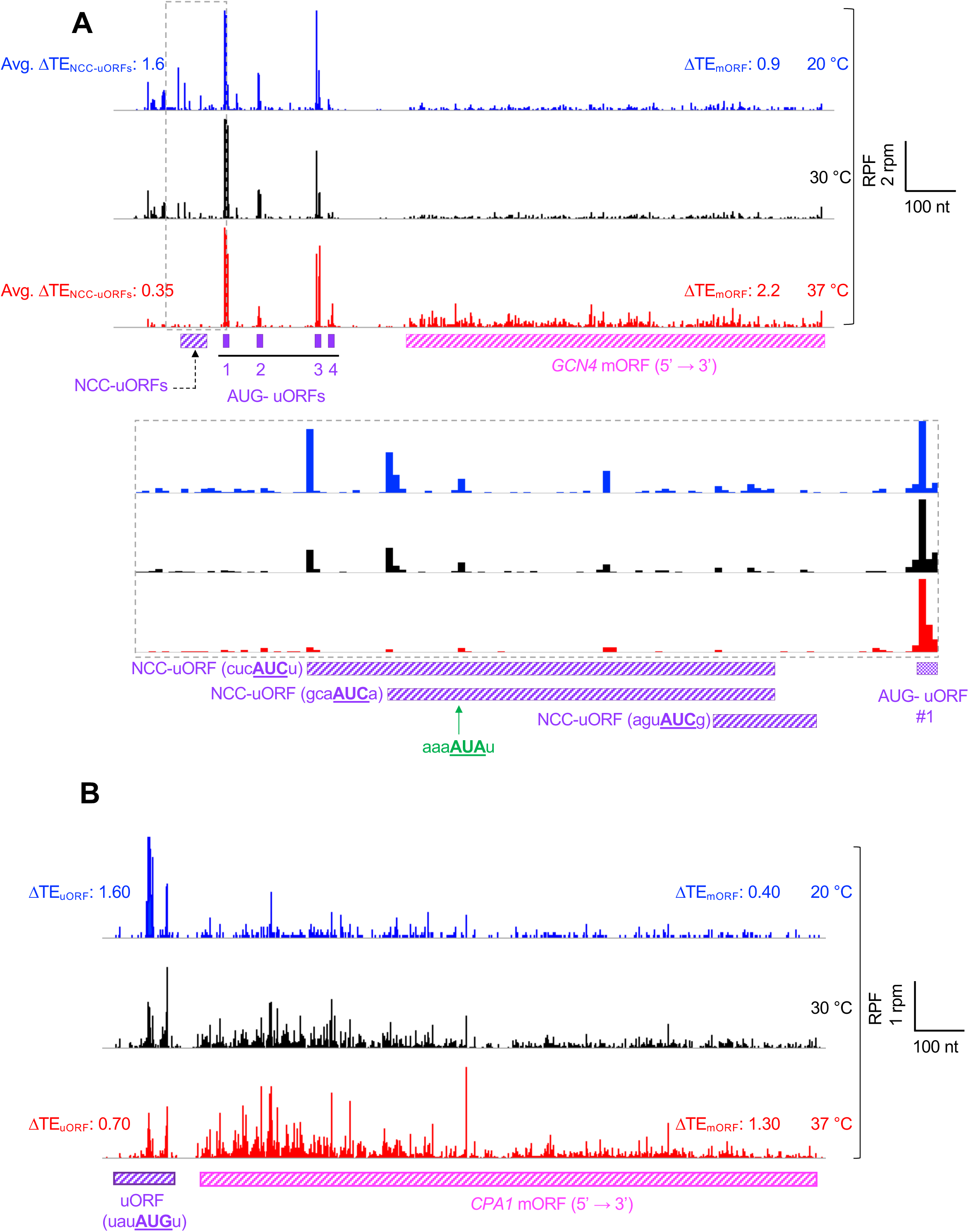
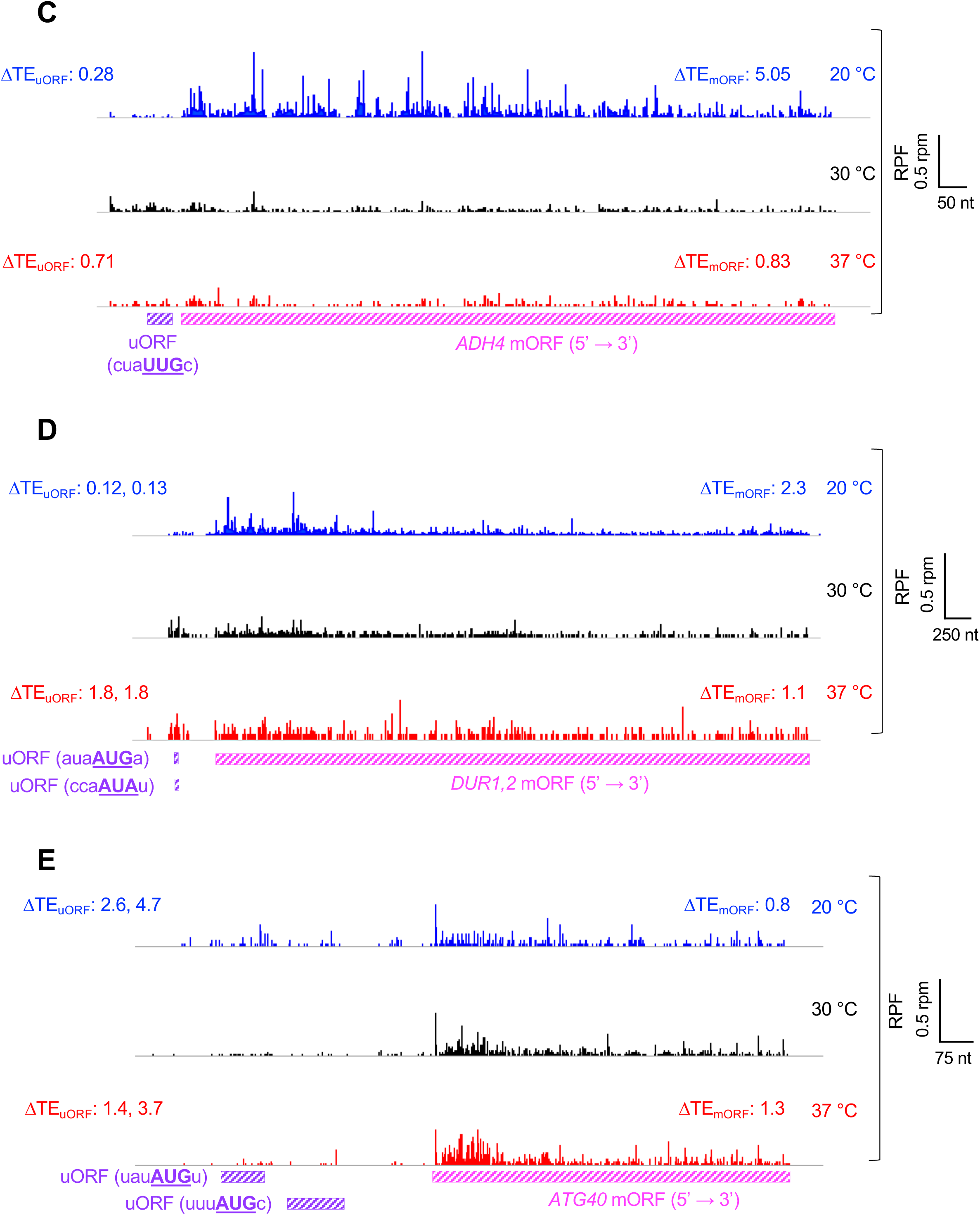
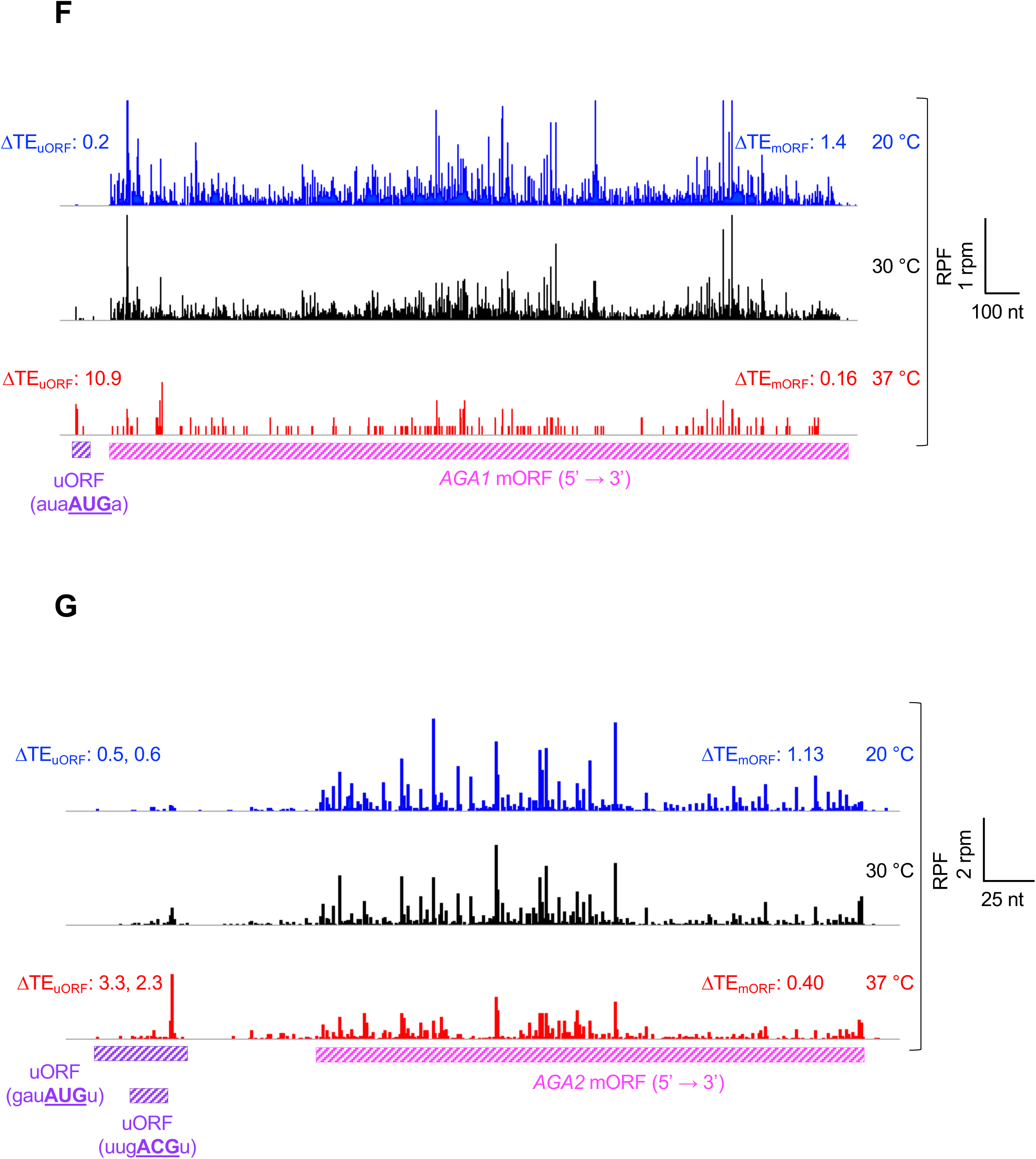
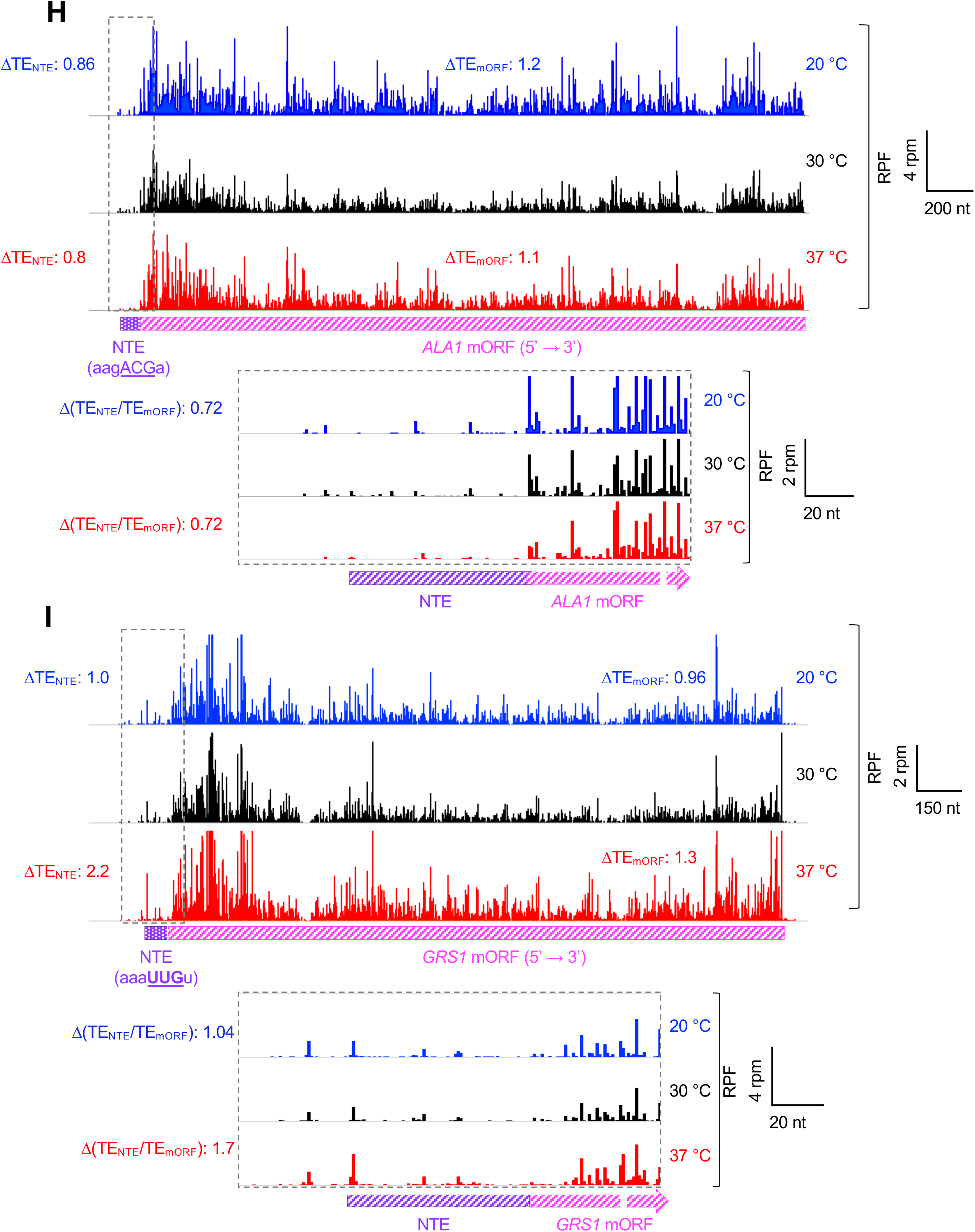
mRNAs that show reciprocal changes in the translation of uORFs and mORFs at multiple temperatures. **(A)** Wiggle track images showing ribosome-protected fragments (RPF) on the *GCN4* mRNA in cells cultured at either 20, 30 or 37 °C, in units of rpm (reads per million mapped reads from two replicates at each temperature). The RPF-tracks were normalized to the mRNA levels at each temperature to reflect the changes in translation efficiencies (ΔTE) of uORF and mORF as described in the legend to Figure 2 and Methods. The schematic shows the position of the uORFs (purple rectangles) and mORF (striped pink rectangle). NCC uORFs are shown with striped purple rectangle. AUG uORFs are shown with filled rectangles in purple. Average change in the translation efficiencies of the three NCC uORFs showing significant changes in translation at 20 or 37 °C (Avg. ΔTE_NCC uORFs_) with respect to 30 °C is shown for both 20 °C (blue) and 37 °C (red). ΔTE_mORF_ represents the change in translation efficiency of the mORF at 20 °C (blue) and 37 °C (red) with respect to 30 °C. The enlargement of the boxed area is also shown below. Start sites of NCC uORFs (bold, underlined) and the −3 to −1 and +4 context nucleotides are shown in parentheses. The green arrow shows the uORF start site (AUA; shown in green, bold underlined letters) that has been previously shown to be used as an upstream start site [58]. **(B-G)** Same as in **(A)** but for (B) the *CPA1* mRNA; **(C)** *ADH4* mRNA; (**D**) *Dur1,2* mRNA; (**E**) *ATG40* mRNA; (**F**) *AGA1* mRNA; **(G)** *AGA2* mRNA. **(H, I)** Wiggle track images of the *ALA1* **(H)** and *GRS1* **(I)** mRNAs in cells cultured at either 20, 30 or 37 °C, as described in **A**. The schematic shows the position of the N-terminal extension (striped purple rectangle) and mORF (striped pink rectangle). Changes in translation efficiencies of NTE and mORF (ΔTE_NTE_ and ΔTE_mORF_, respectively) with respect to 30 °C are shown for both 20 °C (blue) and 37 °C (red). The enlargement of the boxed area is also shown. Relative TE_NTD_ (TE_NTE_/TE_mORF_ ratio) reflects the ratio of translation efficiency of initiation at the upstream start site of the NTE (ACG in the case of *ALA1* and UUG in the case of *GRS1*) to that of the combined initiation events at the upstream start site and mORF start site (AUG in both cases).

Another well-studied example of uORF-mediated translational regulation is the *CPA1* mRNA, which encodes an arginine biosynthetic enzyme. At high arginine levels, translation of the AUG uORF represses translation of the downstream mORF [59–61]. Strikingly, in our data the *CPA1* mRNA has the highest degree of negative regulation of mORF translation of any mRNA with an AUG uORF when the translation of the uORF increases at 20 °C relative to 30 °C (Figure 8E, Quadrant 3). At 20 °C TE_uORF_ was elevated ∼1.6-fold while TE_mORF_ was repressed ∼2.5-fold (effective increase in RRO_uORF_ ∼4-fold) (Figure 9B, Additional file 1: Figure S10B, compare black and blue traces). Conversely, at 37 °C TE_uORF_ was repressed ∼1.4-fold and TE_mORF_ was elevated by 1.3-fold (effective decrease in RRO_uORF_ ∼2-fold) (Figure 9B, compare black and red traces). These data suggest that *CPA1* translation may be regulated by temperature-dependent changes in uORF translation, either through a direct effect of temperature on uORF TE or indirectly, for example due to temperature-dependent changes in arginine levels in the cell.

A number of other mRNAs that were not previously known to be subject to uORF-mediated translational control appear in our data as having reciprocal, temperature-dependent changes in uORF and mORF TE values suggestive of a regulatory relationship (Figure 8). For example, *ADH4* mRNA, which encodes an alcohol dehydrogenase enzyme, undergoes a 5-fold increase in mORF TE at 20 °C while the TE of its NCC uORF decreases 3.6-fold (effective decrease in RRO_uORF_ ∼18-fold) (Figure 8A, C; Figure 9C, Additional file 1: Figure S10C, compare black and blue traces). Likewise, the *DUR1,2* mRNA, which encodes a urea amidolyase enzyme, has one NCC uORF and one AUG uORF. The TEs of these uORFs decrease nearly 10-fold at 20 °C with a corresponding increase of 2.3-fold in the TE of the mORF (effective increase in RRO_uORF_ ∼20-fold) (Figure 8A,C; Figure 9D and Additional file 1: Figure S10D, compare black to blue).

The *AGA1* and *AGA2* mRNAs, which encode the subunits of the a-agglutinin receptor, are particularly striking examples of possible uORF-dependent translational regulation (Figure 8A, B, D-F, Quadrants 1 and 3). Translation of the *AGA1* AUG uORF increases over 10-fold at 37 °C relative to 30 °C, which is accompanied by a 6-fold decrease in TE for the mORF (effective increase in RRO_uORF_ ∼60-fold) (Figure 9F and Additional file 1: Figure S10F, compare black and red traces). TE of the uORF is also decreased 5-fold at 20 °C, with a corresponding increase in mORF TE of 1.4-fold (effective decrease in RRO_uORF_ ∼7-fold). The fact that the increase in TE_mORF_ is smaller than the decrease in TE_uORF_ can be explained by the fact that at 30 °C the uORF is translated at only a small percentage of the level of the main ORF, with an RRO value of 0.18. *AGA2* mRNA, which has two translated uORFs, displays similar behavior (Figure 9G and Additional file 1: Figure S10G). Intriguingly, the constitutive agglutinability of yeast cells, the process mediated by a-agglutinin, has been reported to be diminished at growth above 30 °C [62], consistent with the putative uORF-mediated regulation of *AGA1* and *AGA2* expression suggested by our ribosome profiling data.

In addition to altering uORF translation, changes in the efficiency of initiation site usage can alter the levels of inclusion or exclusion of N-terminal extensions (NTEs) that are in-frame with the main ORF of an mRNA, changing the balance between different protein isoforms. We found 130 cases where initiation upstream of the annotated mAUG appears to lead to formation of an NTE (Additional file 2: Supplementary Table 4). Interestingly, >95% of these upstream start sites are NCCs. Two well-characterized examples of N-terminal extensions that encode mitochondrial localization signals are the yeast alanyl- and glutamyl-tRNA synthetases encoded by the *ALA1* and *GRS1* mRNAs, respectively [5, 6]. Both of these extensions initiate with NCCs, ACG for *ALA1* and UUG for *GRS1*. We see evidence of translation of both NTEs in our ribosome profiling data (Figure 9H, I and Additional file 1: Figure S10H, I), with relative ribosome occupancies for the NTE vs. mORF of 0.1 and 0.23, respectively at 30 °C. Although the TE of the NTE of *ALA1* is only modestly affected by growth temperature (Figure 9H and Additional file 1: Figure S10H) with an effective decrease in RRO_uORF_ ∼1.4-fold at both 20 °C and 37 °C, the TE of the *GRS1* NTE increases over 2-fold at 37 °C relative to 30 °C (Figure 9I and Additional file 1: Figure S10I, compare black and red traces, effective increase in RRO_uORF_ ∼1.7-fold). This change could potentially lead to an increase in the mitochondrial concentration of the glutamyl-tRNA synthetase at elevated growth temperatures.

## Discussion

In this report, we provide evidence for growth temperature-induced changes in the efficiency of translation of a subset of uORFs in the 5’-UTRs of mRNAs in *S. cerevisiae*. Using a multi-filter pipeline, we identified 1367 uORFs with strong evidence of translation at one or more growth temperatures in our ribosome profiling datasets. Most (90%) of the translated uORFs begin with near-cognate codons (NCCs), over half of which are 1^st^ position changes from AUG (UUG, GUG, CUG). The 10% that begin with AUG codons have higher TEs at the optimum growth temperature of 30 °C than those beginning with NCCs, as expected, and their downstream main ORFs have correspondingly lower TEs than the mORFs downstream of NCC uORFs. Moreover, the ribosome occupancies of the uORFs relative to the mORFs (relative ribosome occupancy; RRO) were higher for the AUG uORFs than for the NCC uORFs (median value of 1.8 and 0.17, respectively). These findings suggest that translation of AUG uORFs (at 30 °C) frequently reduces the fraction of scanning ribosomes that reach the mORF start codon, consistent with the usual inhibitory effect of uORF translation (for review see [13]), whereas the majority of NCC uORFs are translated at levels too low to exert this regulatory function.

Of the 1367 translated uORFs in our data set, we found ∼10% exhibited changes (activation or repression) in their translation at 20 and/or 37 °C relative to 30 °C that met our dual criteria for temperature-regulated uORF translation (Figure 4A and 4B, panel (iii)). A majority of the regulated uORFs that begin with AUG codons have reduced translational efficiencies at 20 °C and increased efficiencies at 37 °C (Figure 5A-B & E). These trends were also evident for the entire set of translated AUG uORFs (Figure 5C-D & F). In contrast, regulated NCC uORFs do not display a consistent trend as their translation can be activated, repressed or unaffected at either temperature. One possible explanation for this difference might be that the rate-limiting step for initiation on most AUG uORFs, which are generally well translated relative to the NCC uORFs (Figure 3D) and reside on shorter, less structured 5’-UTRs (Additional file 1: Figure S6), has a temperature dependence such that its rate increases with growth temperature. NCC uORFs, which tend to be on longer, more structured 5’-UTRs (Additional file 1: Figure S6) might have different rate-limiting steps for initiation depending on a variety of factors (e.g., position in the 5’-UTR, structural features of the 5’-UTR, length of uORF, etc.) and these differing rate-limiting steps could result in differing temperature dependencies. It is noteworthy that most of the regulated NCC uORFs (70/98) are in Q3-Q4 (Figure 5E), indicating repression of TE at 37 °C despite no clear trend at 20 °C. One possible explanation is that the imperfect codon-anticodon helices formed by NCCs are destabilized at high temperature, whereas the perfect pairing with AUGs is stable enough to resist this effect.

The position of the uORF in the 5’-UTR exerts a significant influence on the direction and magnitude of the temperature dependence of translation of uORFs. Translation of uORFs that are closer to the 5’-cap than the average distance for all uORFs tends to be inhibited at 20 °C relative to 30 °C, whereas translation of uORFs that are farther from the cap than the average tends to be activated at 20 °C (Figure 6 and 7D). In addition, for NCC uORFs, the distance from the mORF AUG codon also correlates with the temperature dependence of translation such that those farther from the mORF AUG are more likely to be activated at 37 °C (Figure 6 and 7E).

One simple explanation for some of the observed effects could be that low temperature stabilizes structure in 5’-UTRs, which is generally inhibitory towards uORF initiation, whereas higher temperature tends to destabilize overall 5’-UTR structure. Such an effect could explain uORFs whose translation decreases at 20 °C and increases at 37 °C, including the majority of AUG uORFs. On the other hand, in particular cases, stabilization of mRNA structures at low temperature and destabilization at higher temperatures could have the opposite effects if the structures are located downstream from the uORF start codon and cause PICs to pause near the sub-optimal start codons, increasing the probability of initiation on them [63, 64]. This mechanism could account for the activation of uORF translation at 20 °C and its repression at 37 °C. If a structural element is already unstable at 30 °C, it could be the case that increasing temperature to 37 °C has no significant effect, and if a structural element is already stable enough to produce a maximal effect at 30 °C, decreasing temperature to 20 °C might produce no additional observable effect. Thus, it is possible to rationalize most of the classes of effects we see at decreased or elevated growth temperatures simply by invoking the influence of temperature on RNA structure.

Nonetheless, it is likely that the temperature-dependent effects on uORF translation we observe are not all the result of a single mechanism. Other possible influences might include the rate of scanning, as slower scanning could lead to increased initiation on suboptimal start codons; changes in the thermodynamics of codon:anticodon pairings; alterations of the levels or activities of specific mRNA binding proteins or other factors; or the temperature-dependence of required structural rearrangements within the PIC or of enzymatic reactions such as GTP hydrolysis by eIF2 or eIF5B. Other factors such as the position of the uORF in the 5’-UTR, and hence the distance the PIC must scan to reach it, could further influence most of these possible mechanisms of regulation, leading to some of the observed correlations.

Although our data indicate that translation of a significant number of uORFs is regulated by growth temperature and suggest some cases in which these effects influence expression of the main ORF in the mRNA, it is also striking that this is not a general effect and that translation of most uORFs is relatively insensitive to changes in temperature, at least between 20 and 37 °C. Only 8% of the translated uORFs we identified (112/1359) had changes in TE that met our criteria for temperature-regulated translation, showing altered uORF translation without a similar increase or decrease in translation of the downstream mORF. Translation of the large majority of uORFs was refractory to temperature, which is remarkable in that temperature exerts effects on most reactions and interactions and has been shown to influence transcription, metabolism and overall cellular physiology in yeast [65, 66]. One possible explanation for this seeming conundrum could be that yeast cells evolved homeostatic mechanisms to damp down the influence of growth temperature on general translation in order to maintain appropriate levels of protein products. Because read numbers and TEs in each ribosome profiling experiment reflect values relative to the average observed value for the parameter in that experiment, changes reported are also relative to the population averages. Thus, it is possible that the absolute rates of translation are changing as a function of temperature for most uORFs, but in a linear fashion such that we do not observe changes relative to the population averages in each ribosome profiling experiment. Those uORFs for which we observe significant changes in TE are ones in which translation is increasing or decreasing more than the average change in the experiment. Nonetheless, our data indicate that translation of most uORFs behaves the same with respect to temperature, which implies a general mechanism to prevent relative translation rates of different ORFs from diverging when the growth temperature shifts and thereby changing global proteomic ratios in suboptimal ways.

More studies will be required to understand the mechanistic basis of the temperature-dependent regulation of uORF translation described here and of the physiological consequences of these phenomena. The observed effects of temperature on the translation of N-terminal extensions also warrants additional analysis because protein isoforms with different N-termini can have altered cellular localization patterns, functions or activities [5, 6, 67]. The set of temperature-dependent uORFs and N-terminal extensions we have identified using transcriptome-wide approaches should serve as a useful starting point for in-depth studies to elucidate the roles and underlying mechanisms of these intriguing systems.

## Declarations

### Ethics approval and consent to participate

Not applicable

### Consent for publication

Not applicable

### Availability of data and material

The ribo-seq data generated in this study have been submitted to the NCBI Gene Expression Omnibus and the accession numbers are listed https://elifesciences.org/articles/31250/figures#supp1 in the additional file table S2. Submission of the RNA-seq data from this study to NCBI Gene Expression Omnibus is pending.

### Competing interests

None declared.

### Funding

This work was supported by the Intramural Research Program of the National Institutes of Health (AGH and JRL). The funders had no role in the study design, data collection and interpretation, or the decision to submit the work for publication.

### Authors’ contributions

**Shardul D. Kulkarni:** Conceptualization, Resources, Data curation, Formal analysis, Investigation, Visualization, Methodology, Writing—original draft, Writing—review and editing

**Fujun Zhou:** Resources, Data curation, Formal analysis, Methodology.

**Neelam Dabas Sen:** Resources, Data curation, Formal analysis, Methodology.

**Hongen Zhang:** Methodology

**Alan G. Hinnebusch**: Conceptualization, Formal analysis, Supervision, Funding acquisition, Visualization, Project administration, Writing—review and editing

**Jon R. Lorsch**: Conceptualization, Formal analysis, Supervision, Funding acquisition, Visualization, Project administration, Writing—review and editing

## Acknowledgements

We thank Thomas Dever and Nicholas Guydosh for thoughtful comments and suggestions. We are grateful to Nick Ingolia for advice in the early phases of this project. We would also like to thank all the members of Lorsch, Hinnebusch, Dever and Guydosh labs for their useful comments.

## Additional file 1: Supplementary methods, Supplementary Figures and legends

### Supplementary Methods

#### The 5’-UTRs of mRNAs with uORFs tend to be longer and more structured than the genomic average

To determine whether the general characteristics of the 5’-UTRs of mRNAs with uORFs are different than those of mRNAs that don’t have evidence of translated uORFs, we examined their lengths and propensities to form secondary structures. We interrogated the dataset of ∼2700 yeast mRNA 5’-UTR lengths and propensities of forming secondary structures [1]. As described previously, we identified 1367 uORFs (NCC and AUG, Figure 3A) which were present on mRNAs (*uORF mRNAs*) with an average 5’-UTR length of 195 nt, which was significantly greater than the average 5’-UTR length of ∼79 nt calculated for all mRNAs (*All mRNAs*) in the PARS dataset (Additional file 1: Figure S6A). We also examined the mRNAs that do not show evidence of an actively translated uORF (*Non-uORF mRNAs*) as identified by our pipeline discussed in Figure 3A. These 2157 non-uORF mRNAs had an average 5’-UTR length of 66 nt, significantly shorter than the average for *All mRNAs*. Figure S6A (Additional file 1) shows the cumulative fraction distribution of these three sets of mRNAs.

To examine 5’-UTR secondary structure, we analyzed the PARS (Parallel Analysis of RNA Structure) data available from the same study [1]. In this study, each nucleotide in ∼ 3000 mRNAs was assigned a PARS score, which is a measure of its propensity to be in the double stranded conformation. The PARS score is based on the susceptibility of each nucleotide in the transcribed mRNAs to in vitro digestion with RNase V1 (specific to double-stranded mRNA) and RNase S1 (specific to single-stranded mRNA). Higher PARS score indicates a greater tendency of a nucleotide to be in a double-stranded conformation. We analyzed the PARS scores for each of the sets of mRNAs: *All mRNAs, uORF mRNAs and non-uORF mRNAs*. We considered several PARS features, each of which is an indication of an extent to which a region of the 5’-UTR or coding region near the 5’-UTR can form a secondary structure (Additional file 1: Figure S6B): sum of PARS scores of all the nucleotides (nt) present in the 5’-UTR (*Total*); average PARS score per nt (*Average*); sum of PARS scores for the first 30 nts (*First30*); sum of PARS scores for 30 nts surrounding the start codon (for mRNAs with a 5’-UTR of ≥15 nt; *Start30*); and highest total PARS score measured for a 30-nt region anywhere across the 5’-UTR (*Max30*). We also analyzed PARS scores downstream from the mAUG (start codon of the main ORF), with an interval of +1 to +30 (*Plus15*).

Interestingly, the *uORF mRNAs* were shown to have higher PARS scores than *All mRNAs* for most of the PARS features considered (Additional file 1: Figure S6C, compare red versus purple columns). Notably, the *Non-uORF mRNAs* showed significantly lower PARS scores for some of the PARS features compared to *All mRNAs* (Additional file 1: Figure S6C, compare red versus blue columns), suggesting that these *non-uORF mRNAs* are less structured than the genomic average. Together, these data indicate that mRNAs containing actively-translated uORFs typically have longer and more structured 5’-UTRs than those without uORFs.

### Supplementary Figures and Legends

**Supplementary Figure 1.**
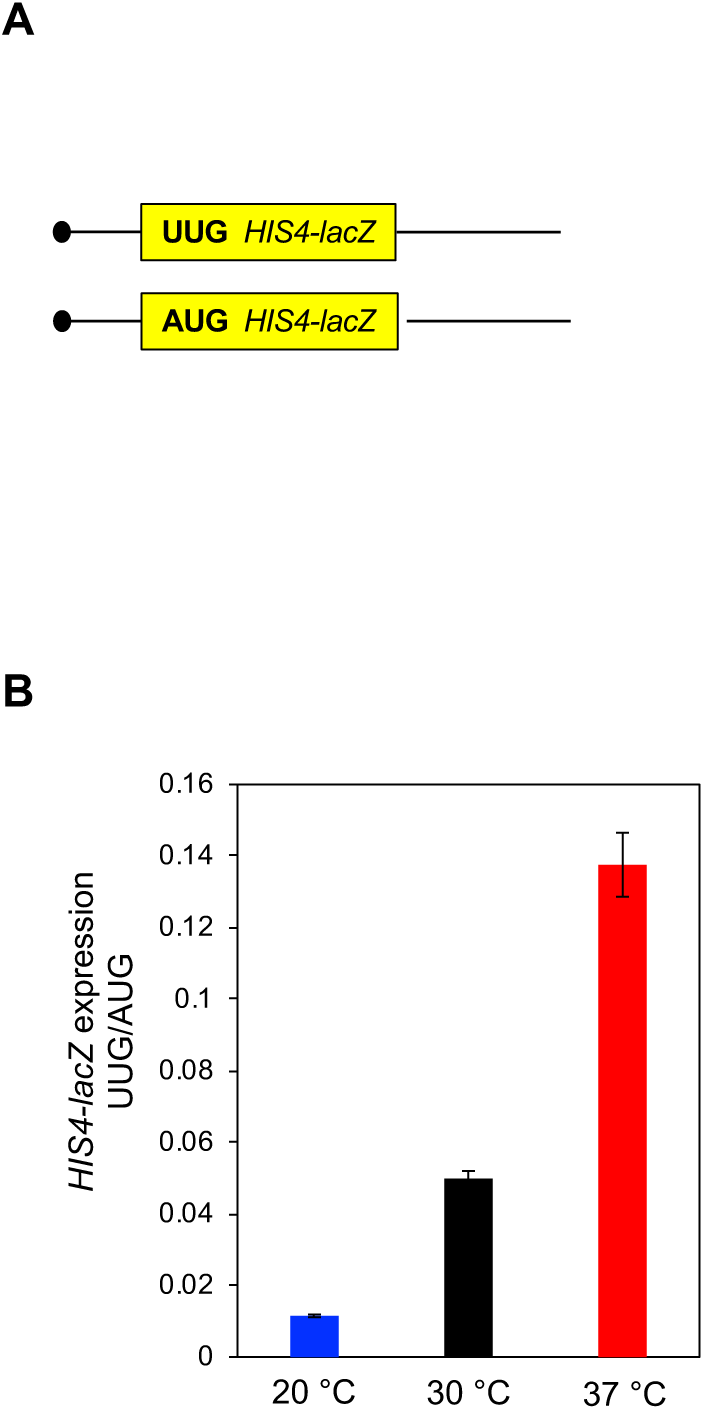
β-galactosidase activity assay to calculate usage of NCC start site at multiple growth temperature. **(A)** Schematic of *HIS4-lacZ* reporters used in this study. **(B)** BY4741 cells with *HIS4-lacZ* reporters with AUG or UUG start codons were cultured at 20 °C, 30 °C or 37 °C to an A_600_ of 0.6 to 0.8 and β-galactosidase specific activity was measured in total cell extracts in units of ONPG cleaved per minute per mg of total protein in the extract. Ratios of expression from the UUG reporter to the AUG reporter were calculated and plotted as averages from two independent biological replicates with average deviation as error bars.

**Supplementary Figure 2.**
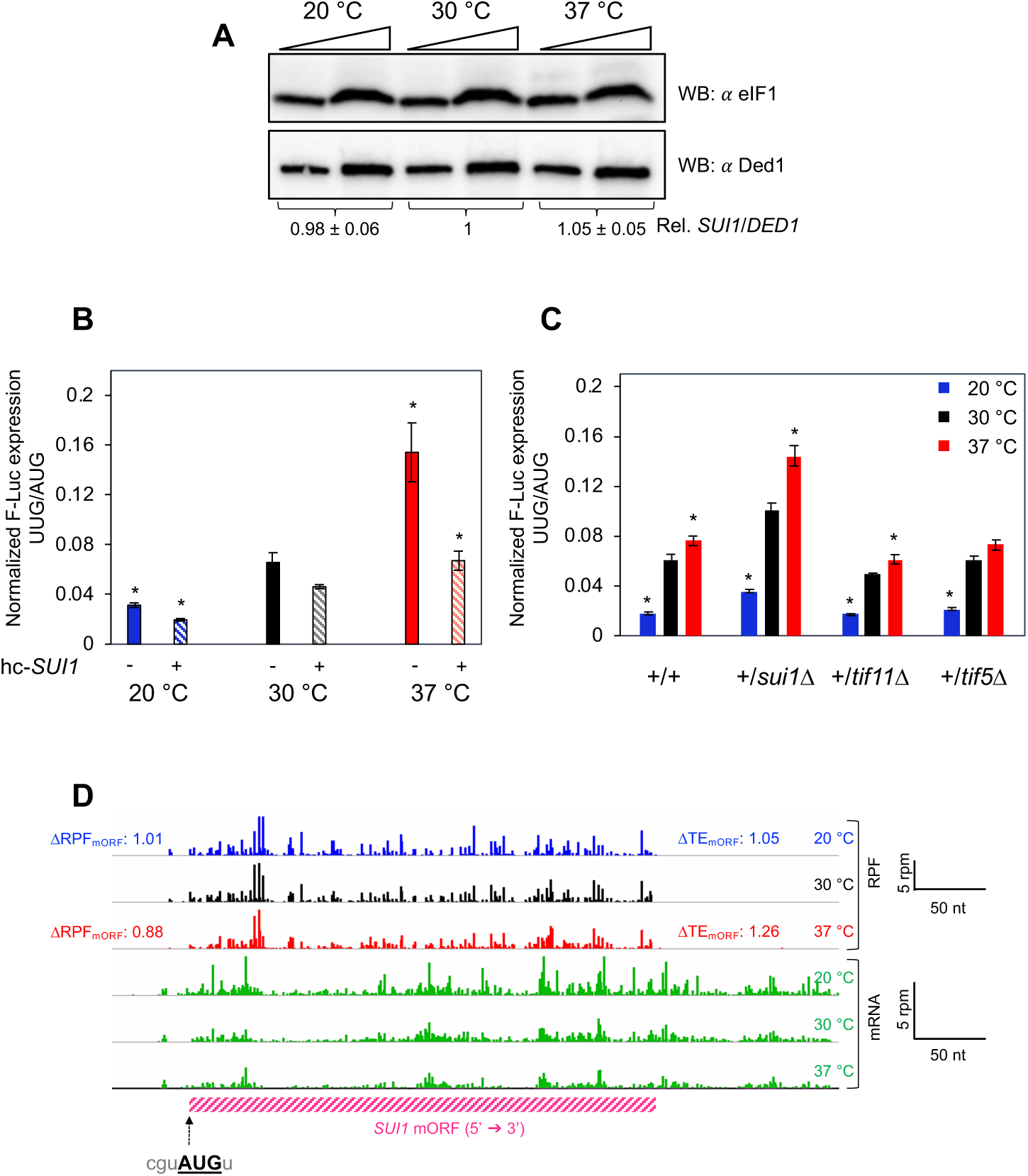
Temperature dependent alterations in usage of NCC start site are not evoked due to changes in the levels of initiation factors. (**A**) BY4741 cells harboring F-Luc reporter (UUG) were cultured at either 20 °C, 30 °C or 37 °C to an A_600_ of 0.6 to 0.8 and whole cell extracts were subjected to Western blot analysis using antibodies against eIF1 and Ded1 (loading control). Extracts were loaded in two amounts differing by a factor of two. eIF1 western blot signal was normalized to that of Ded1, to calculate relative *SUI1* levels, which was then set as 1 for 30 °C and the relative *SUI1* expression at 20 °C or 37 °C was then calculated. The values represent the average from 4 independent experiments with error bars indicating standard error of mean. (**B**) F-Luc expression (UUG/AUG ratio) was measured in a strain H3984 (hc-*SUI1*) and compared with wild-type (BY4741) at 20 °C, 30 °C or 37 °C. The data represent the average of 4 independent biological replicates with error bars representing standard error of mean. (**C**) F-Luc expression (UUG/AUG ratio) was measured at 20 °C, 30 °C or 37 °C in haplo-insufficient diploids for eIFs 1 (+/*sui1Δ*), 1A (+/*tif11Δ*), and 5 (+/*tif5Δ*), and compared with diploid wild-type BY4743 (+/+). The data represent the average of 4 independent biological replicates with error bars representing standard error of the mean. For B and C: * denotes p-values < 0.05 calculated by Student’s *t*-test when compared with 30 °C. (**D**) Ribosome-protected fragments (RPFs) and mRNA reads (RNA-seq) on the *SUI1* gene in cells cultured at either 20, 30 or 37 °C, in units of rpm (reads per million mapped reads). The RPF reads are shown in blue (20 °C), black (30 °C) or red (37 °C). The mRNA reads at each temperature are shown in green. The dashed arrow represents the start site of the mORF of the *SUI1* gene along with its context nucleotides. The changes observed in RPF counts and TE for the mORF of eIF1 are shown in blue (20 °C) and red (37 °C).

**Supplementary Figure 3.**
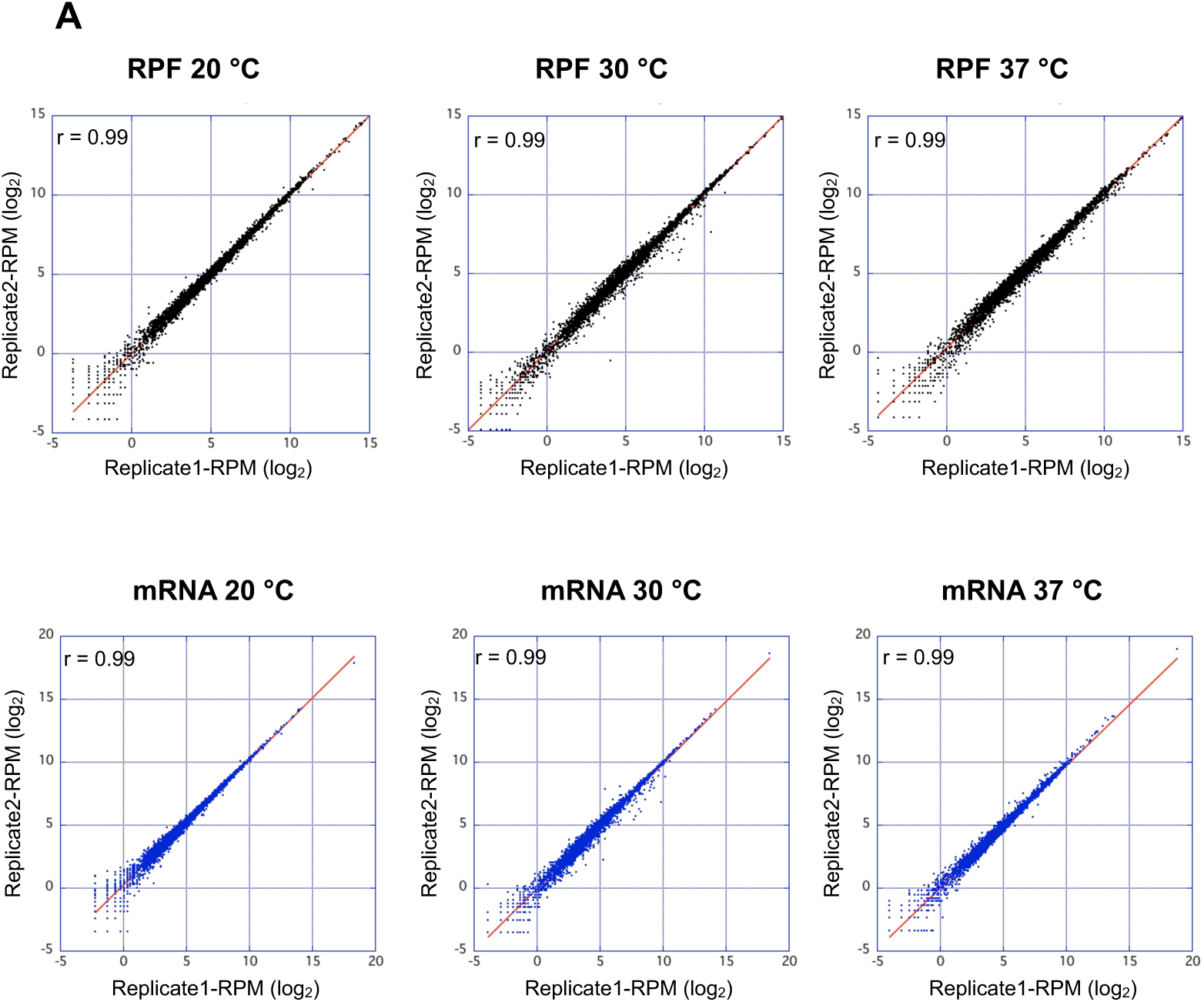
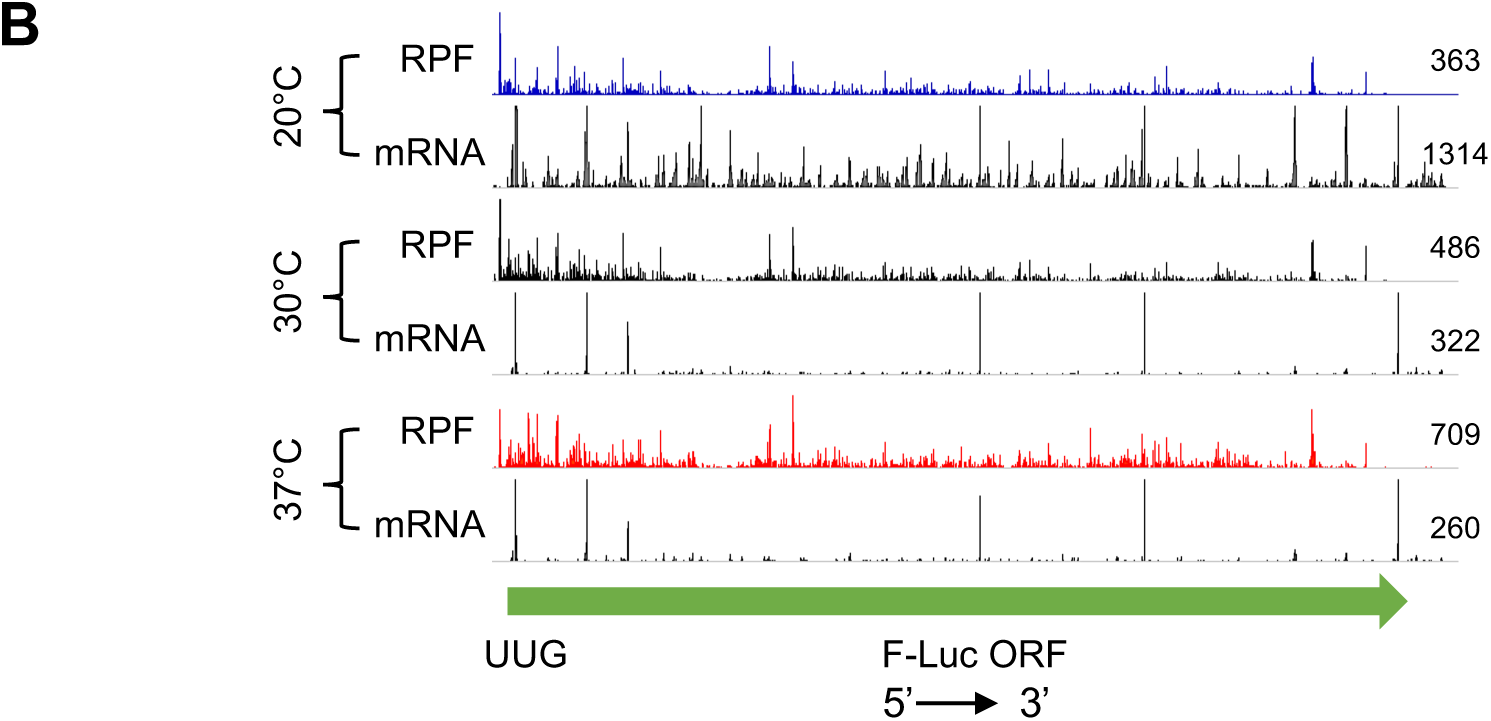
Ribosome profiling at multiple growth temperatures. **(A)** Data reproducibility of ribosomal footprint and mRNA densities at multiple temperatures. Scatterplots of ribosomal densities [ribosome protected fragments (RPF) reads] in top panels and mRNA densities (RNA-seq reads) in bottom panels at 20, 30 and 37 °C. The densities were represented by the reads mapped to the coding region of a gene per million mapped reads in the individual library of a biological replicate. The r values on each plot represent the Pearson correlation coefficients calculated for all genes (n∼5400). **(B)** Ribosome protected fragments (RPFs, from ribo-seq) and mRNA reads (RNA-seq reads) on the F-Luc^UUG^ reporter at multiple temperatures. RPF reads or mRNA reads (mapped to coding regions) were normalized to the total number of mapped reads at each temperature. RPF reads for 20, 30 and 37 °C are shown in blue, black and red, respectively, while the mRNA reads are shown in gray for all the temperatures. The numbers at the end of each wiggle track on the right indicate the average read densities (reads per million, RPM) of the two replicates. For the RNA density calculations, the presumed PCR jackpot peaks from the mRNA traces were removed.

**Supplementary Figure 4.**
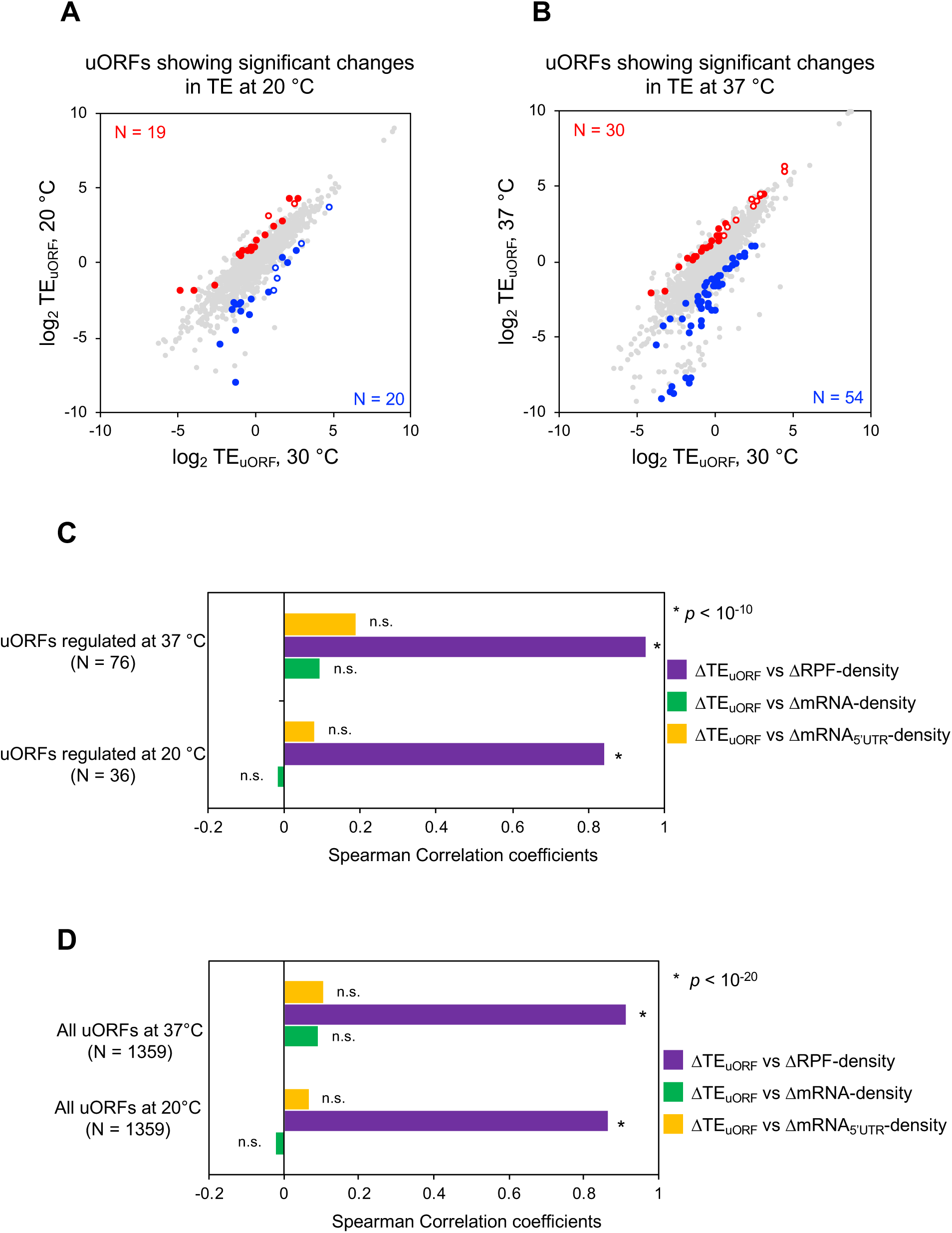
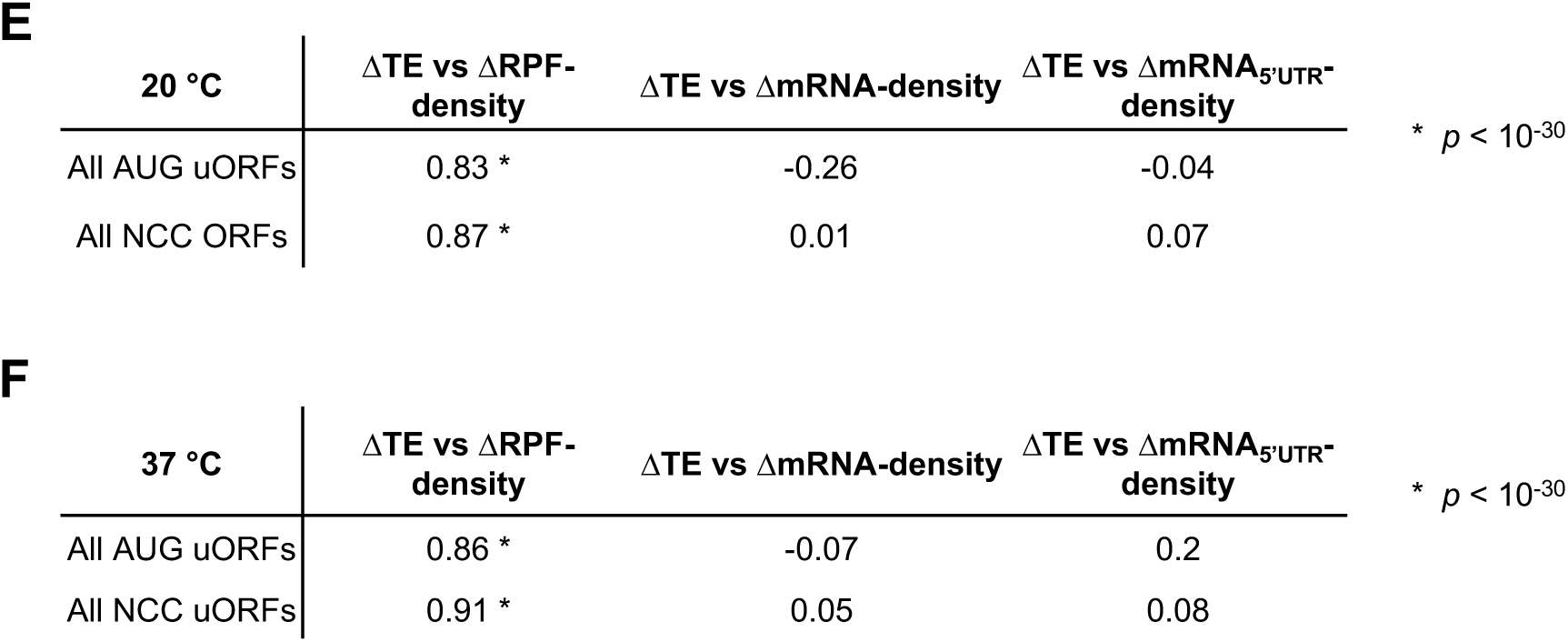
Changes in growth temperature lead to changes in translation of uORFs. **(A)** Scatterplot of TE_uORF_ for cells grown at 30 and 20 °C. The scatterplot shows uORFs exhibiting ≥ 2-fold changes (repression or activation) in TE_uORF_ and at a False Discovery Rate (FDR) < 0.1. Red circles represent such uORFs whose translation is activated at 20 °C as compared to 30 °C (N = 19), while blue circles represent uORFs whose translation is repressed at 20 °C as compared to 30 °C (N = 20). The open circles are AUG uORFs while the filled circles are NCC uORFs. This plot differs from the plots in Figure 4 A and B (iii) in that no filter was applied here to ensure that the TE of the uORF changes more than the TE of the mORF. **(B)** Same as in **A**, except; scatterplot of TE_uORF_ for cells grown at 30 and 37 °C. Red circles represent uORFs whose translation increases significantly at 37 °C as compared to 30 °C (N = 30), while blue circles represent uORFs whose translation decreases significantly at 37 °C as compared to 30 °C (N = 54). **(C-F)** Spearman correlation coefficient analysis between ΔTE_uORF_ versus Δribo-density_uORF_, ΔTE_uORF_ versus ΔmRNA-density_mORF_ and ΔTE_uORF_ versus ΔmRNA-density_5’-UTR_ for the all translated uORFs and regulated uORFs at 20 and 37 °C. Cases in which the two-sided p-value is less than 10^−10^ (C), 10^−20^ (D) or 10^−30^ (E, F) are denoted with an asterisk. A high Spearman’s coefficient value indicates a strong and directional monotonic relationship between the two variables tested. **(C**) uORFs showing temperature-dependent regulation of translation [as in Figures 4A and 4B, highlighted in red or blue in panel (iii)] are analyzed here. uORFs whose translation was either activated or repressed at 20 °C (N = 36) or at 37 °C (N = 76) were used to calculate the Spearman’s correlation coefficients. **(D**) Same as in **C**; except with all translated uORFs (N = 1359). **(E, F)** Same as in (D) except all translated uORFs were binned into uAUGs (N = 136) and NCCs (N = 1223) and analyzed separately.

**Supplementary Figure 5.**
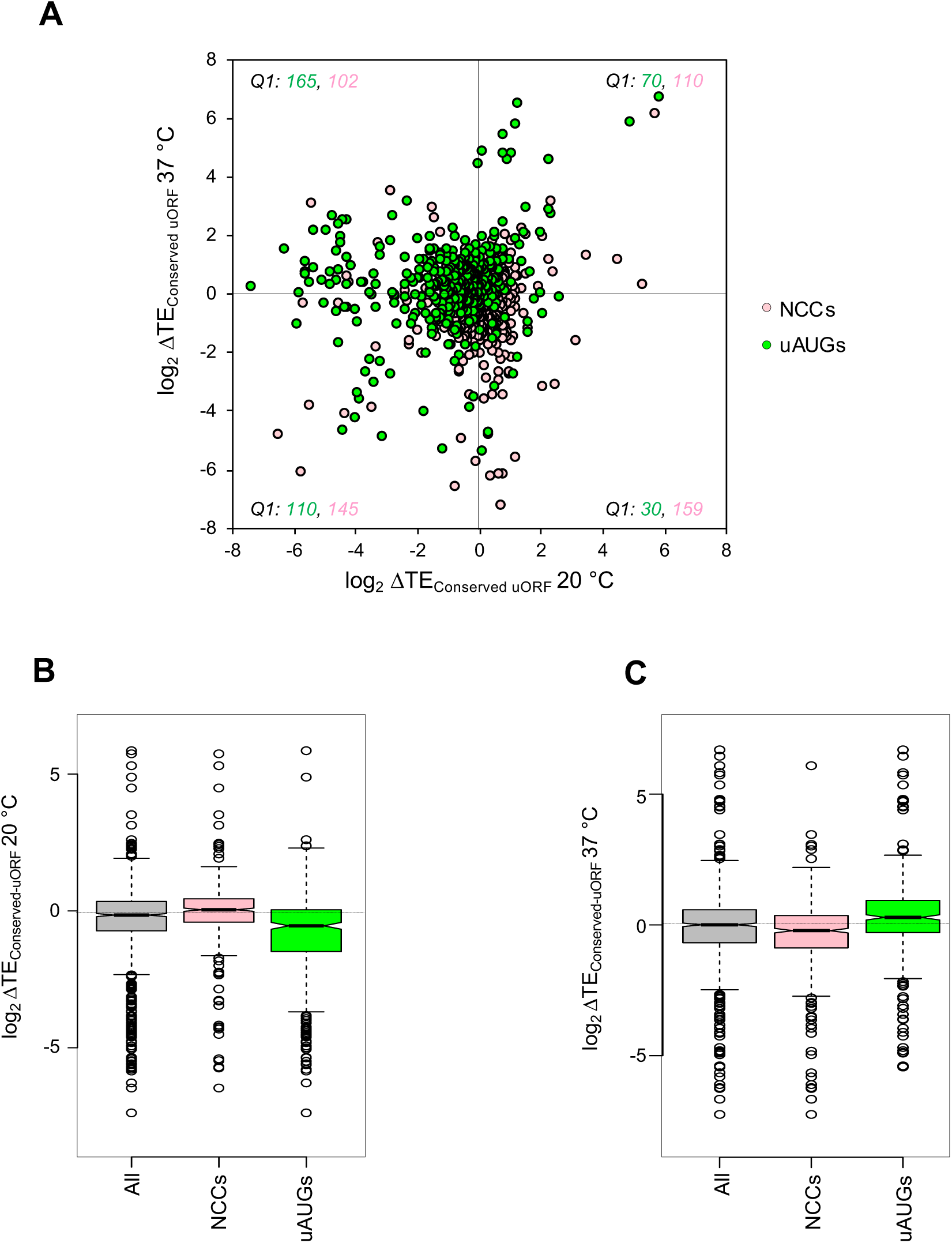
Analysis of translation efficiency changes of conserved uORFs upon changes in growth temperature. **A)** Translated uORFs shown to be conserved previously [2] were used here for scatterplot analysis between changes in TEs at 20 °C (ΔTE_Conserved-uORF_ 20 °C) and 37 °C (ΔTE_Conserved-uORF_ 37 °C). The plot is divided into four quadrants (Q1 to Q4) based on the directionality of TE changes. The numbers in each quadrant represent the number of uORFs present in that quadrant; shown in pink for NCC uORFs and in green for AUG uORFs. **(B, C)** Boxplot analysis of ΔTE_Conserved-uORF_ for the conserved uORFs analyzed in Additional file 1: Figure S5A). *All* : 891 conserved uORFs whose ΔTE values could be calculated using the dataset generated in this study; *NCCs*: 516 conserved NCC uORFs; *uAUGs*: 375 conserved AUG uORFs. The dotted horizontal line represents the median ΔTE for ‘*All*’. **(B)** Analysis done for TE changes at 20 °C with respect to 30 °C. **(C)** Analysis done for TE changes at 37 °C with respect to 30 °C.

**Supplementary Figure 6.**
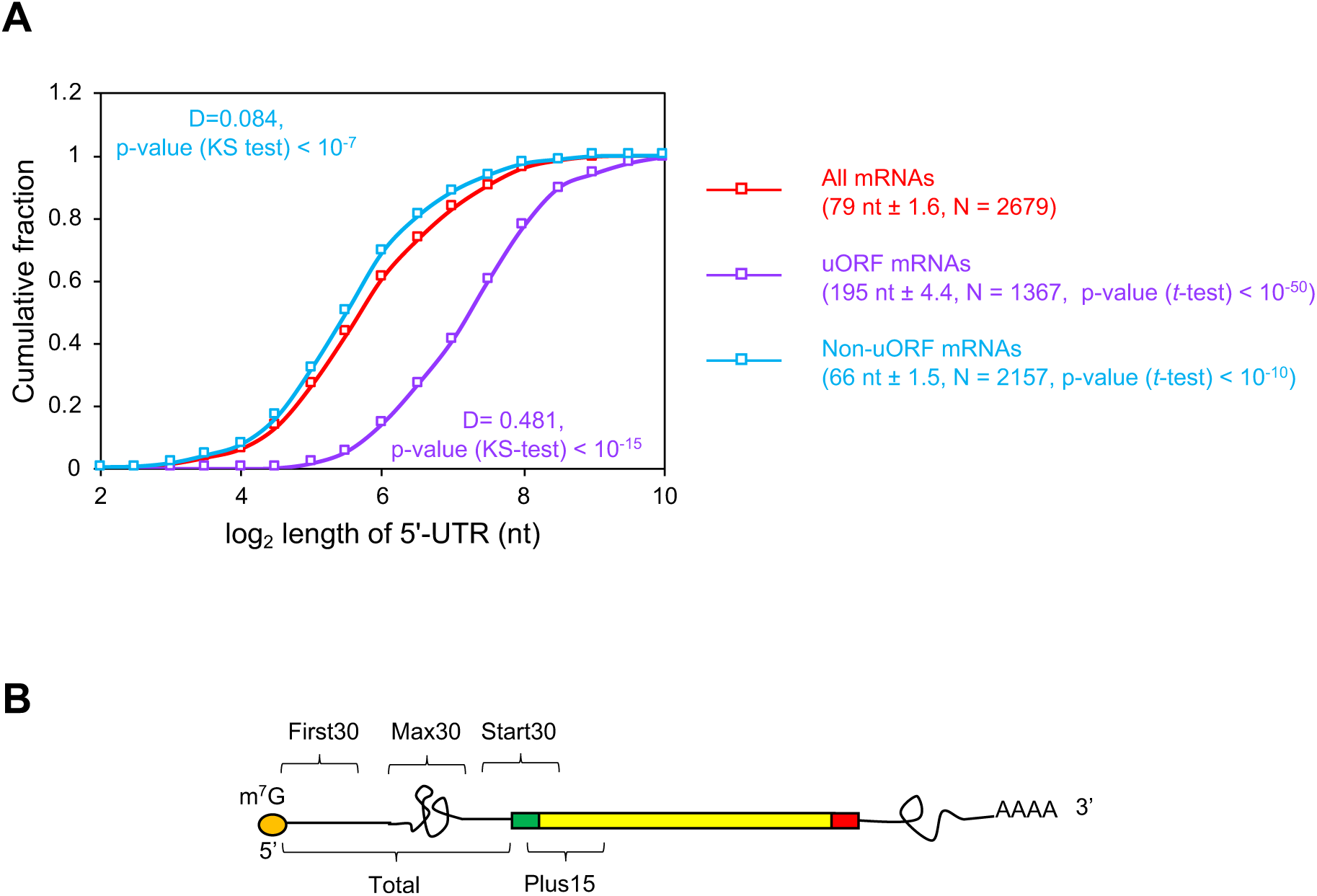
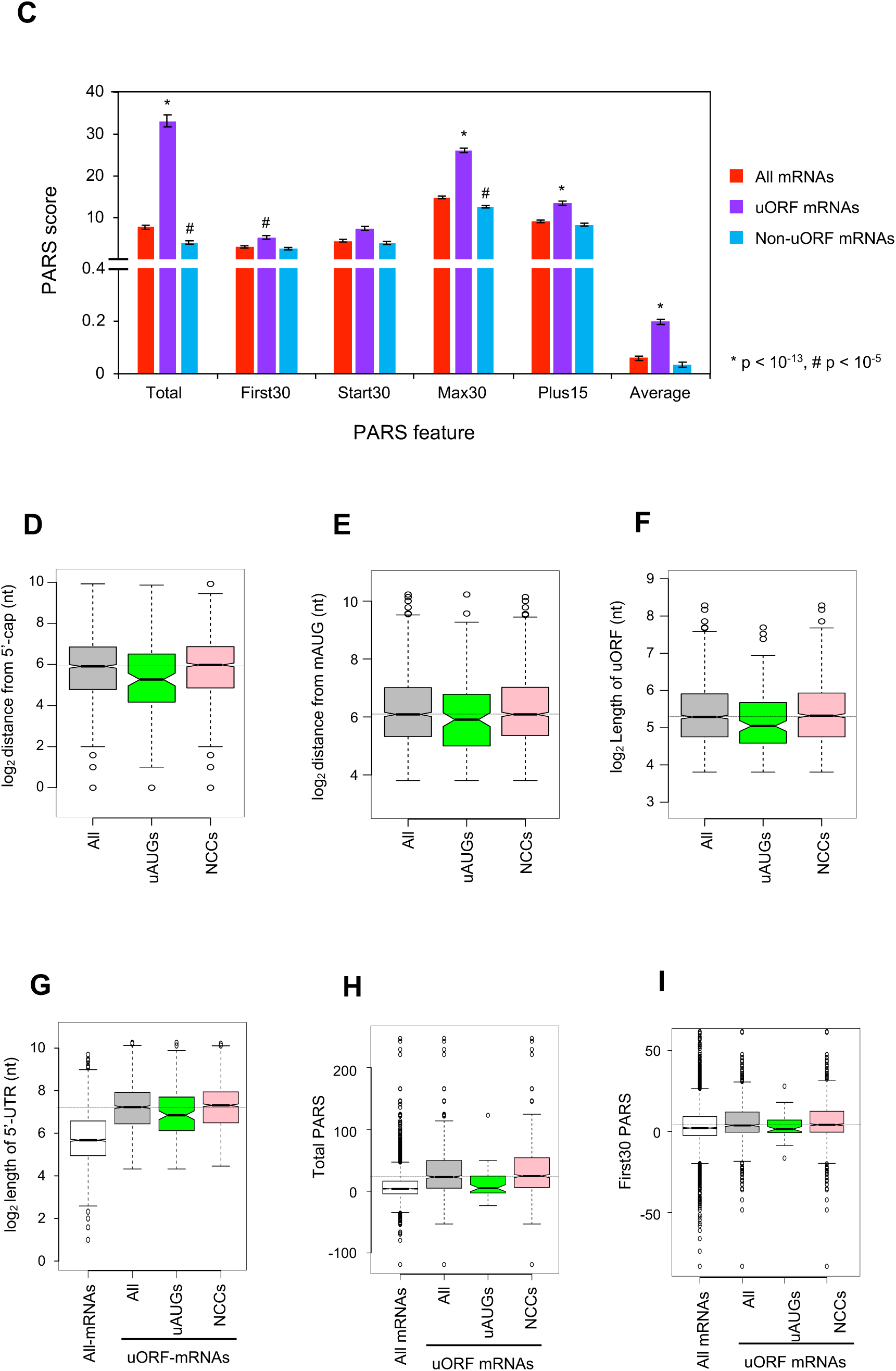
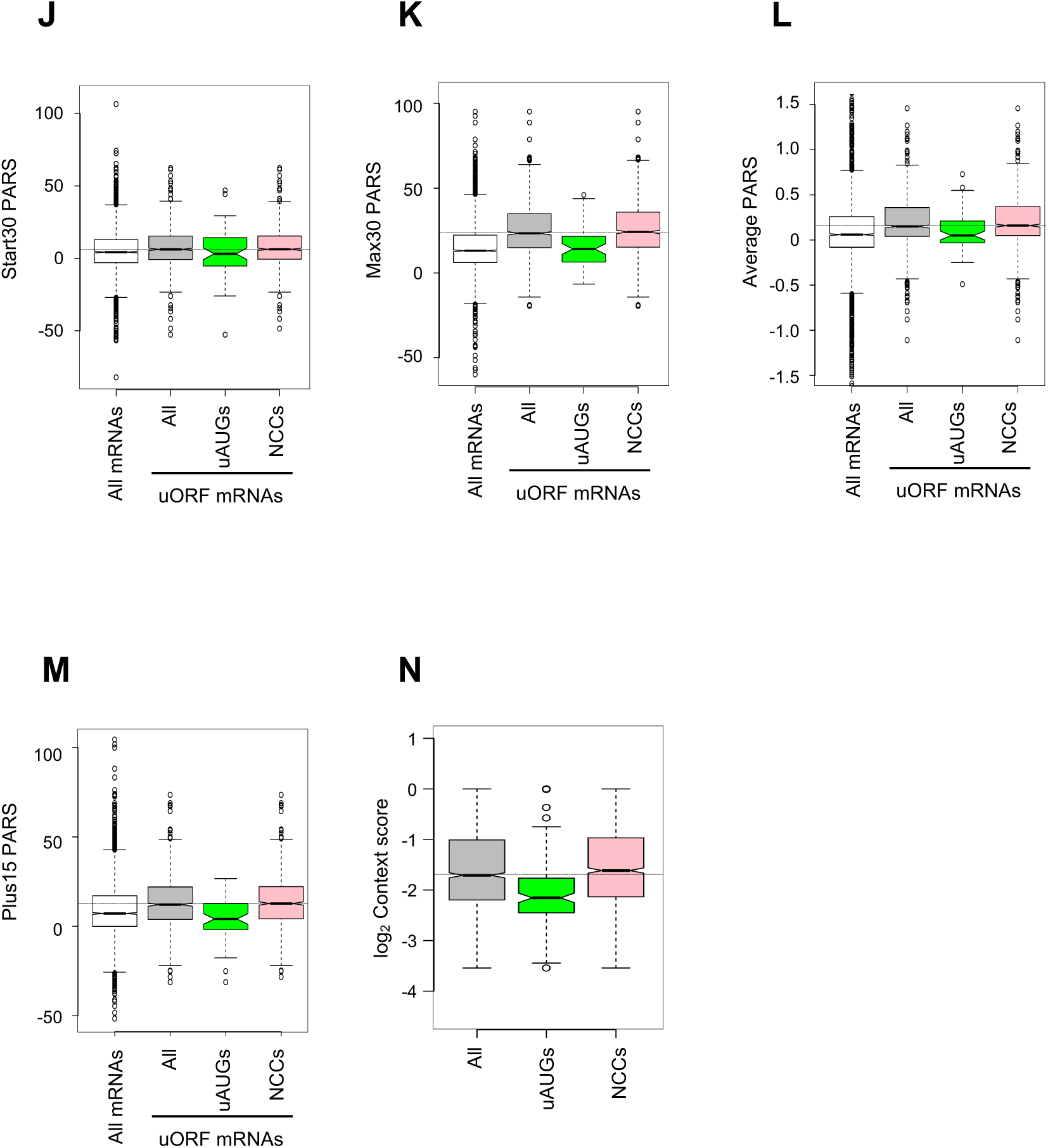
Analysis of the properties of various sets of uORFs and the mRNAs harboring them. **(A-C)** 5’-UTRs of mRNAs with translated uORFs are longer and more structured than the genomic average. **(A)** Cumulative distribution analysis of 5’-UTR lengths for all mRNAs (*All mRNAs* in red), or mRNAs with evidence of at least one translated uORF (NCC or AUG) (*uORF-mRNAs* in purple), or mRNAs that do not have evidence of translated uORFs (*non-uORF mRNAs* in blue). The average 5’-UTR length (in nt) ± SEM, number of mRNAs in each set and p-value by Student’s *t*-test when compared with *All mRNAs* are shown in parenthesis for each set. The maximum difference in cumulative fraction (D) and p-value by Kolmogorov-Smirnov test (when compared with *All mRNAs*) are shown for each set of uORFs on the cumulative distribution plot. To plot the data for *All mRNAs* and *non-uORF mRNAs*, data from previously reported study [1] was interrogated. **(B)** Schematic representation (adapted from [3]) of 5′-UTR intervals (PARS features) used for calculating PARS scores [1]. Orange circle: m^7^G cap; green box: start codon; yellow box: protein coding region; red box: stop codon; black line: mRNA with/without secondary structures. **(C)** Mean PARS scores for each PARS feature calculated for *All mRNAs* (green, N∼2700), *uORF-mRNAs* (purple, N∼1020), *non-uORF mRNAs* (pink, N∼2160) whose PARS scores were available. Error bars indicate standard errors of the mean. P-values of Student’s t-tests for statistical significance of PARS scores for 5’-UTRs between *uORF-mRNAs* and *All mRNAs* or between *non-uORF mRNAs* and *All mRNAs* were calculated. **(D-F)** uORFs were binned with respect to their start sites. *All* represents all the translated uORFs identified in this study as described in Figure 3A (N = 1367); *uAUGs* represents translated AUG uORFs (N = 142) and *NCCs* represents translated NCC uORFs (N = 1225). Boxplot analysis of intrinsic properties viz. (**D)** *Distance from 5’-cap* (distance in nt between the start of the mRNA’s 5’-end and the first base of the start site of the uORF), **(E)** *Distance from mAUG* (distance in nt between the first base of the start site of the uORF and the first base of the downstream mORF), **(F)** *Length of uORF* (distance in nt between first and the last nucleotide of a uORF). **(G-M)** Analysis of 5’-UTR features of uORF mRNAs. **(G)** Boxplots of 5’-UTR lengths [1] for *All mRNAs* (N = 2679); ‘*All uORF mRNAs*’ (all mRNAs with evidence of at least one translated uORF; N = 1367); ‘*AUG uORFs mRNAs*’ (N = 142); and ‘*NCC uORFs mRNAs*’ (N = 1225). **(H-M)** The PARS dataset [1] was interrogated to analyze the propensity of secondary structure formation in and around the 5’-UTRs of the same sets of mRNAs as in (D) for which PARS data are available: *All mRNAs* (n = 2679); ‘*All uORF mRNAs*’ (N = 1024); *AUG uORFs* (N = 59); *NCC uORFs mRNAs* (N = 965). **(N)** Analysis of uORF start codon context scores calculated for positions −3 to −1 and +4 [4] for the same sets of mRNAs as in (D). The dotted horizontal line in each boxplot represents the median value of *All uORFs*.

**Supplementary Figure 7.**
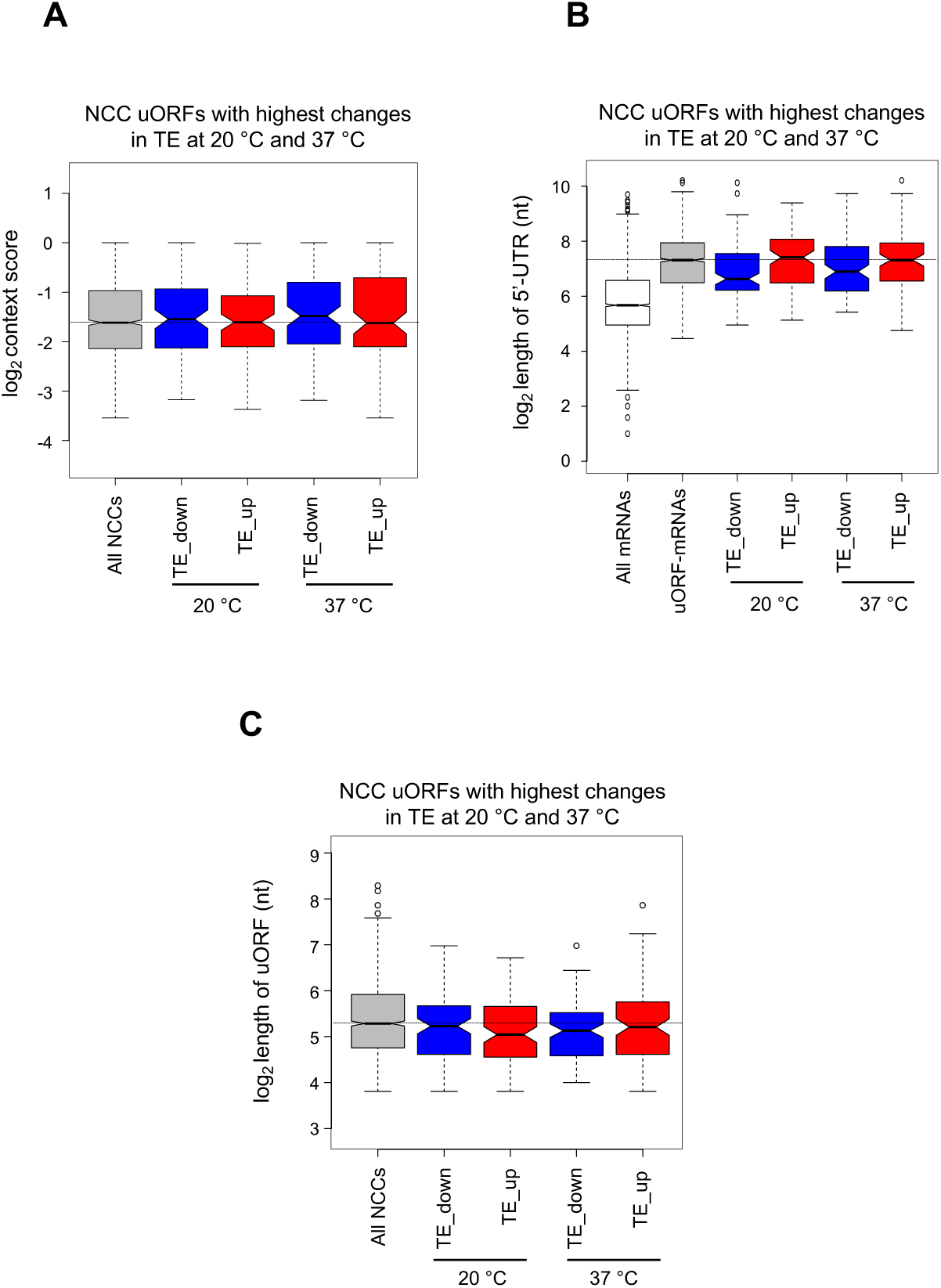
Analysis of uORFs showing highest changes in translation at 20 °C and 37 °C. **A)** Boxplot analysis of uORF start codon context scores for the sets of regulated uORFs described in Figure 7A. *All NCCs* represent all translated NCC uORFs whose context scores were available (N = 1206). The dotted horizontal line represents the median context score for *All NCCs*. For *TE_down* or *TE_up* (20 °C or 37 °C) sets, at least 98/100 uORFs’ context scores were available. **(A)** Boxplot analysis of the lengths of 5’-UTRs of the mRNAs corresponding to various sets of uORFs described in Figure 7A. *All* represents ∼2700 mRNAs whose 5’-UTR length information is available [1]. *uORF-mRNAs* represents mRNAs with NCC uORFs (N = 1223) whose ΔTE_uORF_ could be determined and for which 5’-UTR length information was available. The dotted horizontal line represents the median length of the 5’-UTRs of *uORF-mRNAs*. **(B)** Boxplot analysis of the lengths of uORFs (in nucleotides) in different datasets described in Figure 7A. *All* represents all the uORFs identified in this study starting with NCC and whose ΔTE_uORF_ could be measured (N = 1223). The dotted horizontal line represents the median length of *All NCCs*.

**Supplementary Figure 8.**
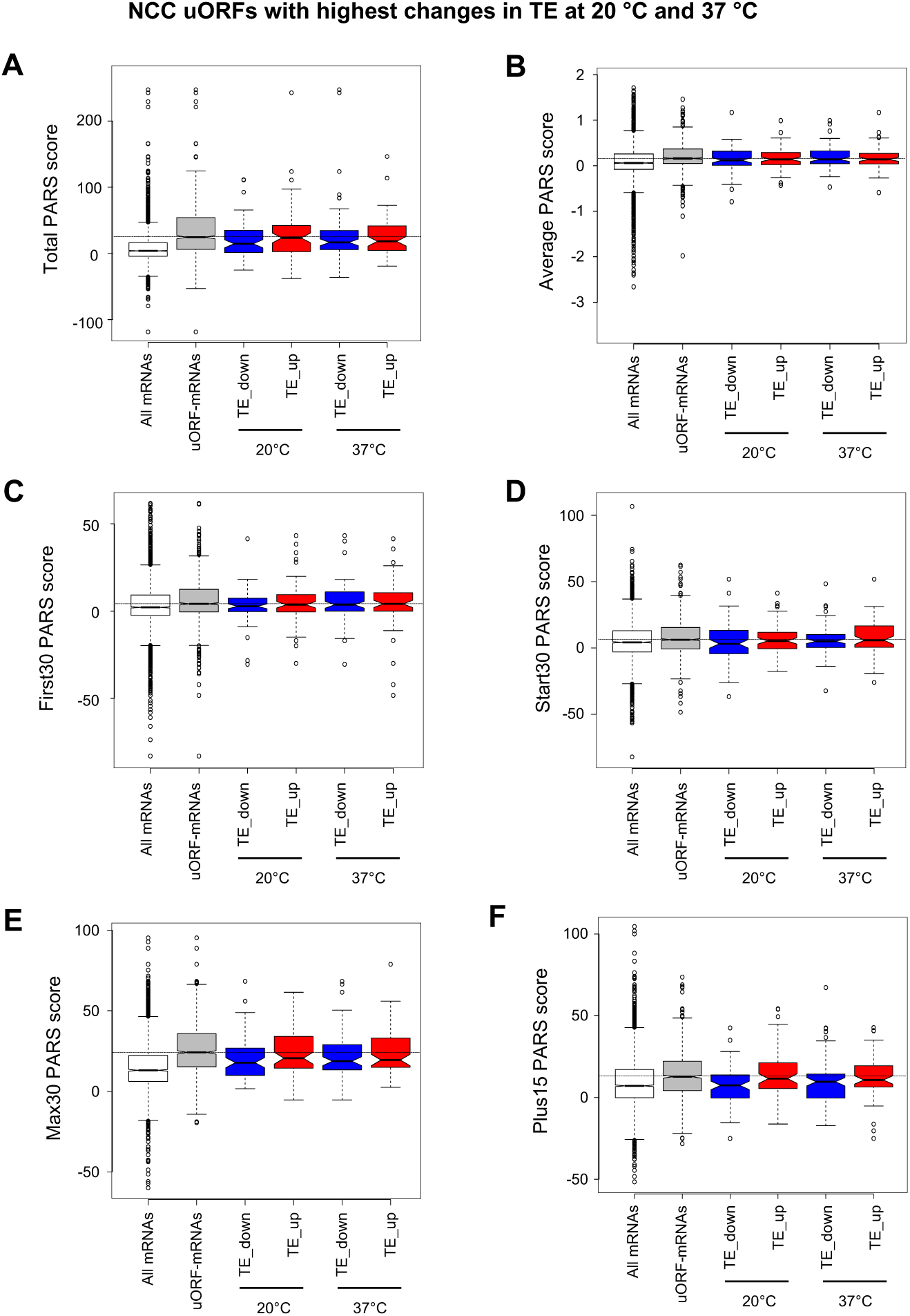
Boxplot analyses of 5’-UTR features for the most highly regulated NCC uORFs. **(A-F)** Boxplot analysis of PARS scores of the 5’-UTRs of TE_up and TE_down NCC uORF mRNAs (described in Figure 7A). *All mRNAs* represent data from 2679 mRNAs whose PARS scores are available [1]. *uORF-mRNAs* represent mRNAs with evidence of at least one translated NCC uORF identified using the pipeline described in Figure 3A (N = 1223) and whose PARS data are available (N = 965/1223). *TE_down* and *TE_up* (at either 20 °C or 37 °C) represent the most highly regulated uORFs as described in Figure 7A. PARS data were available for at least 65/100 uORFs in each category. The dotted horizontal line represents the median PARS score for each feature calculated for the 5’-UTRs of *uORF-mRNAs*. **(A)** Analysis for total PARS score. **(B)** Analysis for average PARS score. **(C)** Analysis for First30 PARS score. **(D)** Analysis for Start30 PARS score. **(E)** Analysis for Max30 PARS score. **(F)** Analysis for Plus15 PARS score. PARS features are as defined in Additional file 1: Figure S6B.

**Supplementary Figure 9.**
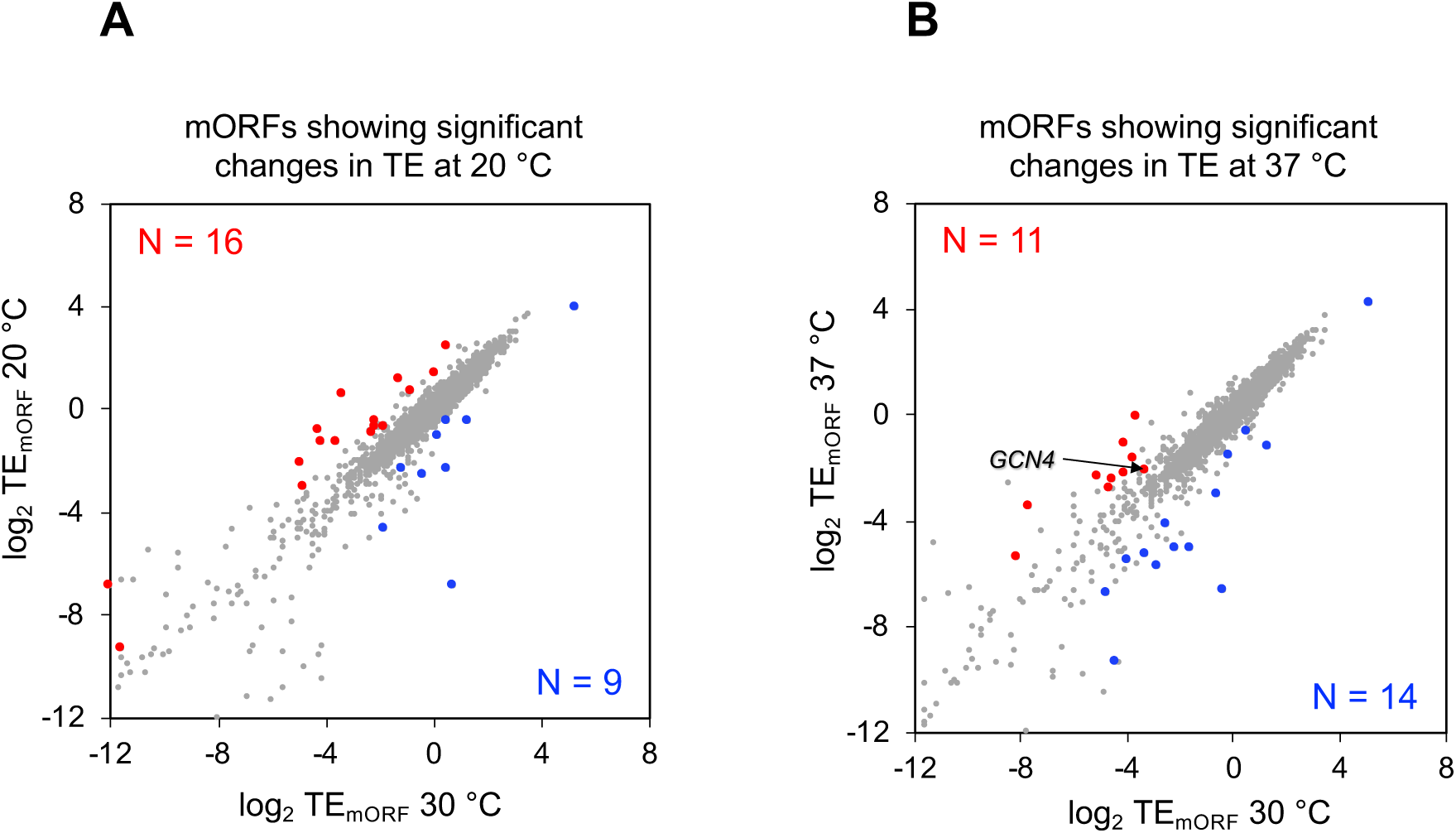
Analysis of TE-changes of mORFs. **(A)** Scatterplot of translational efficiencies (TEs) of mORFs in cells cultured at 30 °C and 20 °C. mORFs exhibiting ≥ 2-fold changes in TE in cells cultured at 20 °C and at FDR < 0.1 are shown in blue circles (repressed) or in red circles (activated). **(B)** Same as in **A**, except translational efficiencies (TEs) of mORFs of cells cultured at 30 °C and 37 °C is plotted.

**Supplementary Figure 10.**
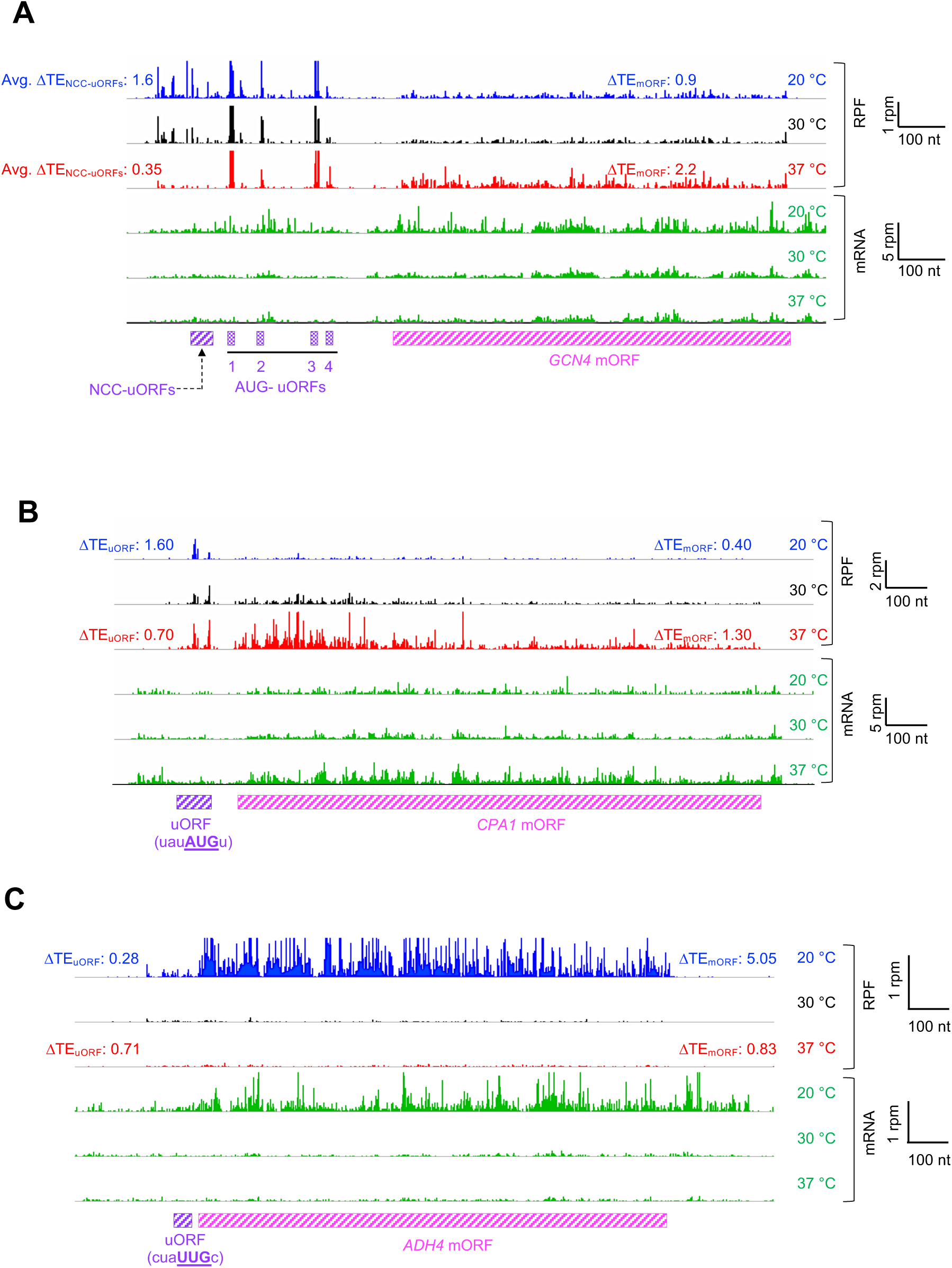
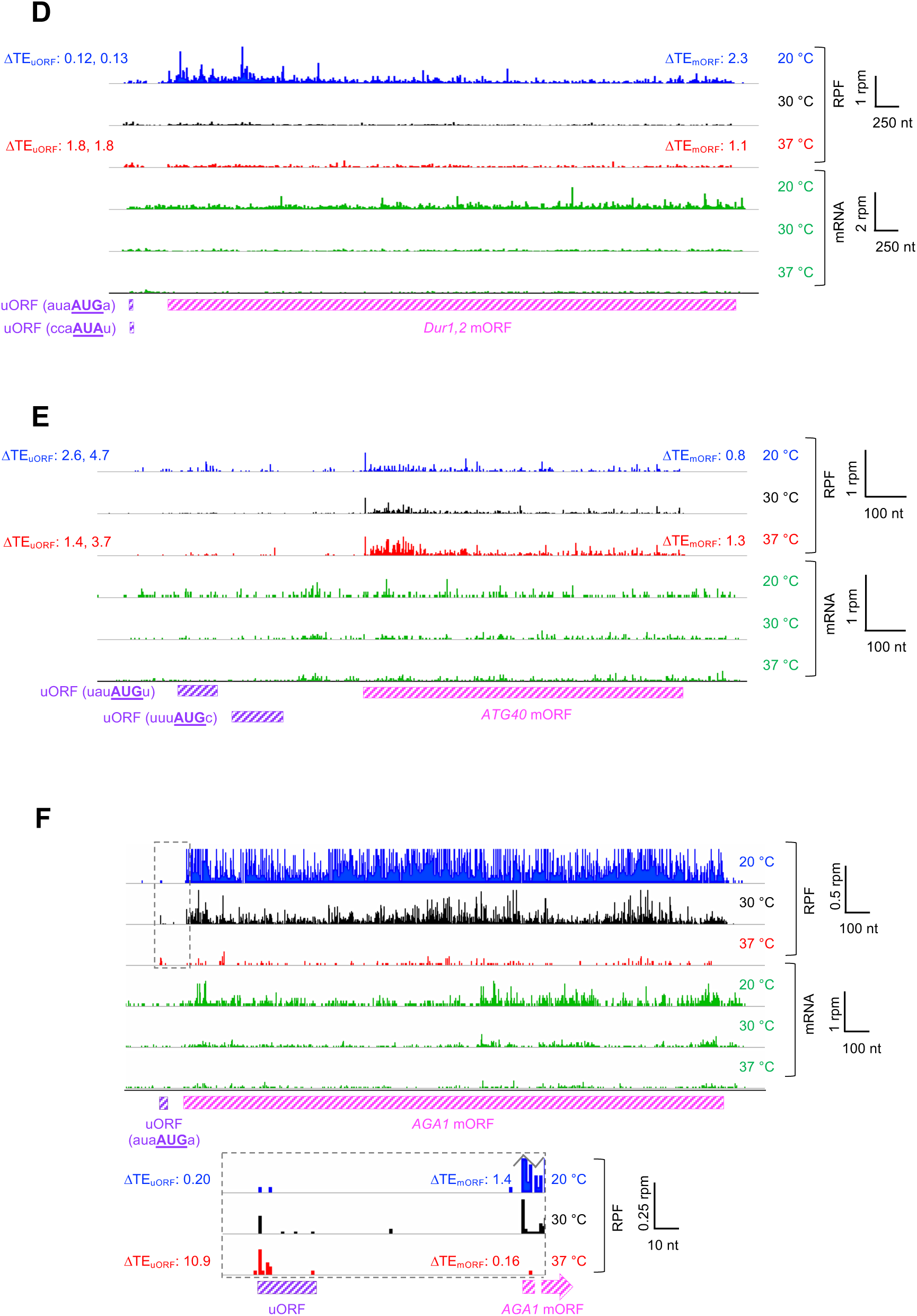
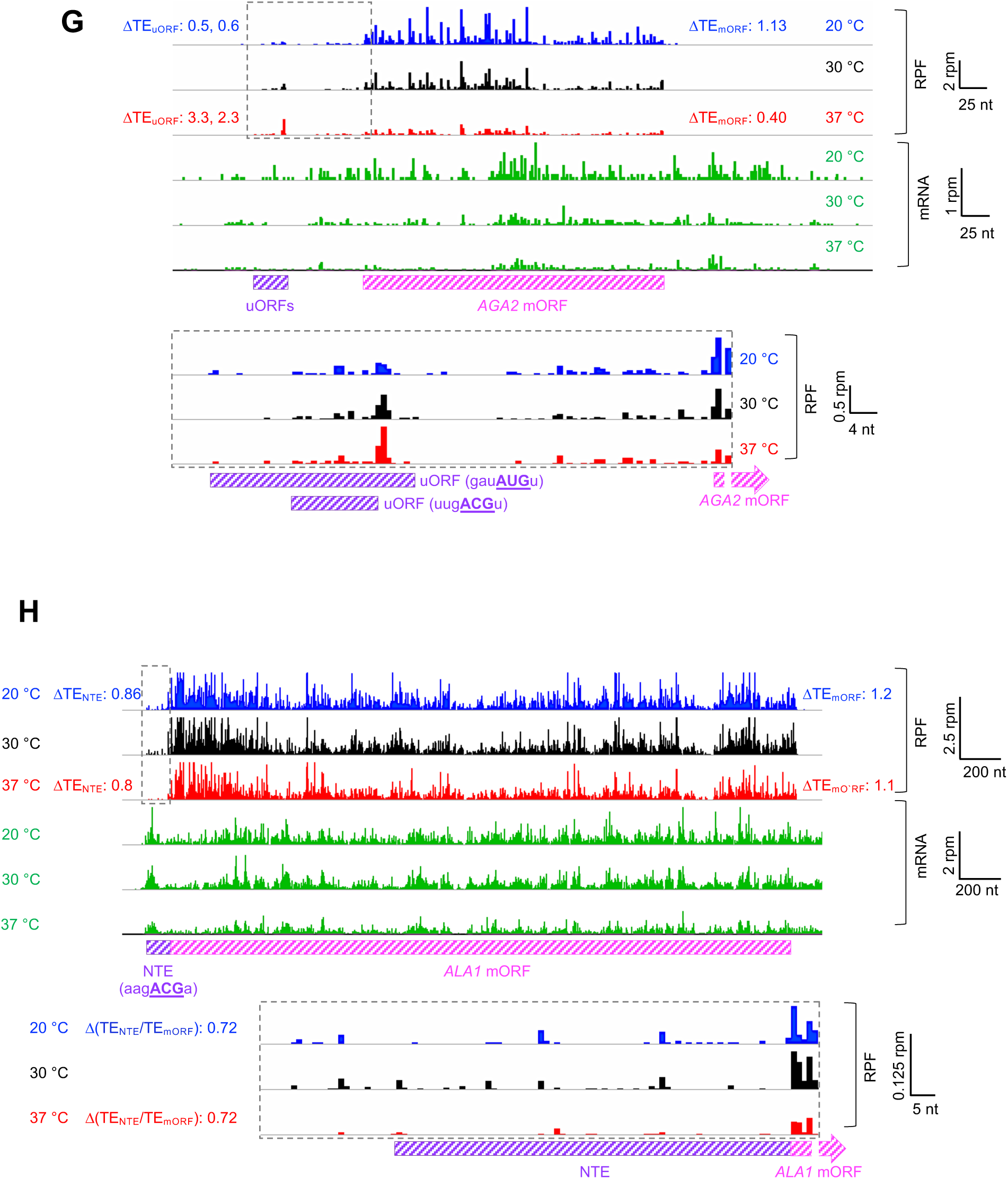
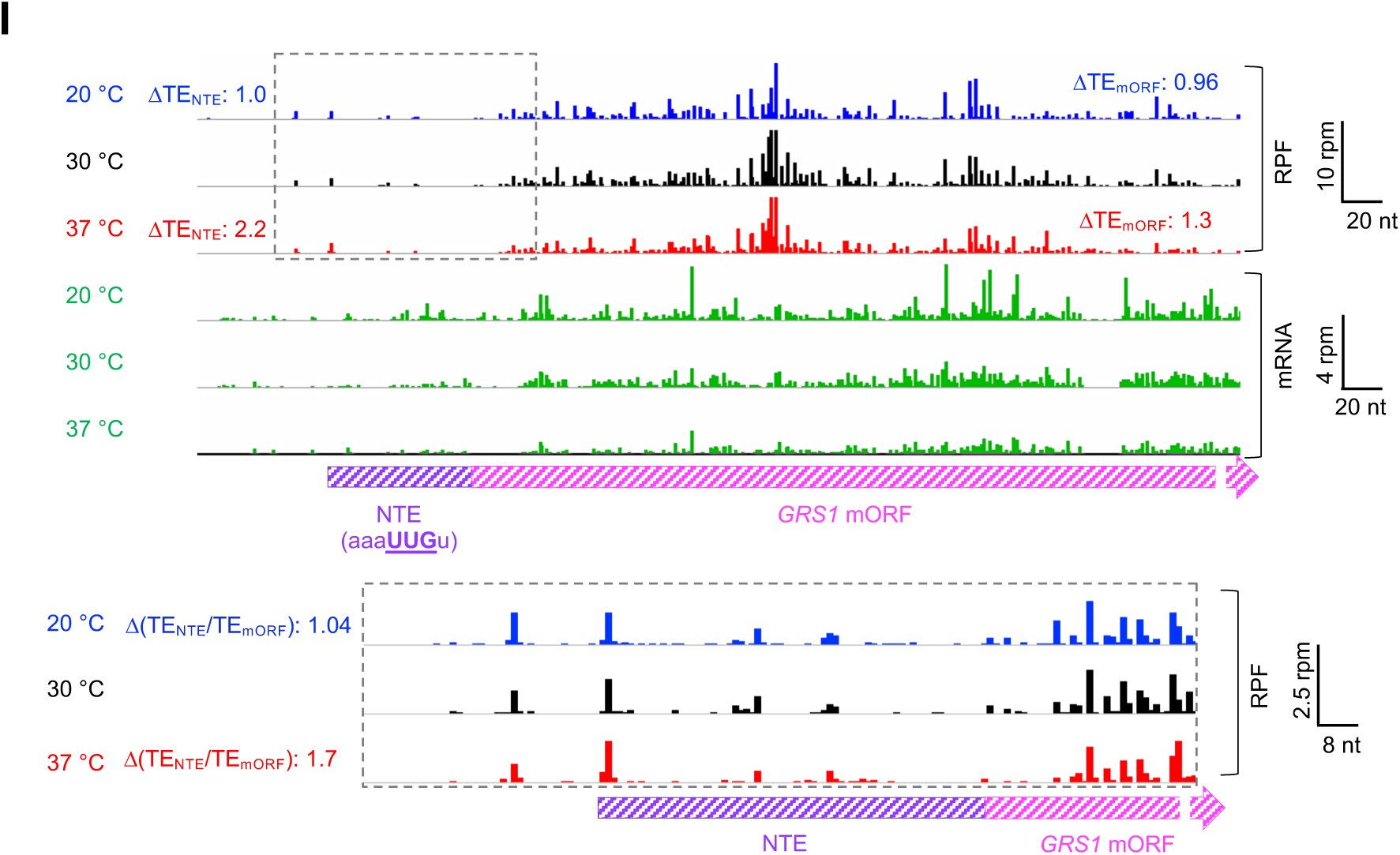
Wiggle track images of examples of mRNAs that show temperature-dependent changes in translation of one or more uORF and the mORF. (**A-I**) Ribosome-protected fragments (RPFs) and mRNA reads are shown for data from cells cultured at either 20, 30 or 37 °C, in units of rpm (reads per million mapped reads) as in Figure 9A-I except the RPF-tracks were <u>not</u> normalized to the mRNA levels and the traces of mRNA reads from each temperature are shown in green.

## Additional file 2: Supplementary Tables 1 to 4

**Supplementary Table 1.** Yeast strains used in this study.

| Sr. No. | Strain | Genotype | Source/reference |
| --- | --- | --- | --- |
| 1 | BY4741 | *MATa his3Δ1 leu2Δ0 met15Δ0 ura3Δ0* | Open Biosystems |
| 2 | *upf1Δ* | *MATa his3 leu2 met15 ura3 upf1:KanMX6* | [1] |
| 3 | BY4743 | *MATa /a his3Δ/his3Δ1 leu2Δ0/leu2Δ0 LYS2/lys2Δ0 met15Δ0/MET15 ura3Δ0/ura3Δ0* | Open Biosystems |
| 4 | YNL244C (+/*sui1Δ*) | *MATa/a his3Δ/his3Δ1 leu2Δ0/leu2Δ0 LYS2/lys2Δ0 met15Δ0/MET15 ura3Δ0/ura3Δ0+/*sui1Δ** | Open Biosystems |
| 5 | YMR260C (+/*tif11Δ*) | *MATa/a his3Δ/his3Δ1 leu2Δ0/leu2Δ0 LYS2/lys2Δ0 met15Δ0/MET15 ura3Δ0/ura3Δ0+/*tif11Δ** | Open Biosystems |
| 6 | YPR041W (+/*tif5Δ*) | *MATa/a his3Δ/his3Δ1 leu2Δ0/leu2Δ0 LYS2/lys2Δ0 met15Δ0/MET15 ura3Δ0/ura3Δ0+/*tif5Δ** | Open Biosystems |

**Supplementary Table 2.** Plasmids used in this study.

| **Sr.**<br>**No.** | **Plasmid** | **Genotype** | **Source/reference** |
| --- | --- | --- | --- |
| 1 | pR<sup>AUG</sup>FF<sup>AUG</sup> | Dual luciferase reporter R-Luc(AUG)-F-Luc(AUG) in *URA3* vector | [2] |
| 2 | pR<sup>AUG</sup>FF<sup>UUG</sup> | Dual luciferase reporter R-Luc(AUG)-F-Luc(UUG) in *URA3* vector | [2] |
| 3 | pR<sup>AUG</sup>FF<sup>CUG</sup> | Dual luciferase reporter R-Luc(AUG)-F-Luc(CUG) in *URA3* vector | [2] |
| 4 | pR<sup>AUG</sup>FF<sup>GUG</sup> | Dual luciferase reporter R-Luc(AUG)-F-Luc(GUG) in *URA3* vector | [2] |
| 5 | pR<sup>AUG</sup>FF<sup>ACG</sup> | Dual luciferase reporter R-Luc(AUG)-F-Luc(ACG) in *URA3* vector | [2] |
| 6 | pR<sup>AUG</sup>FF<sup>AUC</sup> | Dual luciferase reporter R-Luc(AUG)-F-Luc(AUC) in *URA3* vector | [2] |
| 7 | pR<sup>AUG</sup>FF<sup>AUU</sup> | Dual luciferase reporter R-Luc(AUG)-F-Luc(AUU) in *URA3* vector | [2] |
| 8 | pR<sup>AUG</sup>FF<sup>AUA</sup> | Dual luciferase reporter R-Luc(AUG)-F-Luc(AUA) in *URA3* vector | [2] |
| 9 | p367 | sc *URA3* HIS4(AUG)-lacZ | [3] |
| 10 | p391 | sc *URA3* HIS4(UUG)-lacZ | [3] |
| 11 | hc-*SUI1* | *SUI1* on a high-copy plasmid (*LEU2* vector) | [4] |

**Supplementary Table 3.**
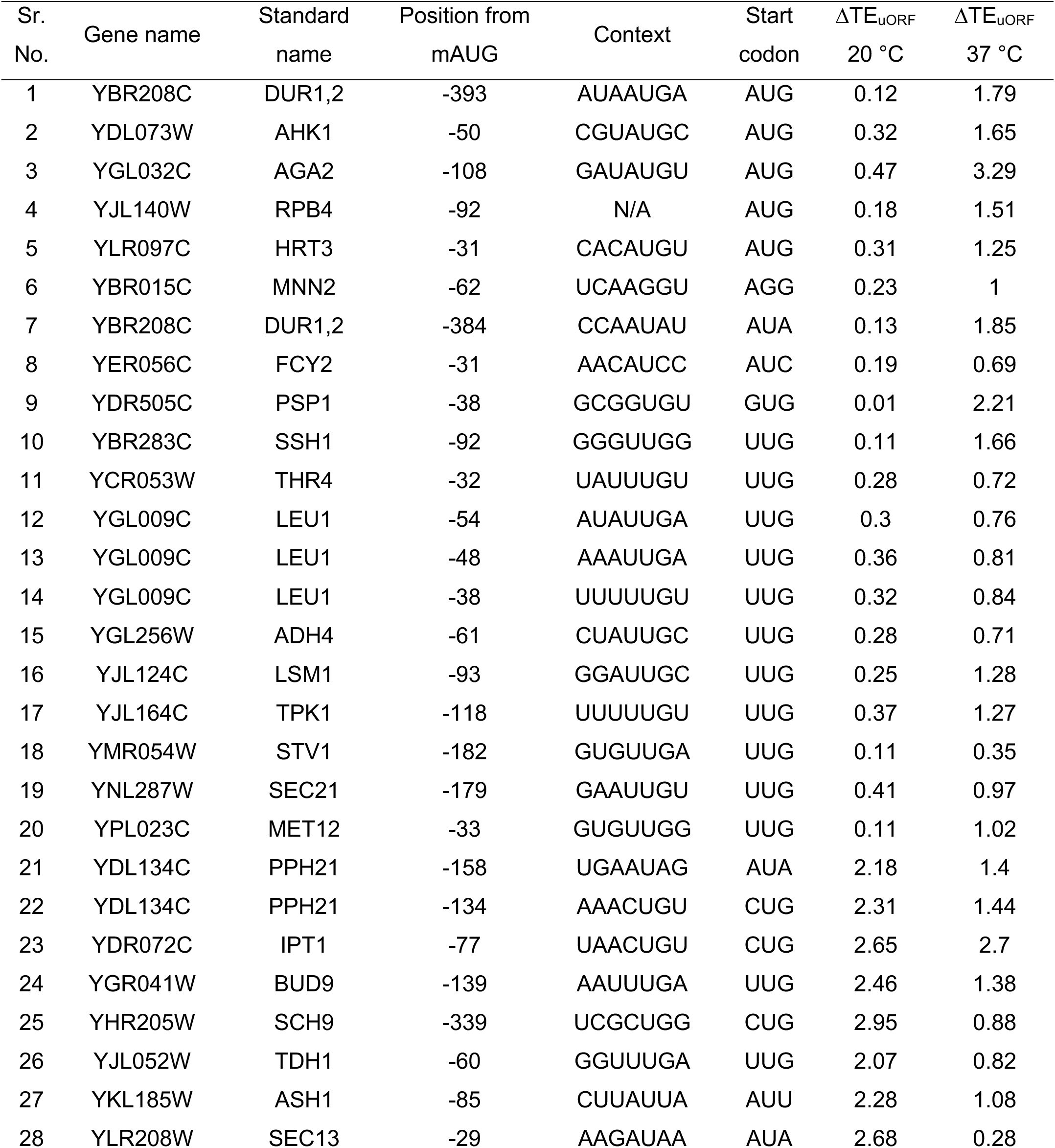
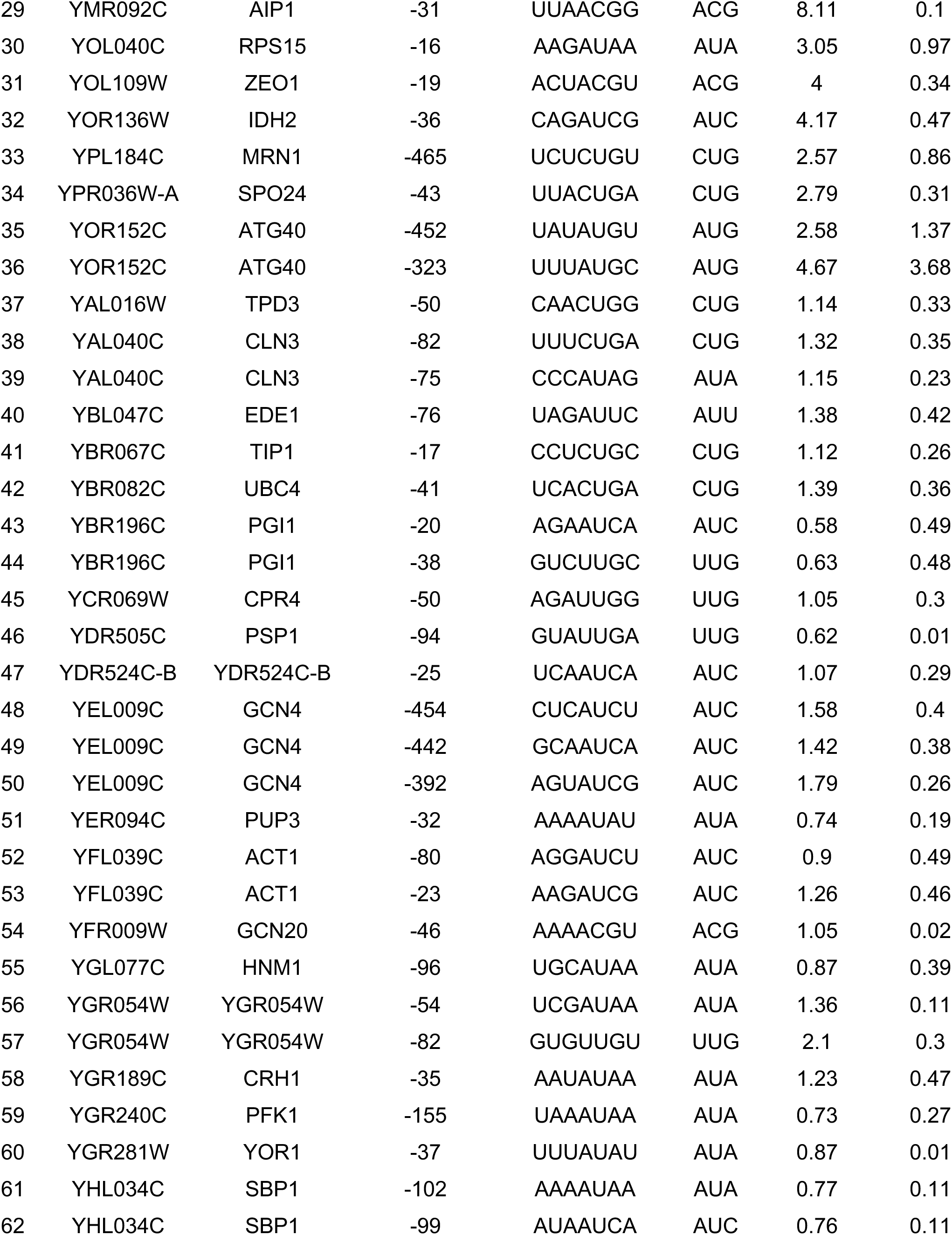
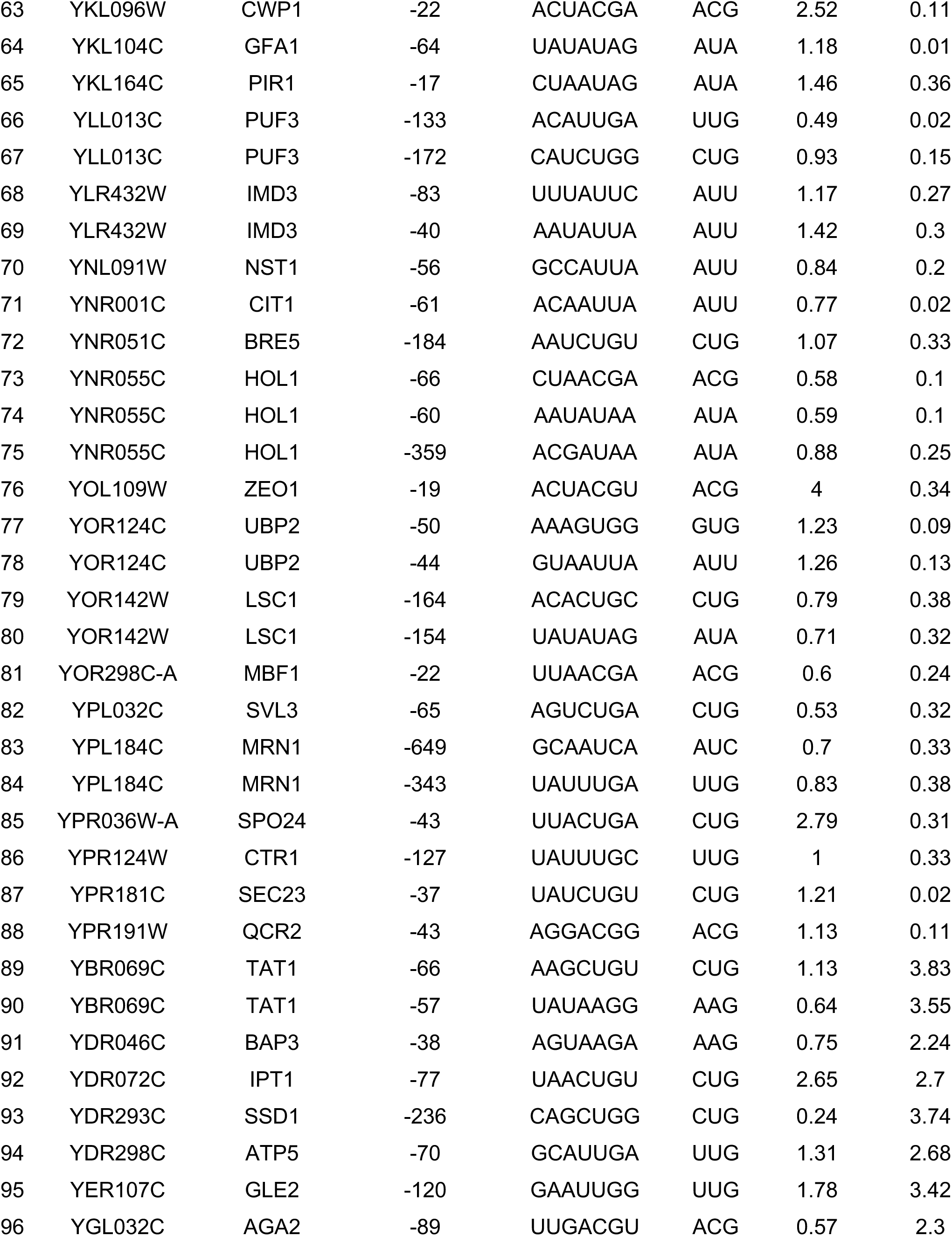
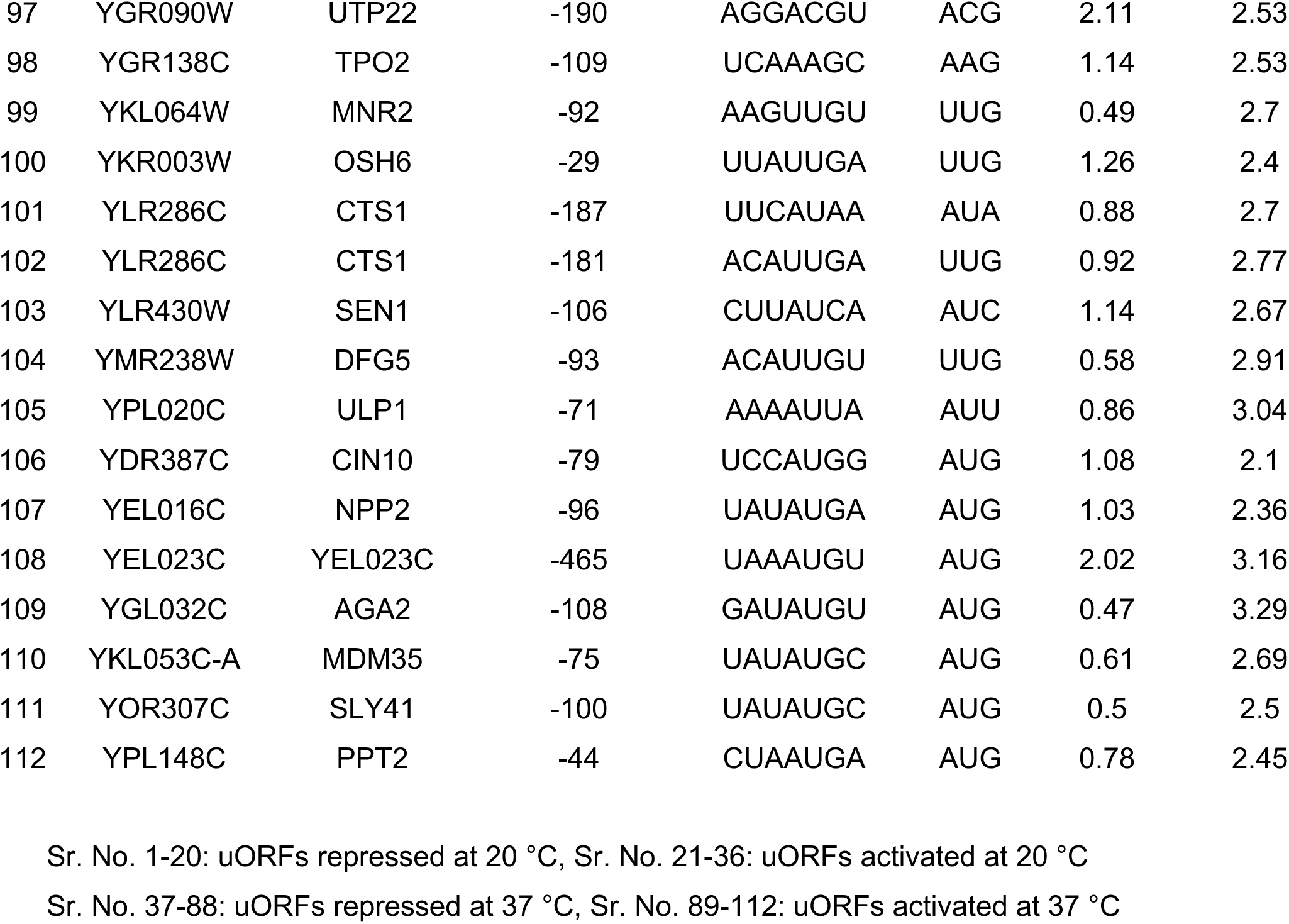
List of uORFs showing temperature-dependent translational regulation.

**Supplementary Table 4.**
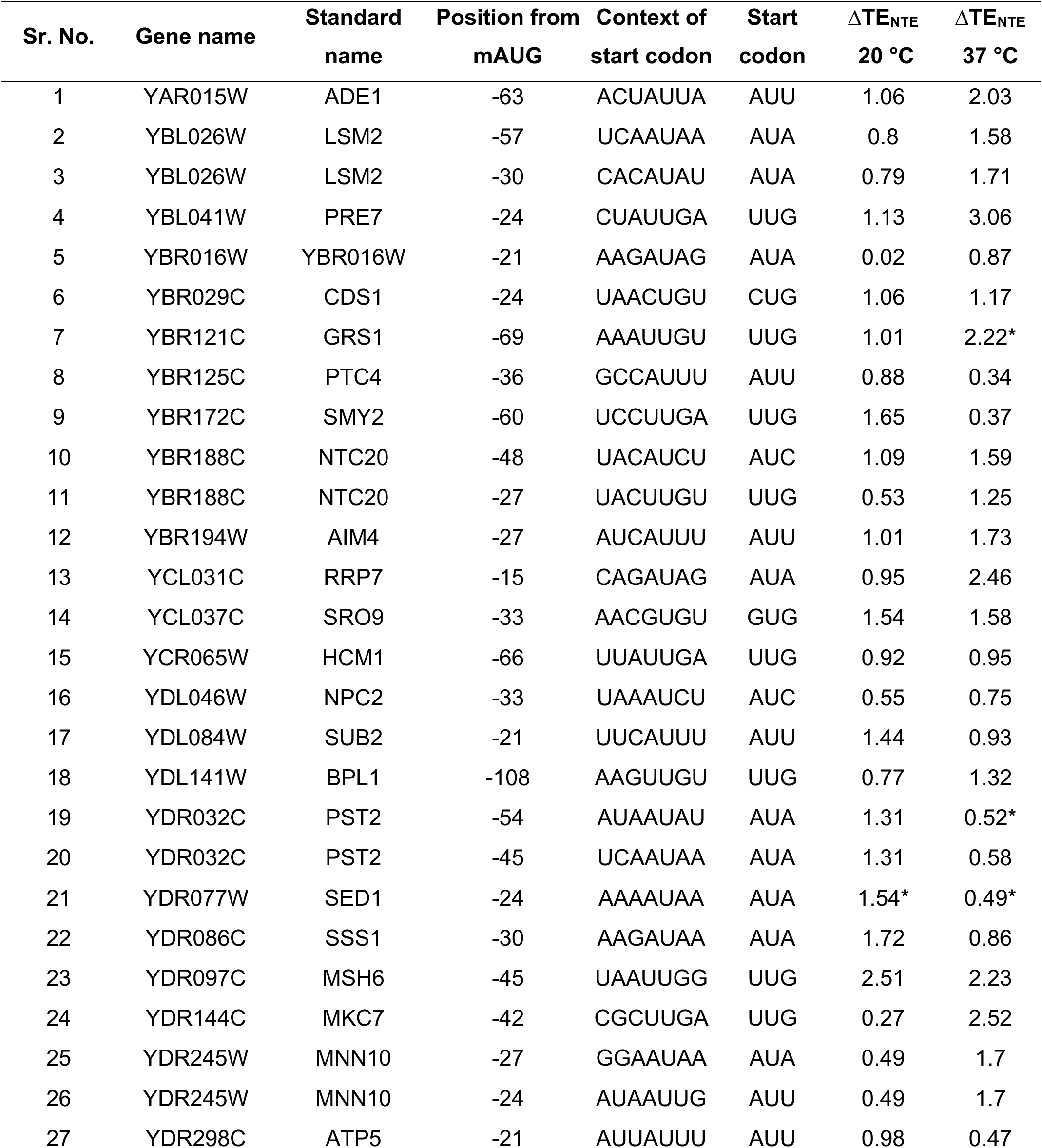
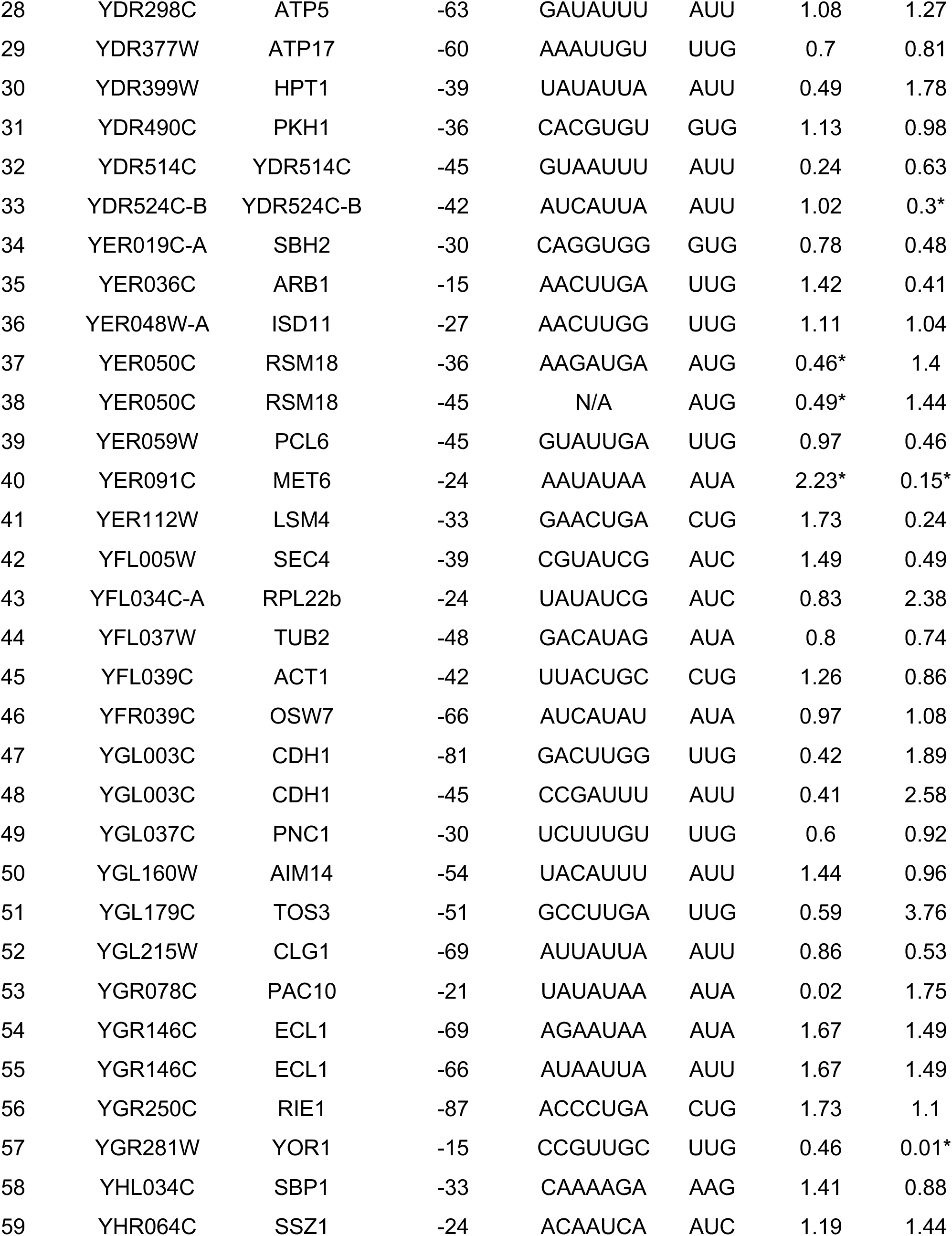
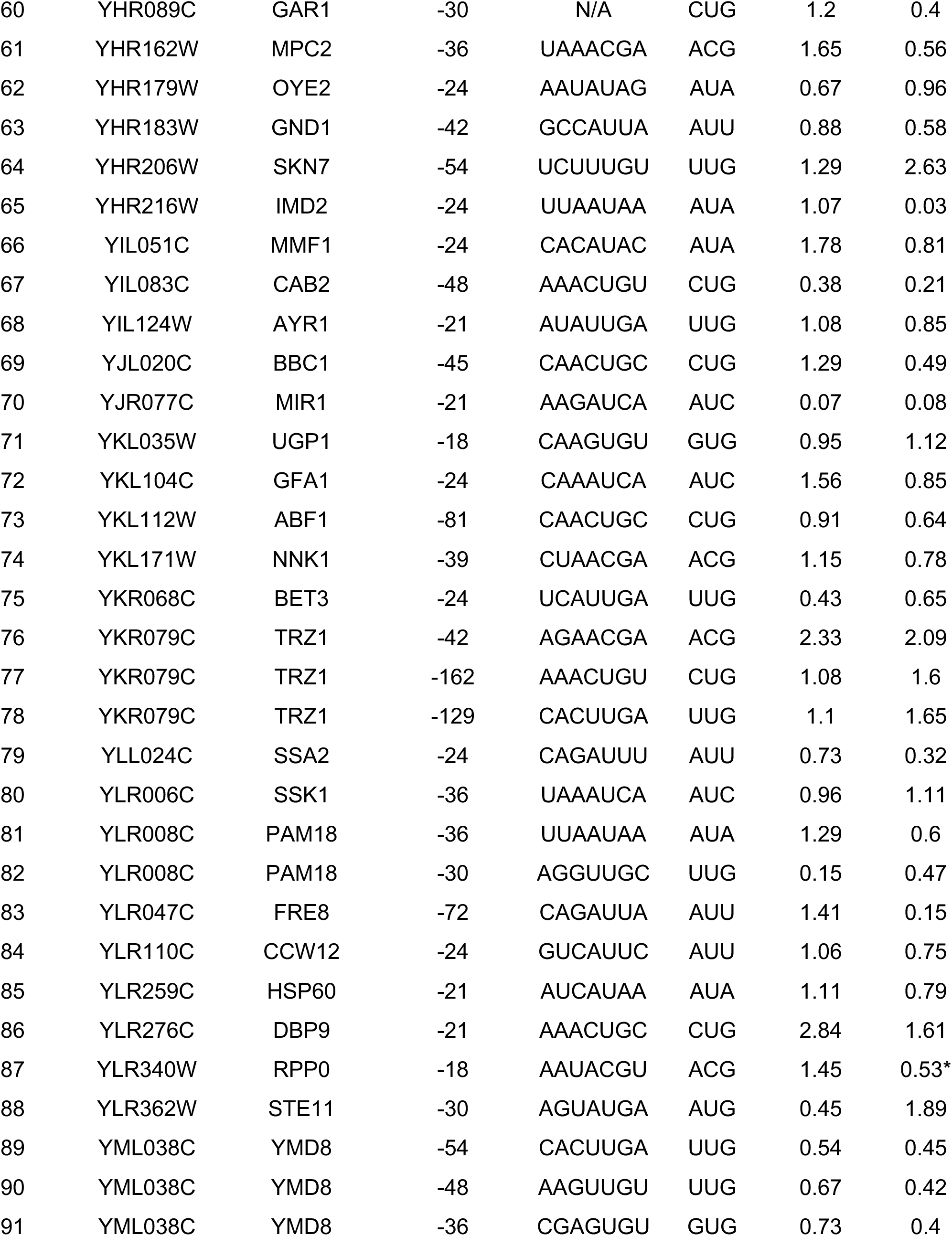
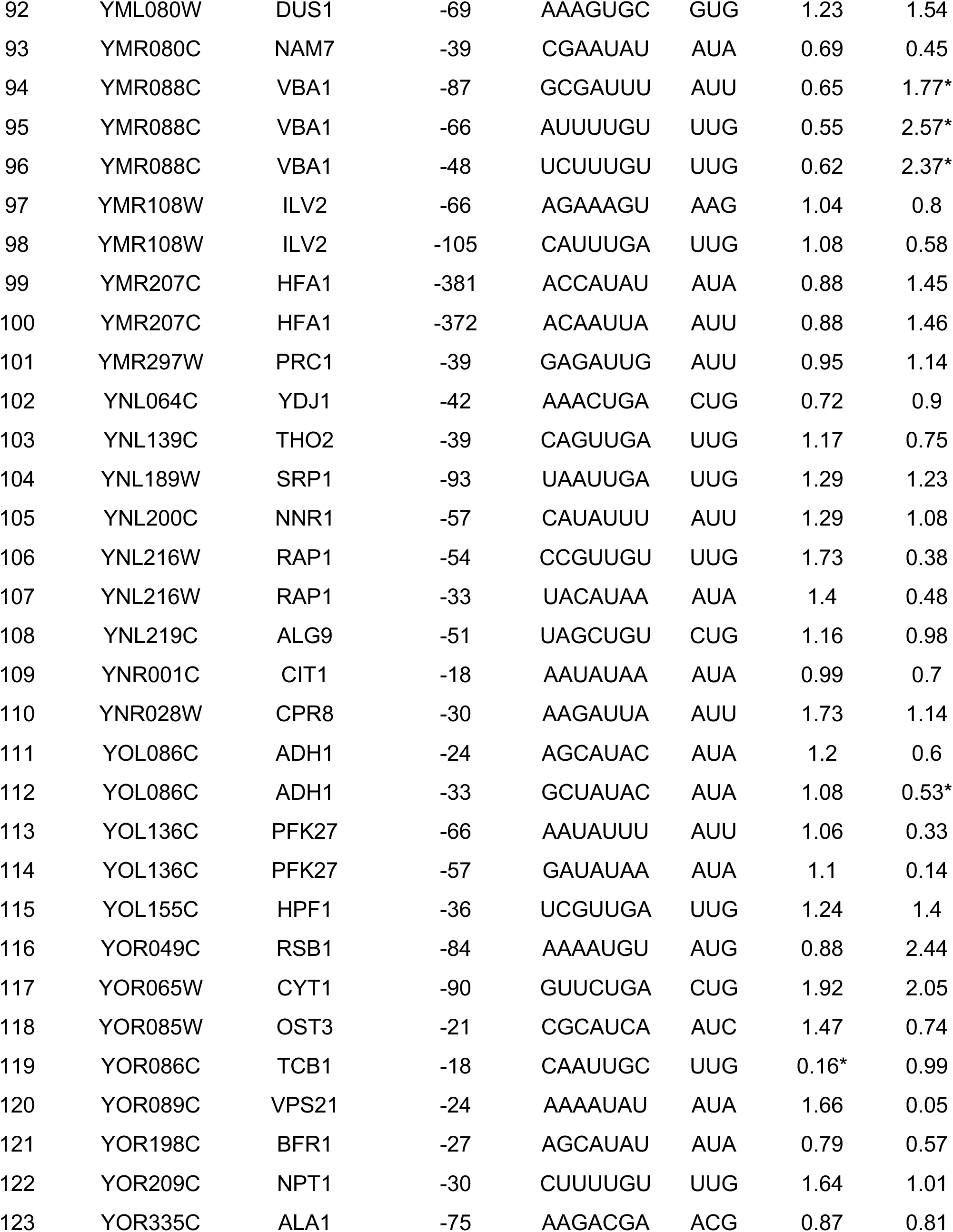
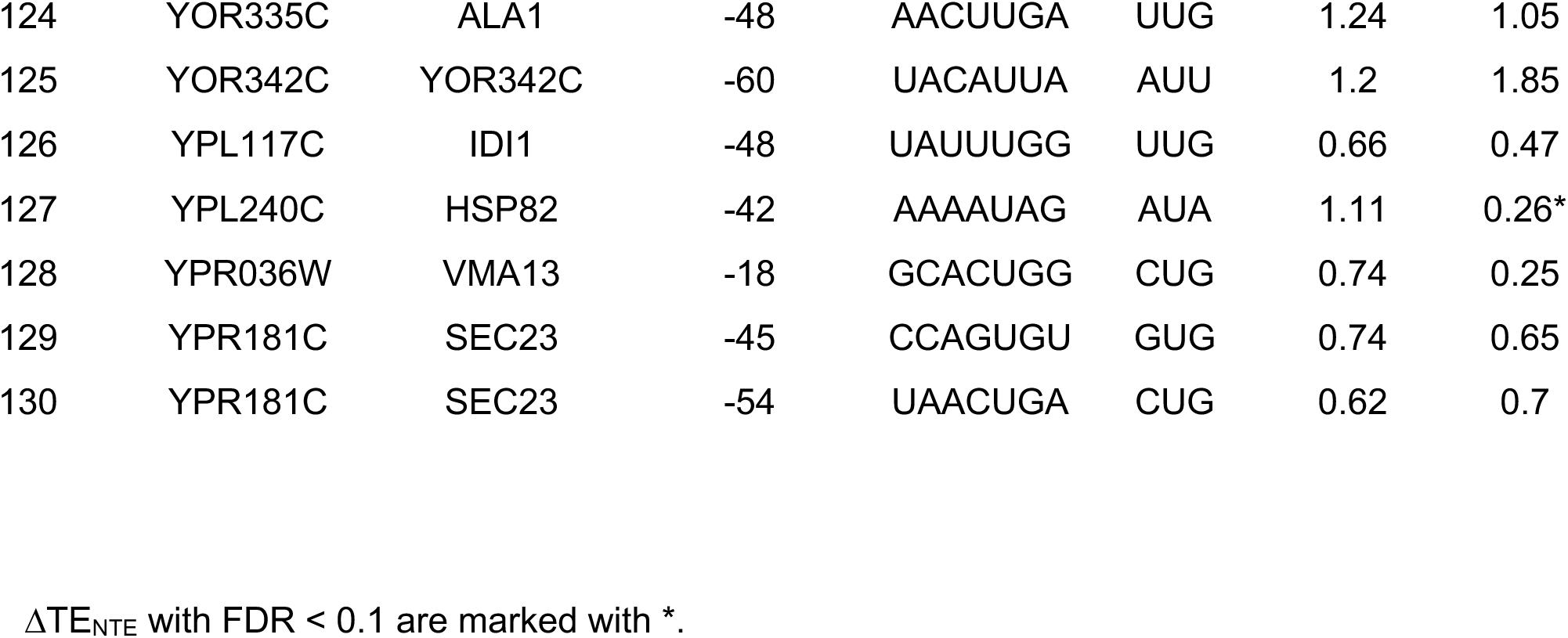
List of putative N-terminal extensions identified in this study.

